# Wildfire, restoration, and post-wildfire rehabilitation effects on wind erosion in the Great Basin

**DOI:** 10.64898/2026.01.16.699976

**Authors:** Ronald S. Treminio, Nicholas P. Webb, Brandon L. Edwards, Beth A. Newingham, Magda Garbowski, Colby Brungard, David Dubois, Akasha Faist, Emily Kachergis, Carrie-Ann Houdeshell

## Abstract

Restoration of degraded areas and post-disturbance rehabilitation after wildfire encompass critical approaches for reducing and reversing impacts of wind erosion and sand and dust storms (SDS). However, the broad outcomes of dryland restoration and rehabilitation for wind erosion and SDS remain underexplored. Wind erosion is an emerging issue in the Great Basin of the western United States, exacerbated by invasive annual grasses and associated wildfire. Here, we assess potential wind erosion and SDS responses to wildfire, restoration, and post-wildfire rehabilitation treatments at the regional scale in the Great Basin. We used 13 years of rangeland monitoring data, the Aeolian EROsion model, and the Land Treatment Digital Library to produce counterfactual model-predictions to estimate treatment effects. Our results revealed reductions in aeolian sediment fluxes (Ln *Q* < 0 g m^-1^ d^-1^) across wildfire-affected regions (mean ± SE: −0.070 ± 0.077 Ln *Q*), restoration treatments in unburned areas (range: −0.867 ± 0.398 to 0.480 ± 0.253 Ln *Q*), and post-wildfire rehabilitation (range: −0.821 ± 0.183 to 1.278 ± 0.909 Ln *Q*). In particular, aerial seeding and soil disturbance restoration treatments, and post-wildfire closure-treatments had higher perennial grass cover and the most decreased Ln *Q* compared to untreated controls. These results represent an important regional scale assessment of wind erosion responses to restoration and post-wildfire rehabilitation. Our findings underscore the application of integrating wind erosion and SDS mitigation into restoration and post-disturbance rehabilitation programs to provide land managers with strategies to reduce land degradation while fostering ecosystem resilience.

## 1. Introduction

Wind erosion and sand and dust storms (SDS) are symptoms and drivers of land degradation that impact agriculture, ecosystems, and societies globally. Policy initiatives and practical guidance for mitigating wind erosion and SDS, including the United Nations Convention to Combat Desertification (UNCCD) Sand and Dust Storms Coalition (UNCCD, 2022), World Overview of Conservation Approaches and Technologies (WOCAT) Sustainable Land Management Database (WOCAT, 2025), and USDA Dust Mitigation Handbook (Smarik et al., 2019), support the challenge of avoiding, reducing, and reversing wind erosion and SDS impacts. However, while integrating wind erosion and SDS mitigation into restoration and rehabilitation programs to address land degradation is increasingly recognized as important (UNCCD Decision 27/COP.16, 2024), studies linking land management to wind erosion and SDS remain limited to certain land uses and practices.

Research on wind erosion and SDS mitigation has largely focused on impact reduction for cropland (e.g., Jarrah et al., 2020; Pi et al., 2020), transportation, and energy extraction infrastructure (Middleton and Al-Hemoud 2024; Duniway et al. 2019). Rangelands have received comparatively less attention (van Oudenhoven et al., 2015; Webb et al., 2017). The dearth of rangeland wind erosion and SDS mitigation research contrasts with studies, “on-the-ground” application, and formal programs that support rangeland restoration and post-disturbance rehabilitation (e.g., Allred et al., 2021; Bestelmeyer, 2006; Eba Tebeje et al., 2014; Mganga et al., 2015). While dryland restoration and rehabilitation objectives often encompass soil stabilization and returning ecosystem functions to a more desirable condition (Copeland et al., 2018; Ma et al., 2017; Pilliod et al., 2021; Shriver et al., 2019), explicitly examining treatment outcomes and co-benefits for wind erosion and SDS mitigation provides an opportunity for direct understanding of these relationships (e.g., Miller et al., 2012; Morra et al., 2024; Pyke et al., 2013; van Oudenhoven et al., 2015). As wind erosion and SDS hotspots are typically dispersed across landscapes, understanding the regional-scale outcomes of restoration and rehabilitation is essential for identifying effective practices and for informing policy and programs to combat wind erosion and SDS (Ma et al., 2017; Mganga et al., 2015; van Oudenhoven et al., 2015). Incorporating wind erosion and SDS into assessments of restoration and rehabilitation treatment outcomes could yield insights into treatment practices that effectively enhance mitigation strategies that also reduce wind erosion risk.

In the Great Basin of the western United States, wind erosion and dust storms are emerging issues due to a myriad of interacting factors (Treminio et al., 2024). Exotic annual grasses, such as cheatgrass (*Bromus tectorum*), are expanding in range (Boyte and Wylie, 2016; Pastick et al., 2020) and have contributed to more frequent and severe wildfires (Balch et al., 2013), loss of historical plant communities (Chambers et al., 2016; Ellsworth et al., 2020; Svejcar et al., 2017), and increased post-fire aeolian sediment transport (Morra et al., 2024; Sankey et al., 2009a, 2009b; Treminio et al., 2024; Wagenbrenner et al., 2013). Agricultural practices (Chambers and Wisdom, 2009), overgrazing (Field et al., 2011), urbanization (Chambers and Wisdom, 2009), and mining (Morris and Rowe, 2014) have further increased wind erosion risk. Extensive restoration and post-fire rehabilitation treatments have been implemented throughout the Great Basin, with a focus on stabilizing soils, controlling invasive species, improving wildlife habitat, and enhancing forage quality and quantity for livestock (Pilliod et al., 2017). However, while many studies have assessed treatment outcomes for plant communities (Peppin et al., 2010) and wildlife (e.g., Knutson et al., 2014; Simler-Williamson and Germino, 2022), overall treatment effects on wind erosion and SDS risk across the Great Basin remain unquantified.

Assessing rangeland treatment outcomes is inherently challenging. Efforts to assess the gains to ecosystem functioning resulting from restoration and post-disturbance rehabilitation have been inconsistent and insufficiently addressed due to large gaps in application of methodology, standardized assessment criteria, and comprehensive long-term monitoring (e.g., Carlucci et al., 2020; Kollmann et al., 2016; Prach et al., 2019). Randomized-controlled studies, which may be used to elucidate causal treatment effects, are often constrained by small sample sizes and logistical limitations, making them unsuitable for assessing regional restoration and rehabilitation outcomes. Consequently, observational studies are common. An emerging approach to elucidate causal treatment effects from observational studies is counterfactual analysis, which uses synthetic controls based on sites with similar ecological potential to test treatment outcomes using remote sensing (e.g., Fick et al., 2021; McNellis et al., 2023; Nauman et al., 2017). Recently, propensity score matching (PSM), has been used with field monitoring and remotely sensed data to address the limitations of observational studies by simulating the balanced treatment assignment seen in randomly controlled studies (e.g., Kluender et al., 2024; Simler-Williamson and Germino, 2022). In the US, the growing availability of rangeland treatment data organized in the Land Treatment Digital Library (LTDL, Pilliod et al., 2019; Pilliod and Welty, 2013), and collection of standardized monitoring data, such as the Bureau of Land Management’s (BLM) Assessment, Inventory and Monitoring (AIM) program data (Toevs et al., 2011) that can be input to the Aeolian EROsion (AERO) model (Edwards et al., 2022), present new opportunities for using counterfactual analytical approaches to assess treatment outcomes across scales incorporating wind erosion and SDS risk.

Here, we assess potential wind erosion and SDS responses to rangeland restoration and post-wildfire rehabilitation treatments across the Great Basin at the regional scale. Specifically, we address three questions: 1) What is the magnitude and direction of aeolian sediment transport responses to wildfire, rangeland restoration in the absence of wildfire, and post-wildfire rehabilitation treatments? 2) How does the magnitude of aeolian sediment transport by treatment vary compared to untreated reference plots? and 3) How do responses of plant functional-group cover to treatments influence potential aeolian sediment transport rates? We hypothesized that aeolian sediment transport rates, an indicator of wind erosion and SDS risk, will differ by treatment type, wildfire, and their interaction depending on vegetative cover responses. Applying the AERO model to AIM monitoring plots sampled (2011-2023) across treatments represented in the LTDL, we used PSM and statistical methods to elucidate rangeland restoration and post-wildfire rehabilitation effects on potential wind erosion. Here, due to sampling constraints, we focus on wind erosion responses to the most recent wildfires and treatments, recognizing that many areas of the Great Basin have experienced multiple wildfires and treatments over the last century.

## 2. Methods

### 2.1 Data

The United States Geological Survey’s Land Treatment Digital Library (LTDL) is a database containing information on known rangeland treatments conducted on US public lands managed by the BLM (Pilliod et al., 2019). These treatment data were collated from over 75 years of treatments conducted across the Great Basin (Pilliod et al., 2017, 2019). We defined the Great Basin region as the area described by Treminio et al. (2024), including the Snake River Plains. From the LTDL, we extracted treatment polygons that occurred within the Great Basin physiographic region (Figure 1b). Data from aerial seeding, closure, drill seeding, ground seeding, herbicide application, seedling planting, soil disturbance, vegetation disturbance, and weeds management treatment-polygons that overlapped with AIM monitoring plots (Figure 1a) were used to assess treatment effects on aeolian sediment transport (Table 1). The treatment types were selected on the basis of their high application frequencies and having the largest number of AIM monitoring plots from which aeolian horizontal sediment mass flux could be estimated. In addition, only LTDL data where sufficient metadata about treatment application and year of completion were available were used in our analyses.

**Figure 1.**
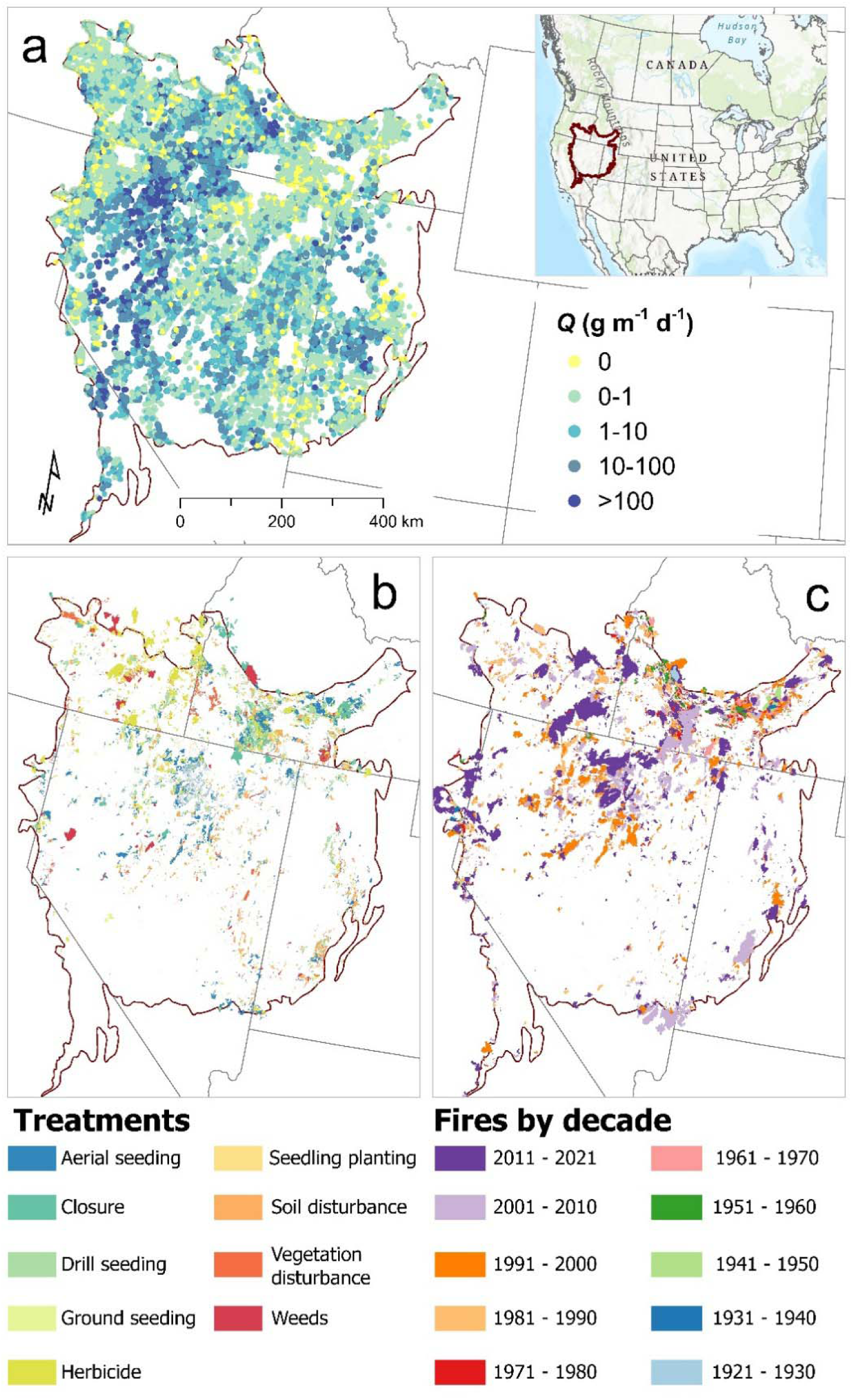
Location and distribution of predicted orders of magnitude in aeolian horizontal sediment flux, *Q* (g m^-1^ d^-1^), resulting from AERO for 20,022 Bureau of Land Management (BLM) Assessment, Inventory, and Monitoring (AIM) plots (a), rangeland treatments (b), and fire perimeters by decade of occurrence (c) across the Great Basin in the western United States (inset). Note that fire perimeters from 2021 have been grouped with those from the 2011-2020 decade.

**Table 1.** Definitions of rangeland treatments from the Land Treatment Digital Library evaluated for their impact on wind erosion in the Great Basin of the western United States acquired from https://www.usgs.gov/apps/ltdl/ (Pilliod et al., 2019; Pilliod and Welty, 2013).

| Treatment | Definition |
| --- | --- |
| *Aerial seeding* | The aerial application of seed mixtures using fixed-wing aircraft or helicopter. This treatment method covers large and hard to access areas quickly, delivering seeds to increase vegetation cover. |
| *Closure* | The establishment of a protected area through fencing or controlled access to reduce external disturbance from grazing or other human activity. |
| *Drill seeding* | A mechanical seeding method that employs a seed-drill to plant seeds at controlled depths and spacing. |
| *Ground seeding* | The application of seeds directly onto the soil either manually or by seed broadcasting equipment. |
| *Herbicide* | Application of chemical agents to control or eliminate undesirable vegetation that competes with native vegetation. |
| *Seedling planting* | Planting native seedlings and using seed balls to enhance germination, especially in dry conditions. |
| *Soil disturbance* | Physical alteration of soil structure and composition that modifies surface properties or chemical conditions to promote vegetation growth often through the use of chaining or discing machinery. |
| *Vegetation disturbance* | Treatment application that modifies the structure, composition, or function of vegetation through thinning, mowing, or chopping to reduce density. |
| *Weeds* | Control or eradication of invasive plant species especially non-native grasses and forbs |

The BLM AIM program collects data at spatially-stratified and randomly-sampled rangeland monitoring plots to assess the condition and trends of public lands (Taylor et al., 2014; Toevs et al., 2011). At each monitoring plot, the standard methods of Herrick et al. (2018), including line-point intercept, all-plant canopy gap intercept, and vegetation height methods were used to monitor the cover, structure, and composition of vegetation. The percent cover of plant functional groups and species of interest, vegetation height, and the size distribution of all-plant canopy gaps were calculated from raw field measurements using the *terradactyl* R package (McCord et al., 2022) and extracted from the Landscape Data Commons (McCord et al., 2023).

AIM monitoring plots sampled in the Great Basin between 2011-2023 were used as sampling units for assessing treatment effects on aeolian sediment transport rates (Figure 1a). Aeolian sediment transport, quantified as horizontal sediment mass flux (*Q*, g m^-1^ d^-1^) — an indicator of wind erosion and dust emission risk — was estimated for each AIM monitoring plot using the Aeolian EROsion (AERO) model (Edwards et al., 2022). Briefly, AERO is a physically based model that integrates rangeland monitoring data, surface soil properties, and meteorological conditions to estimate the horizontal and vertical movement of soil by wind (Edwards et al., 2022). AERO can be applied across landcover types and spatial scales (e.g., Treminio *et al*., 2024). Edwards et al. (2022) describe AERO parameterization using 453 parameter sets with a Generalized Likelihood Uncertainty Estimation framework to produce probabilistic estimates of *Q*. Here, the median AERO prediction was used to describe potential *Q* for monitoring plots following Treminio et al. (2024), using a 30-year (1981-2011) wind speed probability distribution derived from the National Centers for Environmental Prediction North American Regional Reanalysis product (Mesinger et al., 2006).

To examine effects of post-wildfire treatment, historical wildfire data were acquired from the Wildland Fire Trends Tool (Jeffries et al., 2022). Wildfire data described the locations of historical fires (Figure 1c), the number of fire occurrences within a 110-year period (1910 – 2021), and the most recent fire occurrence year at a 30 m spatial resolution (Jeffries et al., 2022). We excluded from analyses any prescribed burn treatments. Elevation varies greatly between low-lying basins and mountain ranges in the Great Basin and is negatively associated with wind erosion (Treminio et al., 2024). Elevation data for AIM monitoring plot locations were acquired from a 90 m digital elevation model for the western United States (Hanser, 2008). Annual precipitation influences primary production and wind erosion risk, thus mean annual precipitation for 2011-2023 was acquired from 4 km gridded PRISM data (PRISM Climate Group, 2024). Percent sand content that influences soil erodibility was acquired for each AIM plot location from 100 m gridded Soil Landscapes of the United States (Nauman et al., 2024). Major Land Resource Areas (MLRA) were used to account for physiographic differences in the locations of AIM plots across the Great Basin. MLRAs are regions with broadly comparable biotic potentials and limitations, having similar climate, soils, vegetation, elevation, topography, water resources, and land use (USDA-NRCS, 2022). Here, we assess treatment outcomes across MLRAs that include sagebrush steppe, salt-scrub desert, and pinyon-juniper woodlands.

### 2.2. Data preprocessing and compilation

Data preprocessing and covariate extractions were conducted in ArcGIS Pro v3.4.2 (ESRI, 2024). AIM plot locations were overlayed on GIS datasets of LTDL treatment polygons, historical wildfire perimeters, most recent wildfire rasters, elevation, percent sand content, precipitation, and MLRA. The Spatial Join tool with a one-to-many relationship was used to extract information from all treatment polygons that overlapped with plot locations. Treatment data included the major treatment type and the year the treatment application was completed. AIM plots that did not coincide with treatment polygons were retained in the preprocessing data compilation to serve as non-treated controls in subsequent analyses. For gridded datasets, the Extract Multi Values to Points tool was applied to obtain raster cell values coincident with AIM monitoring plot locations. Preprocessing resulted in 20,022 AIM monitoring plots that contained an estimate of *Q*, identification of rangeland treatments and the year it was completed, wildfire history and occurrence year, and percent cover estimates for foliar cover, bare ground and its distribution that control wind erosion (Treminio et al., 2024).

AIM monitoring plots were categorized into three distinct groups to account for rangeland treatment and wildfire effects on wind erosion. The first category (no-treatments) included monitoring plots where wildfires occurred prior to AIM sampling and no recorded rangeland treatment had taken place. These plots (N = 1,965) provide a baseline for wind erosion response to wildfire without confounding treatment effects. The second category (restoration) included monitoring plots with no wildfire history where restoration treatments occurred prior to AIM monitoring. These plots (N = 1,118) control for potential wind erosion response to restoration treatments without confounding effects of wildfire. The third category (post-wildfire rehabilitation) consisted of monitoring plots where wildfire occurred and a post-wildfire rehabilitation treatment was implemented before AIM monitoring. These plots (N = 2,054) enable assessment of wind erosion responses to the combined effects of fire and rehabilitation treatment. AIM monitoring plots without treatments or prior wildfire were assigned as controls for no-treatments and restoration (N = 12,529). Post-wildfire controls included plots where wildfire preceded between one and 99 years prior to AIM sampling and no rangeland treatments were applied (N = 1,965). We identified that multiple wildfires and treatments may occur within a single year or have occurred many times prior to AIM sampling for a given monitoring location, such that residual influences from preceding wildfires and treatments could have affected the vegetative cover dynamics and aeolian sediment transport responses. Due to sample size constraints, we were unable to employ statistical models or analytical methods that explicitly account for the variability associated with these effects. We acknowledge these potential sources of unaccounted variability and thus consider our estimates to be conservative approximations of effects.

Propensity score matching (PSM), using the *MatchIt* R package (Ho et al., 2011), was utilized to generate datasets that pair non-treated plots as controls to treated monitoring plots from each treatment within the previously identified categories (i.e., no treatments, restoration, post-wildfire rehabilitation). PSM enables the estimation of treatment effects by creating a balanced set of treated-control units based on shared covariates between samples that were treated or not treated (Ho et al., 2007). Logistic regression models were employed to estimate the probability that each plot would receive a particular treatment based on shared covariates. Covariates used for estimating the probability of treatment assignment and matching included the percent cover estimates of bare soil, total foliar cover, forb cover, grass cover, shrub cover, canopy gaps >1 m, elevation, and annual precipitation, year plot was sampled, and MLRA (Table S1). The biophysical covariates account for systematic differences between treated and control AIM plots that influence wind erosion in the Great Basin (Treminio et al., 2024). The “nearest neighbor” matching function with a 2:1 control to treated plot ratio was used to select non-treated plots that most closely match treated plots within the covariates space and logistic regression probabilities. To maximize covariate balance, a caliper length of 0.2 times the standard deviation in propensity score was employed. A caliper length sets the maximum difference between propensity scores of matched treated and control plots. Similar procedures for pairing control and treated units have been used for estimating treatment effects in rangeland restoration projects (e.g., Kluender et al., 2024; Pearson et al., 2016; Simler-Williamson and Germino, 2022). For no-treatment plots, the probability of being assigned to wildfire (treated) versus no-wildfire (control) served as the basis for PSM. Sampling with replacement of all available control plots, where possible, was conducted to further improve balance in restoration and post-wildfire rehabilitation categories during PSM. The total number of treated and non-treated plots by category before and after matching are described in Table 2. PSM balance between treated and non-treated AIM plots was assessed by comparing empirical cumulative density functions (eCDF) of covariates for the full unadjusted and the matched set of units. Love-plots comparing the standardized mean difference in the covariates between the full unadjusted and matched data were used as a second assessment of balance. Love-plots and eCDF’s are described in Supplementary Figures S1-S17.

**Table 2.**
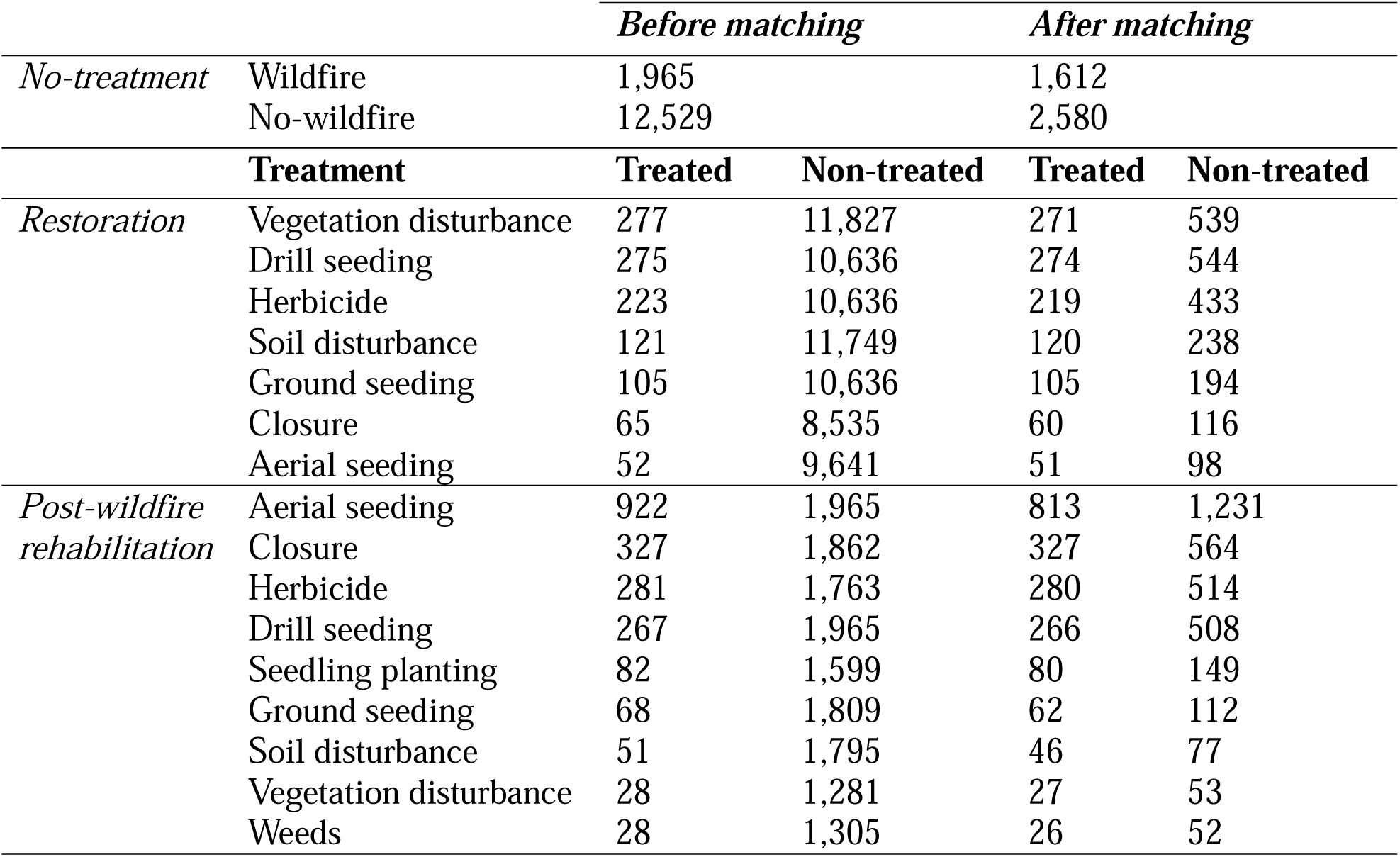
Comparison of the number of monitoring plots available before and after propensity score matching. No-treatment plots are those with no treatment history that could influence aeolian sediment transport response to wildfire occurrence. Restoration plots are those where a treatment was applied but with no wildfire history that could affect aeolian sediment transport responses to treatment. Post-wildfire rehabilitation plots are those where wildfire occurred before treatment, and treatment occurred prior to AIM monitoring. Non-treated control plots for post-wildfire rehabilitation are plots where wildfire occurred before AIM monitoring with no history of treatment.

|  |  | *Before matching* |  | *After matching* |  |
| --- | --- | --- | --- | --- | --- |
| *No-treatment* | Wildfire | 1,965 |  | 1,612 |  |
|  | No-wildfire | 12,529 |  | 2,580 |  |
|  | **Treatment** | **Treated** | **Non-treated** | **Treated** | **Non-treated** |
| *Restoration* | Vegetation disturbance | 277 | 11,827 | 271 | 539 |
|  | Drill seeding | 275 | 10,636 | 274 | 544 |
|  | Herbicide | 223 | 10,636 | 219 | 433 |
|  | Soil disturbance | 121 | 11,749 | 120 | 238 |
|  | Ground seeding | 105 | 10,636 | 105 | 194 |
|  | Closure | 65 | 8,535 | 60 | 116 |
|  | Aerial seeding | 52 | 9,641 | 51 | 98 |
| *Post-wildfire rehabilitation* | Aerial seeding | 922 | 1,965 | 813 | 1,231 |
|  | Closure | 327 | 1,862 | 327 | 564 |
|  | Herbicide | 281 | 1,763 | 280 | 514 |
|  | Drill seeding | 267 | 1,965 | 266 | 508 |
|  | Seedling planting | 82 | 1,599 | 80 | 149 |
|  | Ground seeding | 68 | 1,809 | 62 | 112 |
|  | Soil disturbance | 51 | 1,795 | 46 | 77 |
|  | Vegetation disturbance | 28 | 1,281 | 27 | 53 |
|  | Weeds | 28 | 1,305 | 26 | 52 |

### 2.3. Statistical analyses

The *marginaleffects* R package (Arel-Bundock et al., 2024) was used to conduct G-computation to estimate treatment effects on aeolian sediment transport across the Great Basin. This method — a statistical approach for causal inference when randomized controlled trials are not feasible — enables the estimation of treatment effects from observational data (Naimi et al., 2017; Schafer and Kang, 2008; Snowden et al., 2011). A five-step approach was employed to estimate treatment effects on wind erosion. First, predicted aeolian horizontal sediment flux (*Q*, g m ¹ d ¹) was natural-log transformed when *Q* > 0. For *Q* = 0, a value of 1 was added to allow inclusion in the natural-log transformation. The natural-log transformed *Q* (Ln *Q*) served as the response variable to better fit the assumption of normally distributed errors in multiple linear regression. Second, multilinear regression models with interaction terms between treatment and the covariates used in PSM were fit for each treatment within each category type to predict responses from observed data. Including these covariates in the outcome model ensures a doubly robust estimate of treatment effects by addressing model misspecification and balancing covariates between treated and non-treated plots (Hall et al., 2014). Third, counterfactual outcomes were generated by applying the fitted model(s) to simulate results under alternative treatment conditions. To illustrate, a fitted model is used to predict outcomes for each plot in the dataset as if it had either not been treated (control) or had received treatment. Fourth, the average predicted counterfactual outcomes for both treatment and control scenarios were calculated and tested to determine whether they differ significantly from zero. This revealed the impact of each treatment, or lack thereof, on potential wind erosion. Fifth, the overall treatment effect was determined by calculating the difference between the average predicted outcomes for treated and control scenarios and testing whether this difference was significantly different from zero. We note that under G-computation, the coefficients of covariates from fitting multiple linear regression cannot be interpreted as causal relationships to treatment designations and thus their effect on Ln *Q*. Therefore, the effect of a covariates used in the regression model, such as the percentage of sand are not discussed. In the results, we present treatment outcomes in terms of 1) comparison of the estimated Ln *Q* magnitude between treatments and corresponding untreated controls, and 2) the treatment effect as the statistical difference between treatments and its control.

For no-treatment plots, we estimated how wildfire influenced sediment transport relative to plant cover by comparing between plots sampled within one-year post-wildfire and plots with no wildfire history, sampled in the same year. The comparison focused on percent cover of five plant functional groups: annual forbs, annual grasses, perennial forbs, perennial grasses, and sagebrush.

To further investigate the role of the five plant functional groups, post-hoc analyses of statistically significant overall treatment effects (i.e., results of the fifth step above) on *Q* were conducted using permutation tests for restoration and post-wildfire rehabilitation plots. Permutation tests assessed whether mean cover for the five plant functional groups differed between treated and control plots. The plant groups were chosen because: 1) they provide finer-grained indices of vegetation cover covariates used in PSM and G-computation; 2) their cover is influenced by, or results from, restoration and post-wildfire rehabilitation treatments in the Great Basin; and 3) they avoid the circularity of reusing covariates employed in PSM and G-computation. Permutation tests were used because they do not require assumptions of normality or homoscedasticity. In this procedure, treatment assignments were repeatedly shuffled within the dataset to simulate a null hypothesis of no difference between groups. For each permutation, a distribution under the null hypothesis was created for the mean difference between groups. The observed difference in mean plant functional group cover between treatment and untreated controls was then compared to this distribution. The *p*-value was computed as the proportion of permuted test statistics that were as extreme as, or more than, the observed difference. A total of N = 10,000 permutations were used to generate the null distributions, and a significance level of α = 0.05 was applied. All statistical analyses were conducted in R v. 4.4.2 (R Core Team, 2024) and RStudio v. 27771613 (RStudio Team, 2025). Figures were created using the *ggplot2* R package (Wickham, 2016).

## 3. Results

### 3.1. Aeolian sediment transport response to wildfire without treatments

We present the response of natural log-transformed *Q*, where a unit change in value represents an order of magnitude change in the sediment transport rate. The predicted mean aeolian sediment transport response (Ln *Q*; g m ¹ d ¹) was significantly less than zero for burned and unburned monitoring plots with no rangeland treatment history (Figure 2a). Plots with wildfire history had a mean predicted aeolian sediment transport response of −0.942 Ln *Q* (*p <* 0.001), compared to −0.872 Ln *Q* (*p <* 0.001) for plots with no fire history (Table 3). The overall effect of wildfire on aeolian sediment transport was −0.070 Ln *Q* (*p =* 0.364), suggesting that plots with wildfire history, on average, experience slightly less aeolian sediment transport than those without wildfire history. However, this difference was not statistically significant (Table 4). One-year post-wildfire monitoring revealed that median estimates of *Q* in burned plots were higher than in plots with no wildfire history in some but not all years (Figure 2b). Elevated *Q* within 12 months of wildfire resulted from reduced plant functional group cover, but annual and perennial grasses can recover in this time to levels matching unburned plots (Figure 3) and to percent cover ranges linked to the greatest decline in wind erosion.

**Figure 2.**
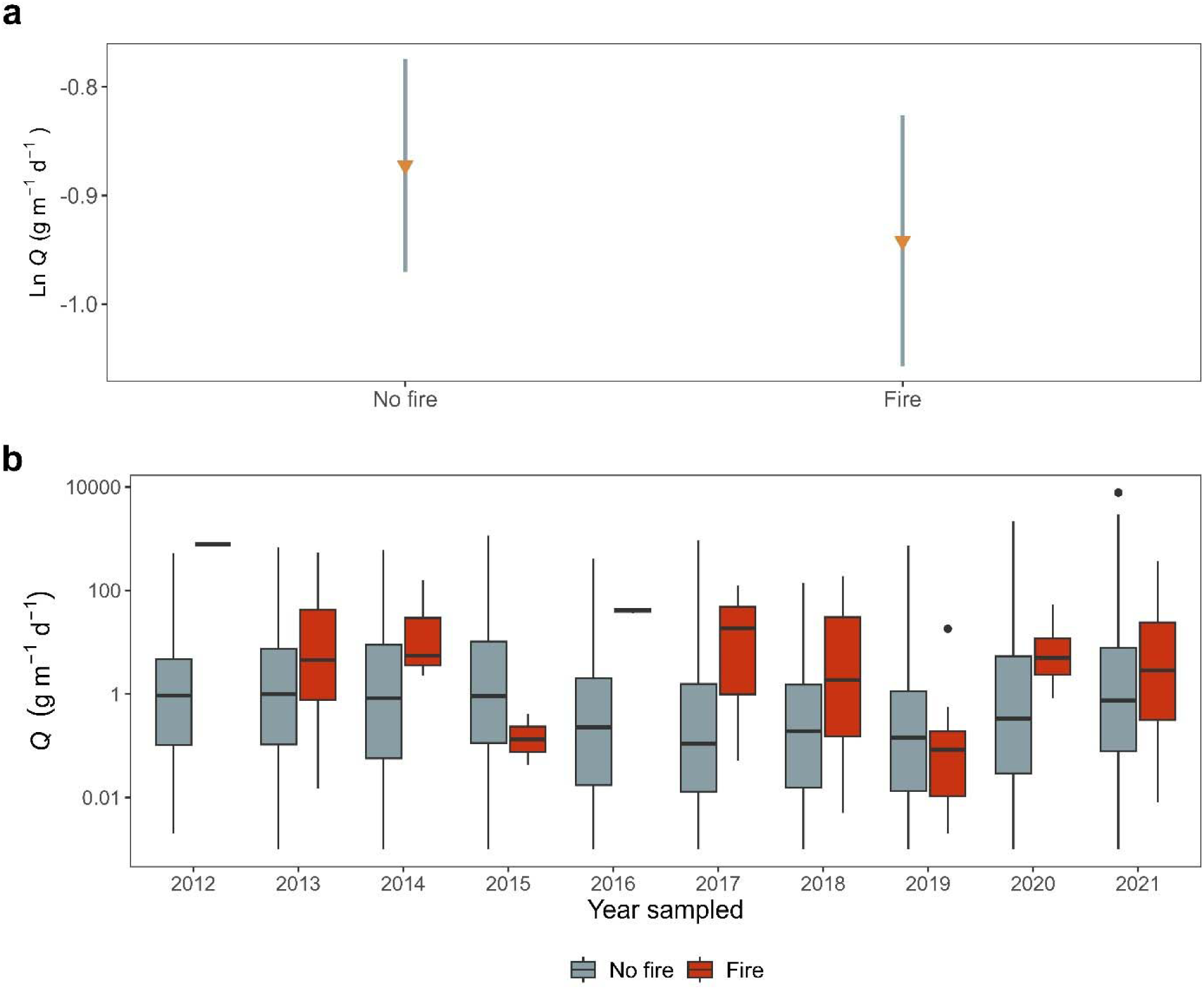
Comparison of mean predicted aeolian sediment transport (Ln *Q*) response for a) plots that have a history of wildfire but no prior treatments. Bars represent 95% confidence interval of the predicted mean (orange triangles) Ln *Q* response. Statistical significance results for a) are described in Table 3. Panel b) is a box-and-whisker plot illustrating the distribution of aeolian horizontal sediment flux (*Q*) between AIM monitoring plots with a history of fire and those without. In box-and-whisker plots, bars above and below boxes represent the range of values within ±1.5 times the interquartile range and points represent outlier values outside this range. The wildfire-affected plots represent data collected within one-year post-wildfire, while the no-fire plots correspond to data sampled during the same years as their wildfire-affected counterparts. Note that plot (b) is not natural log-transform but the y-axis scale is in base-10 orders of magnitude.

**Figure 3.**
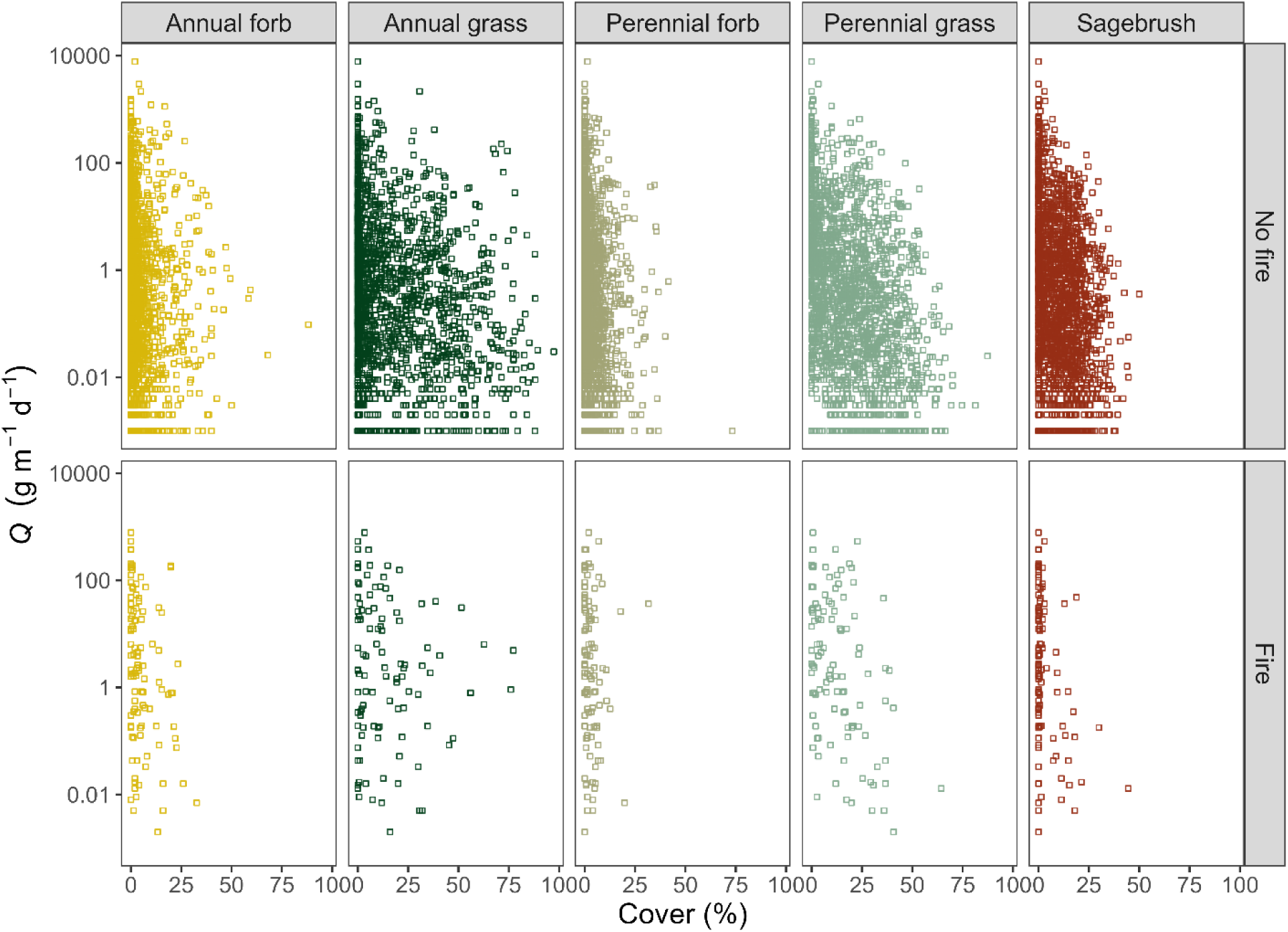
Comparison of aeolian horizontal sediment flux (*Q*) as a function of plant cover between AIM monitoring plots sampled within one-year post-wildfire and plots with no fire history, both sampled in the same years. Note that plots are not natural log-transformed, and the y-axis scale is base-10 orders of magnitude.

**Table 3.**
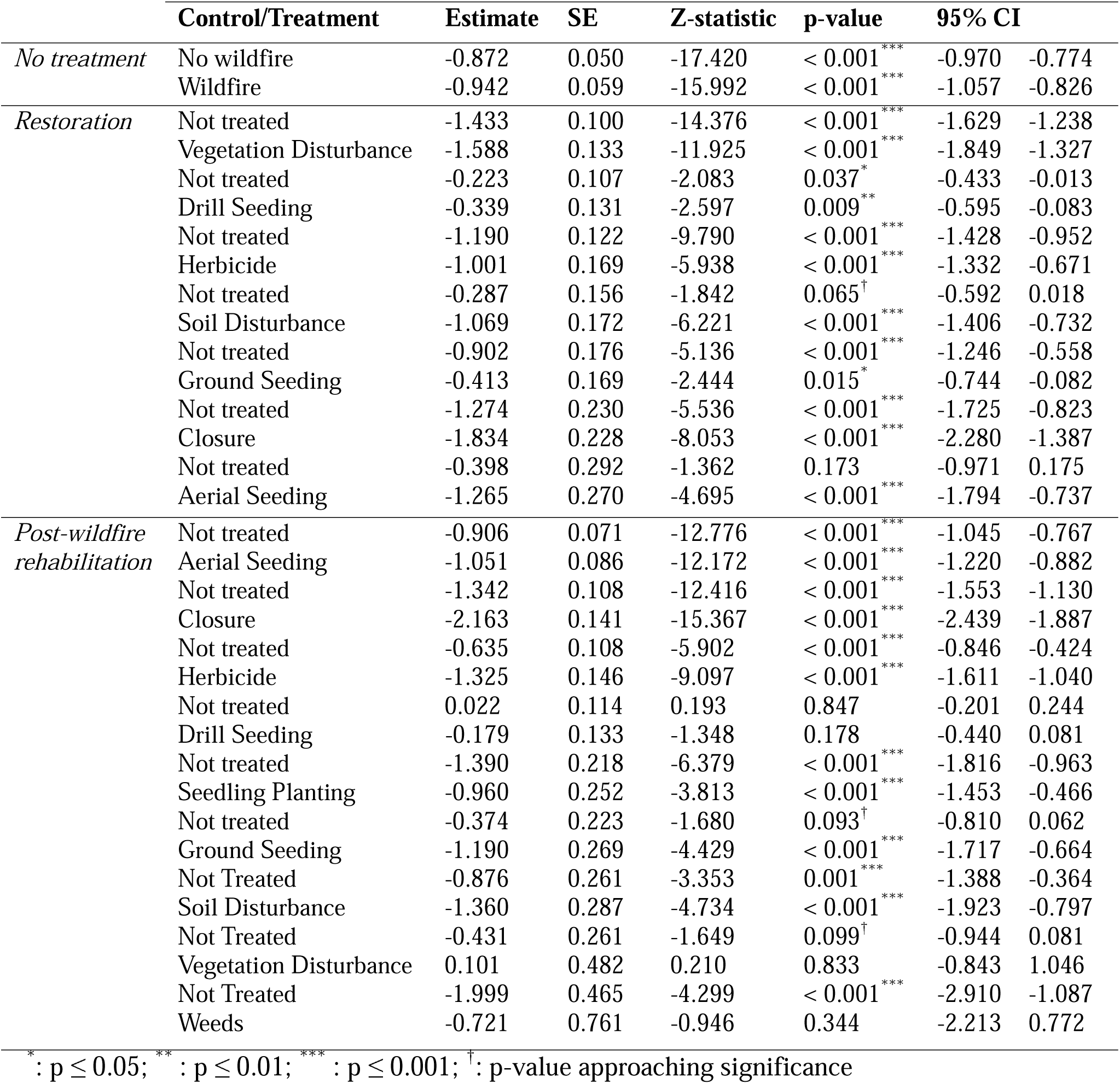
Summary results table of statistical tests comparing the mean predicted estimates of aeolian sediment transport (Ln *Q*) responses for each treatment against their corresponding untreated controls. Each estimate was tested to determine whether it significantly differed from zero Ln *Q*.

**Table 4.** Summary of statistical tests assessing treatment effects on aeolian sediment transport. The treatment effect was evaluated as treatment minus untreated control mean predicted response.

|  | Comparison | Estimate | SE | Z-statistic | p-value | 95 % CI |  |
| --- | --- | --- | --- | --- | --- | --- | --- |
| *No treatment* | Fire - No fire | -0.070 | 0.077 | -0.908 | 0.364 | -0.220 | 0.081 |
| *Restoration* | Vegetation disturbance - Not treated | -0.155 | 0.162 | -0.959 | 0.338 | -0.472 | 0.162 |
|  | Drill seeding - Not treated | -0.116 | 0.172 | -0.671 | 0.502 | -0.453 | 0.222 |
|  | Herbicide - Not treated | 0.189 | 0.208 | 0.907 | 0.364 | -0.219 | 0.596 |
|  | Soil disturbance - Not treated | -0.783 | 0.253 | -3.097 | 0.002** | -1.278 | -0.287 |
|  | Ground seeding - Not treated | 0.489 | 0.253 | 1.935 | 0.053<sup>†</sup> | -0.006 | 0.984 |
|  | Closure - Not treated | -0.560 | 0.324 | -1.726 | 0.084<sup>†</sup> | -1.195 | 0.076 |
|  | Aerial seeding - Not treated | -0.867 | 0.398 | -2.177 | 0.030* | -1.648 | -0.086 |
| *Post-fire rehabilitation* | Aerial seeding - Not treated | -0.145 | 0.112 | -1.294 | 0.196 | -0.366 | 0.075 |
|  | Closure - Not treated | -0.821 | 0.183 | -4.481 | < 0.001*** | -1.181 | -0.462 |
|  | Herbicide - Not treated | -0.690 | 0.185 | -3.739 | < 0.001*** | -1.052 | -0.328 |
|  | Drill seeding - Not treated | -0.201 | 0.173 | -1.164 | 0.244 | -0.540 | 0.138 |
|  | Seedling planting - Not treated | 0.430 | 0.368 | 1.167 | 0.243 | -0.292 | 1.152 |
|  | Ground seeding - Not treated | -0.817 | 0.345 | -2.367 | 0.018* | -1.493 | -0.140 |
|  | Soil disturbance - Not treated | -0.484 | 0.403 | -1.200 | 0.230 | -1.274 | 0.306 |
|  | Vegetation disturbance - Not treated | 0.533 | 0.528 | 1.009 | 0.313 | -0.502 | 1.567 |
|  | Weeds - Not treated | 1.278 | 0.909 | 1.407 | 0.160 | -0.503 | 3.059 |

\* : p ≤ 0.05; \*\* : p ≤ 0.01; \*\*\* : p ≤ 0.001; <sup>†</sup>: p-value approaching significance

### 3.2. Aeolian sediment transport response to restoration treatments without wildfire

All restoration treatments and controls — except for soil disturbance and aerial seeding controls — exhibited significant negative predicted Ln *Q* responses (Table 3). Mean predicted aeolian sediment transport responses for most restoration treatments were more negative than those of untreated control plots except for herbicide and ground seeding treatments (Figure 4a). Herbicide and ground seeding treatments had less negative aeolian sediment transport responses than non-treated controls, suggesting an increase in *Q* (*p <* 0.05; Table 3). Among treatments significantly different from zero, closure had the largest reduction (−1.834 Ln *Q*; *p <* 0.001), while drill seeding treatments had the least negative (−0.339 Ln *Q*; *p =* 0.009). Soil disturbance and aerial seeding were the only treatments that resulted in a treatment effect (e.g., the statistical difference between treatment and control not equal to zero Ln *Q*). This reduction was indicated by the negative direction of mean aeolian sediment transport values accounting for the response in controls, with soil disturbance showing a change of −0.783 Ln *Q* (*p* = 0.002) and aerial seeding showing a change of −0.867 Ln *Q* (*p* = 0.03) (Table 4).

**Figure 4.**
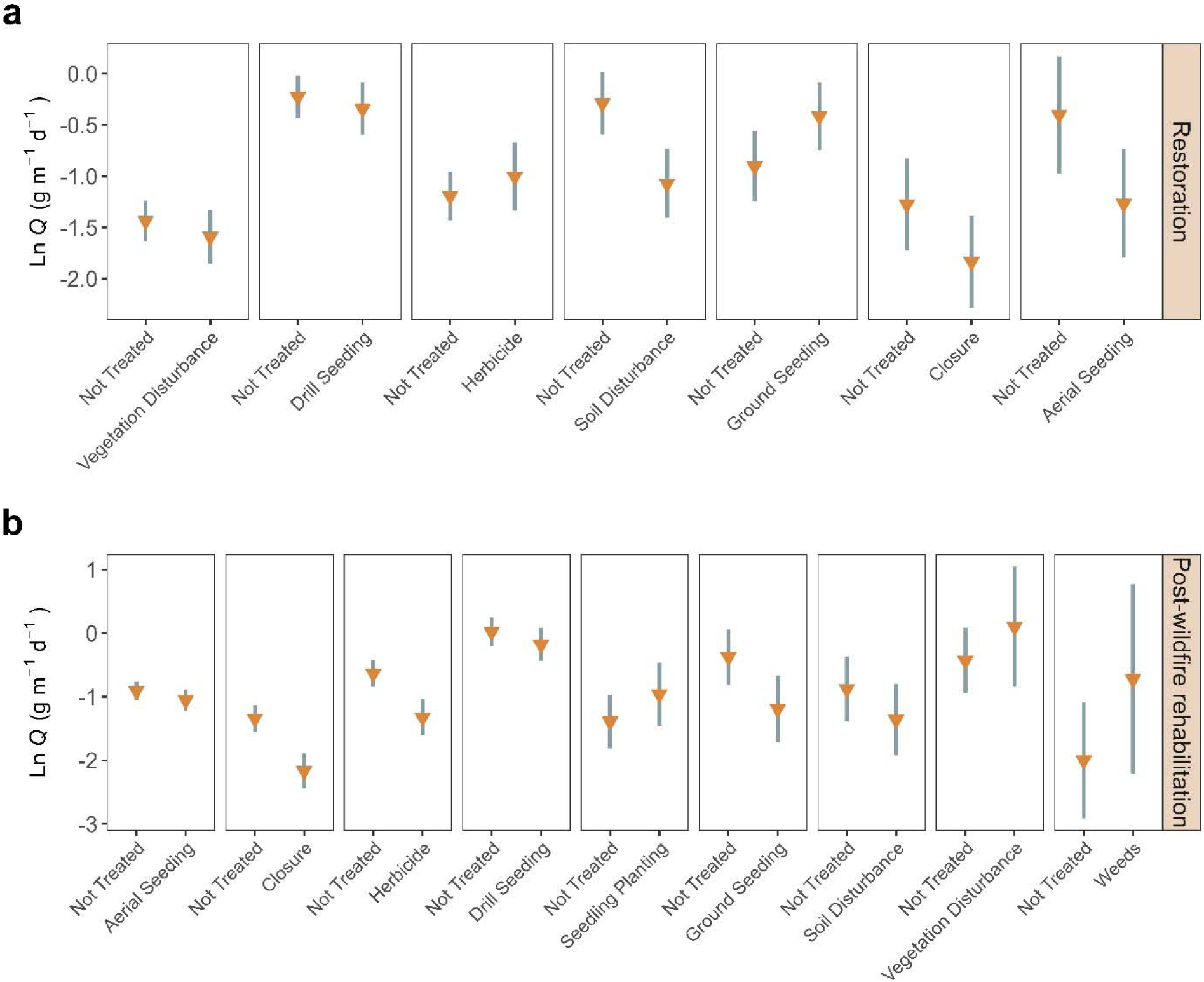
Comparison of mean predicted aeolian sediment transport (Ln *Q*) responses for a) restoration plots with no history of fire and where treatments occurred before AIM monitoring; and b) post-wildfire rehabilitation where fire occurred before treatment and treatment occurred before AIM monitoring. Bars represent 95% confidence interval of the predicted mean (orange triangles) Ln *Q* response. Statistical significance results are described in Table 3.

For permutation tests, distribution values less than zero indicate higher cover in non-treated plots than in treated plots. In contrast, distribution values greater than zero were indicative of higher plant functional group cover in treated compared to non-treated plots. Permutation tests showed significantly greater annual grass cover (−7.35%; *p =* 0.001) and perennial forb cover (−1.54%; *p =* 0.023) in non-treated plots for soil disturbance while perennial grass cover was 6.48% higher in treated plots (*p =* 0.065; Table 5). Permutation tests for aerial seeding treatment effects demonstrated higher annual grass cover in non-treated plots (−7.21%; *p =* 0.045) and higher but not significantly different perennial grass cover in treated plots (4.36%; *p =* 0.188). Comparison of the observed distributions of functional group cover by restoration treatment also found median annual grass to be lower and perennial grass cover to be greater in soil disturbance (bottom panel Figure 5) and aerially seeding plots (bottom panel Figure 6).

**Figure 5.**
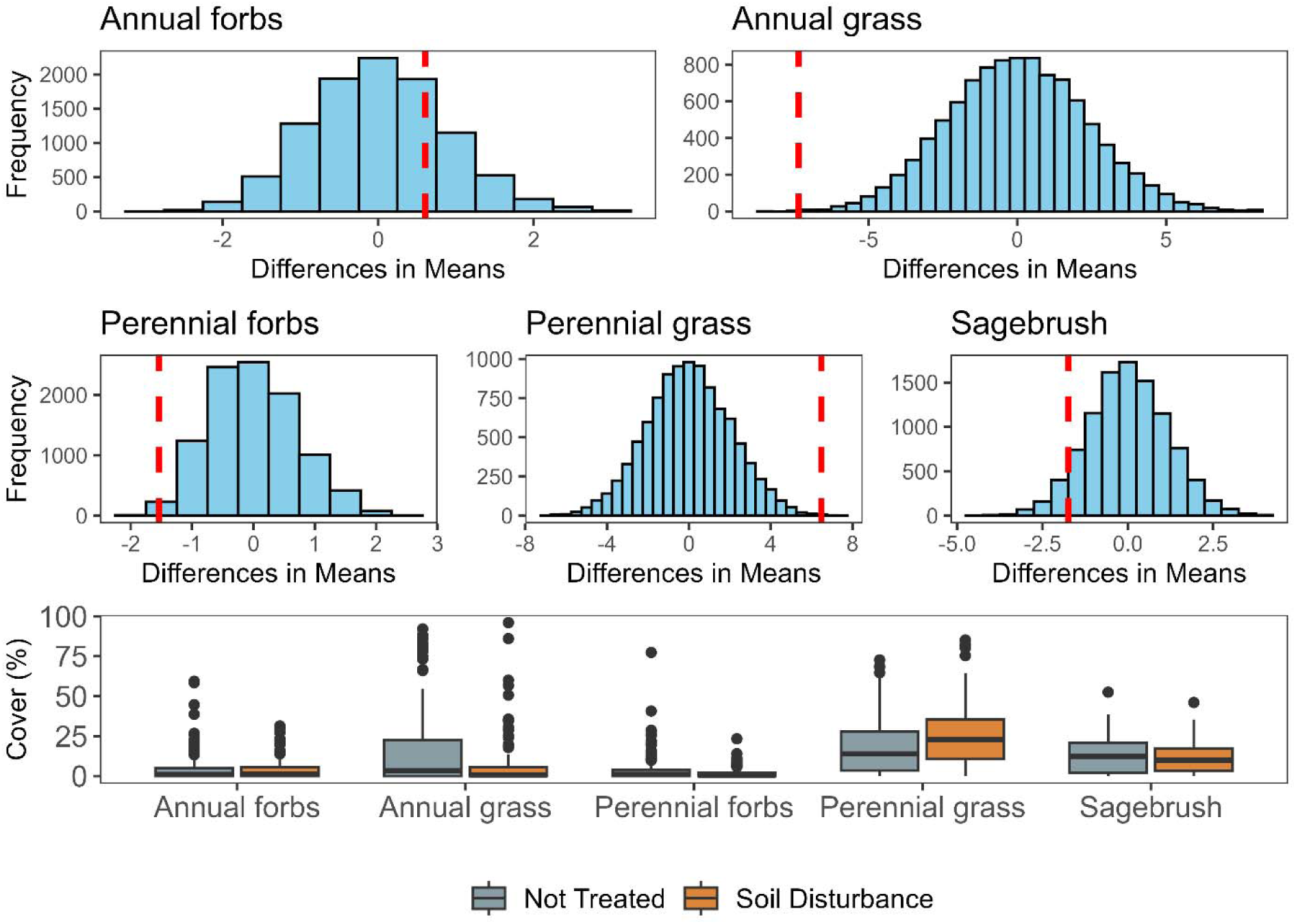
Permutation-test null distributions of mean differences in percent cover for five plant functional groups, comparing soil disturbance treatments to untreated controls for restoration plots (top two rows). Red vertical lines indicate the observed differences based on the data. Distributional values greater than zero indicate larger amounts of cover in treated plots. Distributional vales less than zero indicate larger amounts of cover in non-treated plots. The bottom panel shows box-and-whisker plots describing the observed data distributions of functional group cover by treatment. In box-and-whisker plots, bars above and below boxes represent the range of values within ±1.5 times the interquartile range and points represent outlier values outside this range.

**Figure 6.**
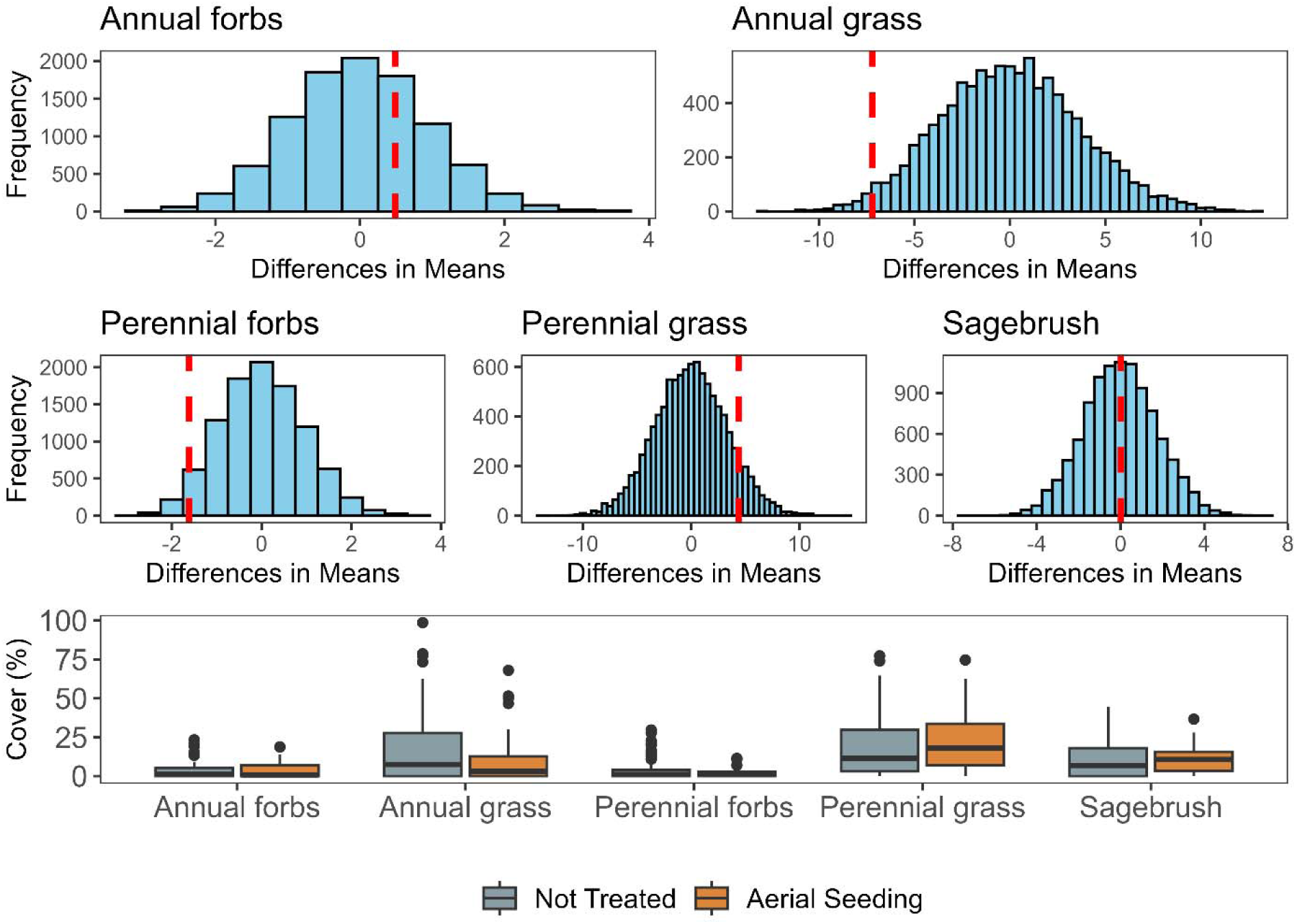
Permutation-test null distributions of mean differences in percent cover for five plant functional groups, comparing aerial seeding treatments to untreated controls for restoration plots (top two rows). Red vertical lines indicate the observed differences based on the data. Distributional values greater than zero indicate larger amounts of cover in treated plots. Distributional vales less than zero indicate larger amounts of cover in non-treated plots. The bottom panel shows box-and-whisker plots describing the observed data distributions of functional group cover by treatment. In box-and-whisker plots, bars above and below boxes represent the range of values within ±1.5 times the interquartile range and points represent outlier values outside this range.

**Table 5.**
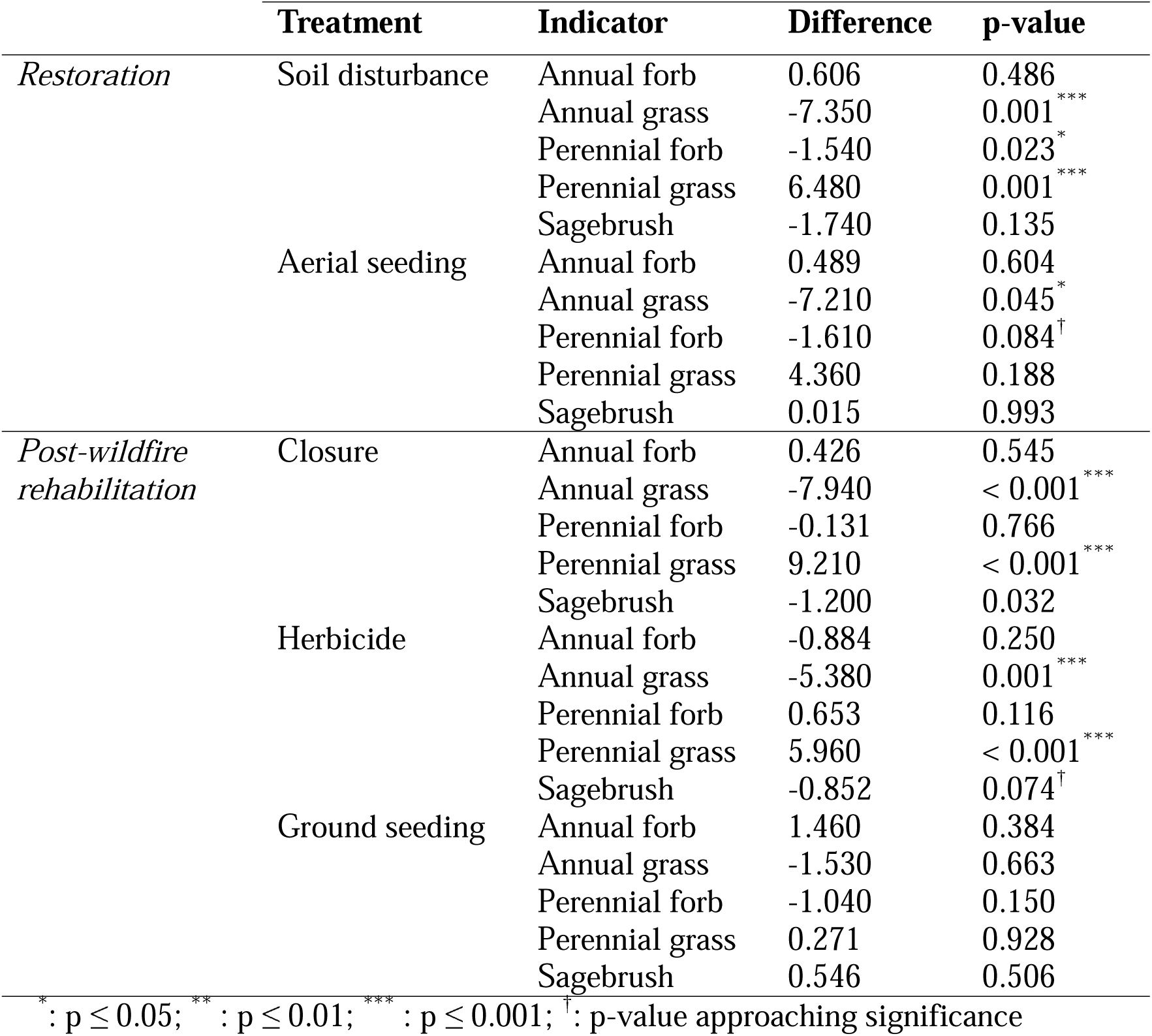
Comparison of mean percent cover differences for five functional group indicators, with significance assessed using permutation tests. Differences are calculated as the mean treatment minus the mean non-treated cover measured for each indicator. Difference values are represent as red vertical lines in Figure 5 through Figure 9.

|  | Treatment | Indicator | Difference | p-value |
| --- | --- | --- | --- | --- |
| *Restoration* | Soil disturbance | Annual forb | 0.606 | 0.486 |
|  |  | Annual grass | -7.350 | 0.001*** |
|  |  | Perennial forb | -1.540 | 0.023* |
|  |  | Perennial grass | 6.480 | 0.001*** |
|  |  | Sagebrush | -1.740 | 0.135 |
|  | Aerial seeding | Annual forb | 0.489 | 0.604 |
|  |  | Annual grass | -7.210 | 0.045* |
|  |  | Perennial forb | -1.610 | 0.084<sup>†</sup> |
|  |  | Perennial grass | 4.360 | 0.188 |
|  |  | Sagebrush | 0.015 | 0.993 |
| *Post-wildfire rehabilitation* | Closure | Annual forb | 0.426 | 0.545 |
|  |  | Annual grass | -7.940 | < 0.001*** |
|  |  | Perennial forb | -0.131 | 0.766 |
|  |  | Perennial grass | 9.210 | < 0.001*** |
|  |  | Sagebrush | -1.200 | 0.032 |
|  | Herbicide | Annual forb | -0.884 | 0.250 |
|  |  | Annual grass | -5.380 | 0.001*** |
|  |  | Perennial forb | 0.653 | 0.116 |
|  |  | Perennial grass | 5.960 | < 0.001*** |
|  |  | Sagebrush | -0.852 | 0.074<sup>†</sup> |
|  | Ground seeding | Annual forb | 1.460 | 0.384 |
|  |  | Annual grass | -1.530 | 0.663 |
|  |  | Perennial forb | -1.040 | 0.150 |
|  |  | Perennial grass | 0.271 | 0.928 |
|  |  | Sagebrush | 0.546 | 0.506 |

\* : $p \leq 0.05$ ; \*\* : $p \leq 0.01$ ; \*\*\* : $p \leq 0.001$ ; <sup>†</sup> : p-value approaching significance

### 3.3 Aeolian sediment transport response to post-wildfire rehabilitation treatments

Most rehabilitation treatments and their controls showed statistically significant negative mean Ln *Q* responses (Table 3). Mean predicted responses were more negative for aerial seeding, closure, herbicide, ground seeding, and soil disturbance treatments than for their respective non-treated controls (Figure 4b). Among these, closure treatments showed the largest negative mean response (−2.163 Ln Q; p < 0.001), whereas aerial seeding exhibited the smallest negative mean response ( −1.051 Ln Q; p < 0.001). Estimated treatment effects also trended negatively, but only closure (−0.821 Ln *Q*; *p <* 0.001), herbicide (−0.690 Ln *Q*; *p <* 0.001), and ground seeding (−0.817 Ln *Q*; *p =* 0.018) significantly differed from zero (Table 4).

Permutation tests for closure treatments revealed significant differences in vegetation cover where annual grass was higher in non-treated plots (−7.94%; *p =* 0.001), and perennial grass cover was significantly greater in treated plots (9.21%; *p <* 0.001; see Table 5 and Figure 7). Permutation tests of herbicide treatments showed a response pattern similar to closure treatments, such that non-treated control plots had significantly higher annual grass cover (−5.38%; *p =* 0.001) and treated plots exhibited a significantly higher perennial grass cover (5.96%; *p <* 0.001; see Table 5 and Figure 8). Permutation tests for ground seeding treatments did not reveal any significant differences in functional plant group cover between treated and untreated control plots (Table 5 and Figure 9). Although measured mean cover estimates for annual forbs (1.46%; *p =* 0.384), perennial grasses (0.27%; *p =* 0.928), and sagebrush (0.56%; *p =* 0.506) were slightly higher in treated plots, these differences were not statistically significant. Median cover estimates for plant function group cover showed similarly negligible differences between ground-seeded and control plots (Figure 9, bottom panel).

**Figure 7.**
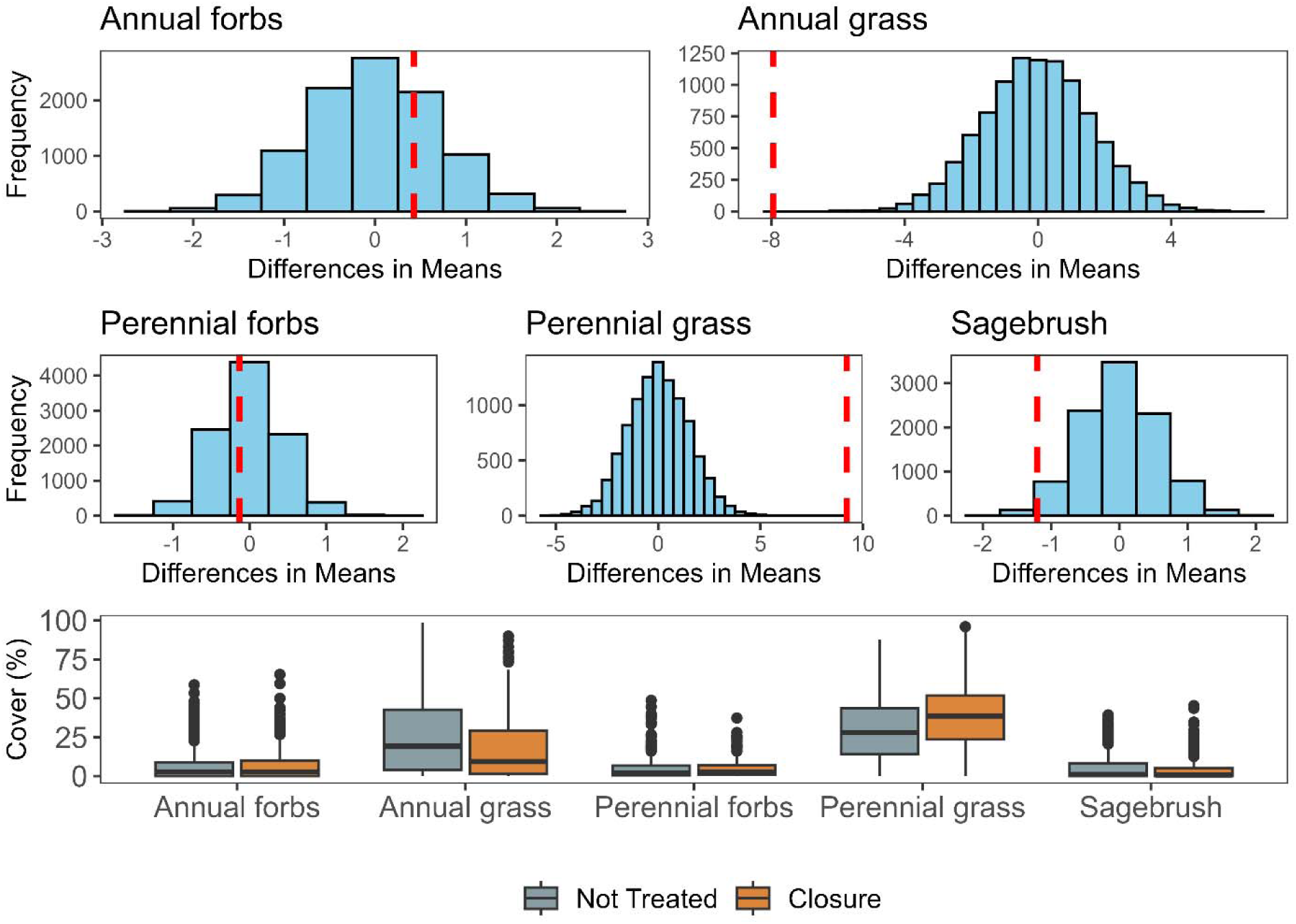
Permutation-test null distributions of mean differences in percent cover for five plant functional groups, comparing closure treatments to untreated controls for post-wildfire rehabilitation plots (top two panels). Red vertical lines indicate the observed differences based on the data. Distributional values greater than zero indicate larger amounts of cover in treated plots. Distributional vales less than zero indicate larger amounts of cover in non-treated plots. The bottom panel shows box-and-whisker plots describing the observed data distributions of functional group cover by treatment. In box-and-whisker plots, bars above and below boxes represent the range of values within ±1.5 times the interquartile range and points represent outlier values outside this range.

**Figure 8.**
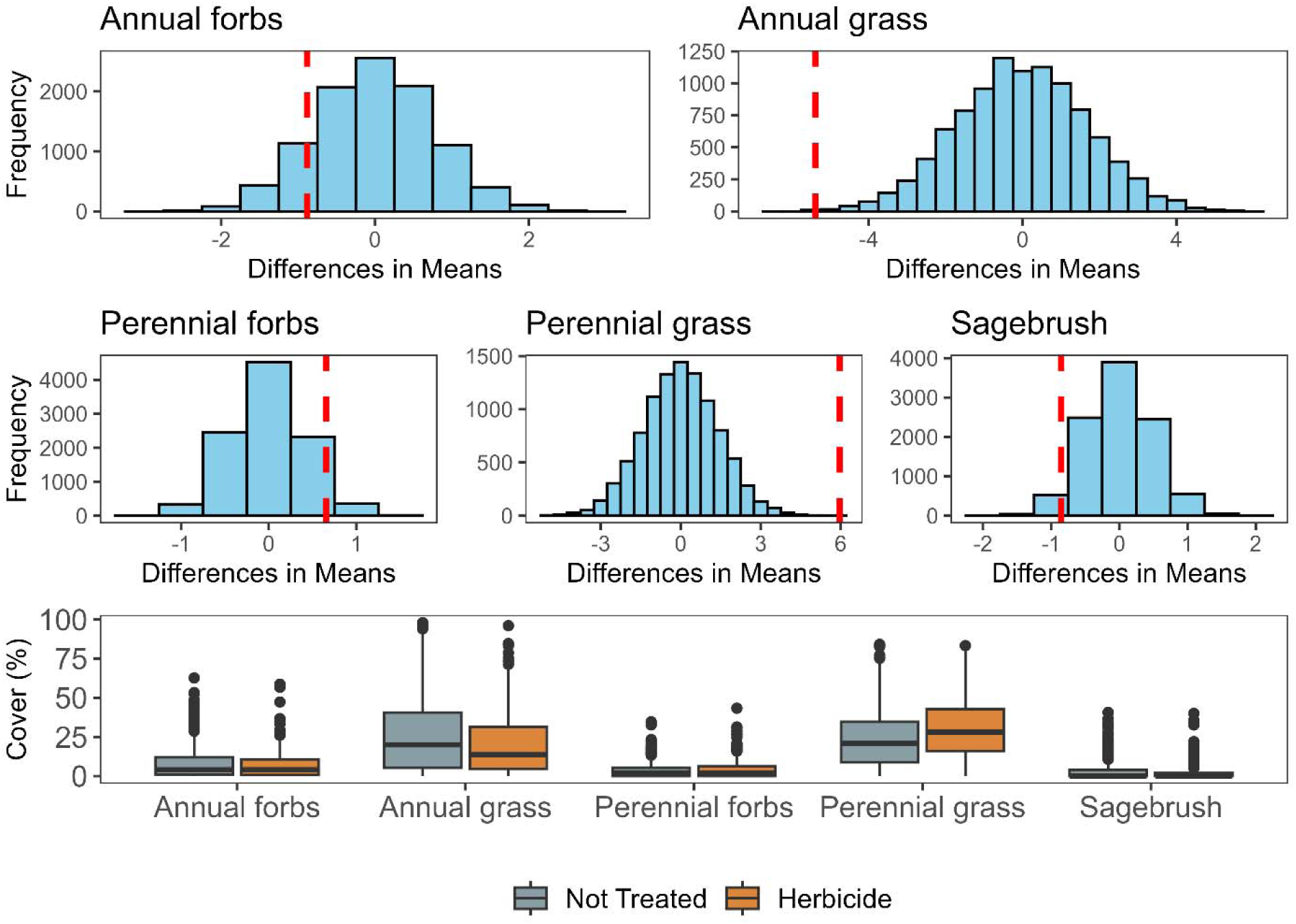
Permutation-test null distributions (blue bars) of mean differences in percent cover for five plant functional groups, comparing herbicide treatments to untreated controls for post-wildfire rehabilitation plots (top two panels). Red vertical lines indicate the observed differences based on the data. Distributional values greater than zero indicate larger amounts of cover in treated plots. Distributional vales less than zero indicate larger amounts of cover in non-treated plots. The bottom panel shows box-and-whisker plots describing the observed data distributions of functional group cover by treatment. In box-and-whisker plots, bars above and below boxes represent the range of values within ±1.5 times the interquartile range and points represent outlier values outside this range.

**Figure 9.**
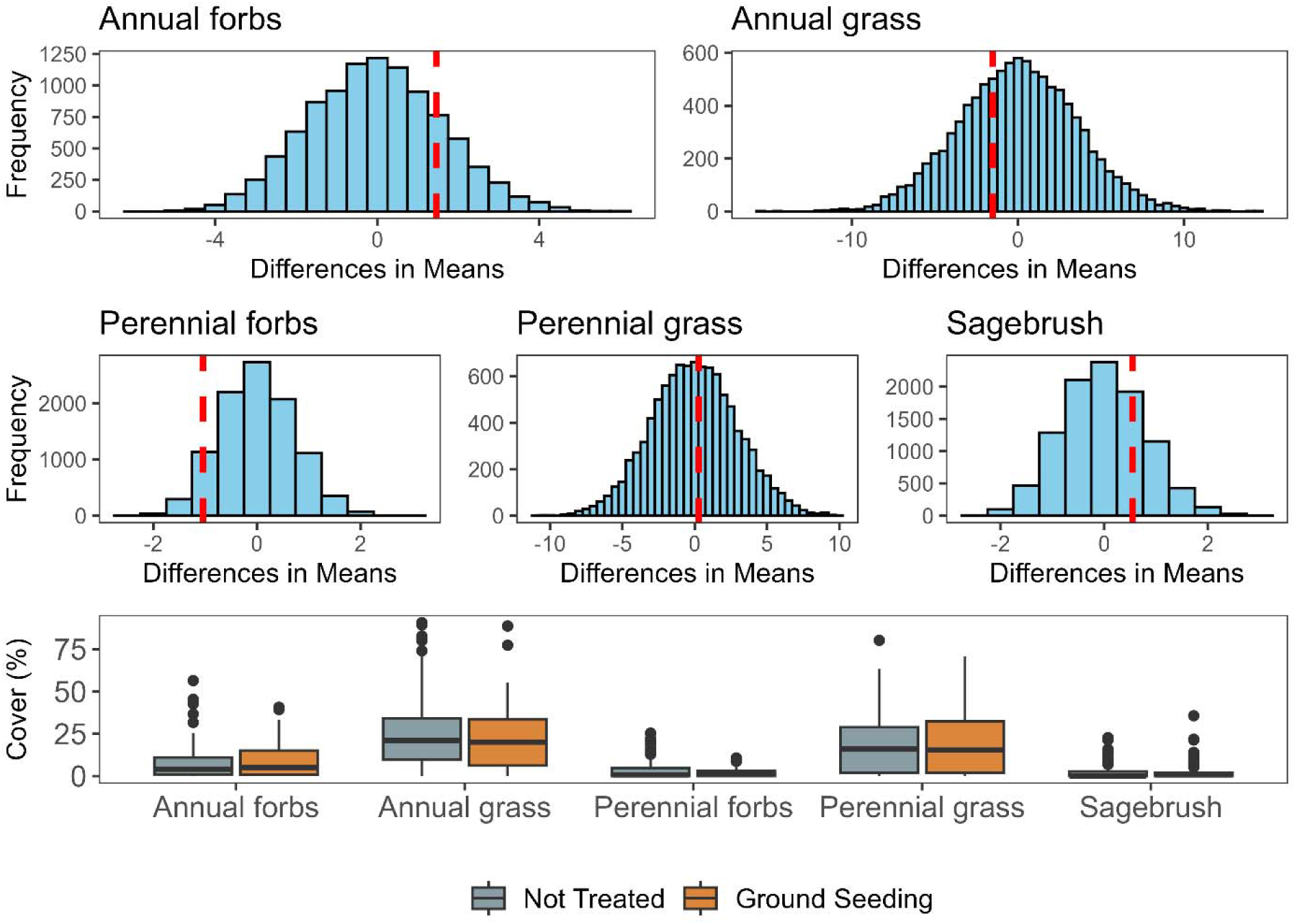
Permutation-test null distributions (blue bars) of mean differences in percent cover for five plant functional groups, comparing ground seeding treatments to untreated controls for post-wildfire rehabilitation plots (top two panels). Red vertical lines indicate the observed differences based on the data. Distributional values greater than zero indicate larger amounts of cover in treated plots. Distributional vales less than zero indicate larger amounts of cover in non-treated plots. The bottom panel shows box-and-whisker plots describing the observed data distributions of functional group cover by treatment. In box-and-whisker plots, bars above and below boxes represent the range of values within ±1.5 times the interquartile range and points represent outlier values outside this range.

## 4. Discussion

Integrating wind erosion and SDS assessments into dryland restoration and rehabilitation initiatives offers valuable insights into treatment outcomes that can enhance mitigation. This study supports such integration by using counterfactual model-estimates from observational monitoring data and the AERO model to determine that some restoration and post-wildfire rehabilitation treatments have mitigated wind erosion risk in the Great Basin over the period 2011-2023. While responses were sometimes small (Ln *Q* < 0; g m^-1^ d^-1^), the combined effects of wildfire occurrence, restoration and rehabilitation treatments, and their interaction suggest a tendency for reducing mean predicted aeolian sediment transport rates. In most cases, restoration and post-wildfire rehabilitation treatment responses were more negative (i.e., magnitude comparisons) than the predicted mean aeolian sediment transport rates for untreated controls. Although the effects of most treatments were not different from zero (i.e., statistical difference), the direction of the treatment effect consistently tended toward reduced aeolian sediment transport rates across comparisons. Moreover, in monitoring plots where treatment effects were statistically significant, we observed differences in annual and/or perennial grass cover between treated sites and untreated controls, that are described below. These findings suggest that restoration and rehabilitation treatments across the Great Basin, have likely increased annual and/or perennial grass cover and decreased bare ground, which in turn reduced the susceptibility of landscapes to wind erosion and dust emission.

Rangeland monitoring plots with a history of wildfire but no treatments had lower mean predicted aeolian sediment transport rates than plots with no fire history. This result may be attributed to most of the plant functional group cover indicators (except perennial forbs) showing recovery to at least 20% cover, within which aeolian sediment transport rates experience the greatest reductions (Webb et al., 2025; Wojcikiewicz et al., 2023). We found that within one-year post-wildfire, vegetation cover levels closely resembled those in areas unaffected by fire (Figure 3). However, post-wildfire wind erosion and SDS can be significantly higher than pre-fire rates (Treminio et al., 2024) and have become a growing concern in regions experiencing more frequent and severe wildfires (Strand et al., 2025). For example, extremely large sediment mass fluxes ranging from a three-month average of 2.84 kg m ¹ (Sankey et al., 2009b) to greater than 44.01 kg m^-1^ d^-1^ (Miller et al., 2012) have been recorded in burned areas compared to negligible amounts in unburned sites, with sediment fluxes decreasing as herbaceous cover increased (Morra et al., 2024). We note that the magnitude of modeled *Q* in our study is generally smaller than short-term field measurements of post-wildfire wind erosion because our AERO model predictions were derived from probabilistic 30-year hourly wind speed data that would not capture higher frequency peaks in wind speed that drive sediment transport (Stout, 1998). Research in the Great Basin shows that increased cover of invasive annual cheatgrass is linked to reduced aeolian sediment transport, but cheatgrass also fuels more frequent fires that trigger orders-of-magnitude increases in sediment transport rates post-wildfire (Miller et al., 2012; Treminio et al., 2024; Wagenbrenner et al., 2013), which would continue rangeland susceptibility to long-term wind erosion problems. Consistent with previous studies, our results show that over time, post-wildfire vegetative regrowth, regardless of plant species’ native status, can play a crucial role in mitigating wind erosion (Morra et al., 2024; Roehner et al., 2020; Sankey et al., 2009b, 2009a).

For restoration treatments not influenced by wildfire, we found most treatments along with their respective untreated controls — excluding soil disturbance and aerial seeding controls — resulted in significantly lower Ln *Q*. Among these, closure produced the greatest reduction in aeolian sediment transport, while drill seeding had the least reduction. However, when assessing treatment effects (i.e., treatment minus untreated control), only soil disturbance and aerial seeding treatments showed a significant effect. These reduction in Ln *Q* are likely due to higher perennial grass cover in treatment plots compared to higher annual grass cover in untreated controls. The effectiveness of aerial seeding may be attributed to its application at higher elevation and wetter sites, consistent with findings from Knutson et al. (2014) and Peppin et al. (2010). Moreover, soil disturbance treatments resulted in the second-largest reduction in sediment transport rates. This outcome was somewhat unexpected, as such treatments typically disrupt soil structure and composition, often leaving bare soil exposed and vulnerable to erosion immediately after application. However, soil disturbance treatments create conditions that promote plant diversity (Müller et al., 2014) and plant cover increases over time, which can help mitigate wind erosion (Nash et al., 2021). Alternatively, soil disturbance treatments applied in areas with high levels of clay or loam may result in soil physical crusting following rain events which may reduce susceptibility to wind erosion (Gillette et al 1980).

Post-wildfire rehabilitation plots treated with aerial seeding, closure, herbicide application, ground seeding, or soil disturbance experienced steadily decreasing aeolian sediment transport rates compared to their untreated counterparts. Of these treatments, closure treatments experienced the largest reduction in aeolian sediment transport rates, while seedling planting had the smallest effect. Across all these treatments — except for non-treated ground seeding, drill seeding, and vegetation disturbance plots — Ln *Q* responses were significantly less than zero. This observation suggests that the treatments showed a slightly greater reduction in aeolian sediment transport compared to background levels that might occur in untreated plots. Yet, significant treatment effects (i.e., treatment minus control) only occurred for closure, herbicide application, and ground seeding. As in the restoration plots, increased perennial grass in treated plots compared to increased annual grass cover in non-treated plots for closure areas after post-fire rehabilitation probably led to the observed treatment effect. It is likely that, due to the exclusion of grazing in closure plots, perennial grasses took advantage of post-wildfire nutrient enrichment, which boosts soil fertility in moderate to severe burns (Strain et al., 2024). Furthermore, nutrients released by combustion of vegetation can be redistributed from their source (Roehner et al., 2020; Sankey et al., 2012b, 2012a) and potentially fertilize plant communities downwind. Outside of the closure plots, perennial grasses likely were preferentially grazed, which allowed annual grass to reach higher cover compared to treatments (e.g., Souther et al., 2019). In plots treated with herbicide, the observed reduction in sediment transport was likely attributable to the removal of resource-competing plants, thereby facilitating an increase in perennial plant cover relative to untreated plots (Ehlert et al., 2019). For ground seeding, the reason for a significant treatment effect is less clear, as permutation tests did not reveal substantial differences in plant functional group cover between treated and untreated plots. Our interpretation is that a change in vegetation height associated with small changes in cover of annual forb (1.46%), perennial grass (0.271%), and sagebrush (0.506%) may have contributed to the observed effect.

In this study, we applied propensity score matching (PSM) with model-estimated counterfactual treatment effects to evaluate how dryland restoration and rehabilitation treatments influence potential wind erosion. By maintaining a consistent set of covariates across both PSM and multiple linear regression analyses, we ensured that the causal impact of the treatments could be assessed at a regional scale. While individual effects of covariates on treatment responses remain challenging to interpret (Westreich and Greenland, 2013), the approach presented in our analyses provides a valuable opportunity for refining methodologies that assess wind erosion and SDS dynamics. The covariates — capturing plant cover attributes that influence wind erosion — offer a foundation for expanding studies that compare burned and unburned locations, as well as restoration and rehabilitation treatments, across diverse environmental contexts. Future work incorporating larger sample sizes and mixed-model statistical approaches presents an opportunity to examine how specific factors, such as soil texture, influence treatment outcomes and interact with restoration strategies, ultimately advancing the applicability of treatments and their effect on wind erosion mitigation.

Our use of multiple linear regressions, driven by sample size constraints, provides an opportunity to explore advanced modeling approaches in future studies. While mixed-model frameworks accounting for random effects can enhance population-level inferences, integrating these methods in future research could improve our understanding of group-level heterogeneity. For example, addressing temporal and spatial variability through expanded datasets may allow for the incorporation of time since treatment and preceding treatment effects as variables of interest. With larger sample sizes, within-region variability could be modeled as random effects, contributing to broader generalizability. Additionally, leveraging remote sensing and gridded data presents exciting possibilities for strengthening observational insights (e.g., Fick et al., 2021; McNellis et al., 2023; Nauman et al., 2017; Simler-Williamson and Germino, 2022). Future research could explore interactions among multiple wildfires and treatments, particularly in areas with sufficient sampling, to advance our ability to link fine-scale vegetation treatment responses to wind erosion and SDS. Incorporating mixed-model frameworks would also allow for a deeper investigation of covariates like soil texture, improving our ability to identify the mechanisms driving treatment outcomes. By building upon the connections between standardized observational monitoring data, wind erosion and SDS estimates, and statistical analyses, new studies could further integrate spatial and temporal dynamics, broadening the applicability of findings to restoration and post-disturbance rehabilitation and their effect on wind erosion and SDS mitigation.

## 5. Conclusion

This study addresses the critical challenge of integrating wind erosion and SDS mitigation into dryland restoration and rehabilitation monitoring efforts with a specific focus on rangelands in the Great Basin. Our findings demonstrated that various treatment types, including aerial seeding, ground seeding, soil disturbance, and closure significantly reduced predicted aeolian sediment transport rates, highlighting their effectiveness in mitigating wind erosion risk. Enhanced plant cover, in part driven by differences in perennial grass cover and reduced bare ground exposure, are key factors driving these positive outcomes, supporting our hypothesis that vegetative cover responses to restoration and rehabilitation treatments moderate aeolian sediment transport rates. These results provide empirical evidence for incorporating wind erosion and SDS mitigation into restoration initiatives and underscore the importance of promoting resilient plant communities to combat land degradation. This study introduces innovative applications of propensity score matching, the AERO model, standardized monitoring data, and treatment information to advance regional-scale assessments of restoration and rehabilitation outcomes and their effect on wind erosion and SDS. Ultimately, these findings equip land managers and policymakers with actionable insights to optimize restoration strategies for reducing wind erosion and SDS risks across vulnerable landscapes.

Finally, our study contributes to the body of evidence showing potential benefits of dryland restoration and post-wildfire rehabilitation for reducing wind erosion — a key symptom and driver of land degradation. Over the past 75 years, numerous dryland restoration and post-wildfire rehabilitation treatments in the Great Basin have been implemented to address soil stability, invasive species management, improve wildlife habitat, and improve the quality and quantity of livestock forage. The success of these efforts in general, and specifically for increasing cover of native vegetation, has been variable relative to treatment objectives. The effectiveness of restoration treatments in mitigating wind erosion depends primarily on how they modify vegetation cover and bare ground exposure. Any treatment that successfully increases the spatial and temporal extent of plant cover and its aerodynamic sheltering will tend to reduce wind erosion. While restoration and rehabilitation treatments may not always bring back historical plant communities, our results show that it is feasible to reduce wind erosion and SDS risk at the regional scale — a key step toward restoring ecosystem function, reversing land degradation, and improving air quality.

## Acknowledgements

This research was funded by the United States National Science Foundation, the United Kingdom Research and Innovation Natural Environment Research Council Grant # 1853853, the United States Department of Agriculture — Natural Resources Conservation Service — Conservation Effects Assessment Project (CEAP) Grazing Lands Agreement # 60-3050-1-003, and the Bureau of Land Management Agreement # 4500104319. We are sincerely grateful to Kristina Young for conceptual discussion of treatment effects and Michelle Jeffries for guidance with Land Treatment Digital Library data. This research was a contribution of the Long-Term Agro-ecosystem Research (LTAR) network supported by the U.S. Department of Agriculture (USDA). Any use of trade, product, or firm names is for descriptive purposes only and does not imply endorsement by the U.S. Government.

## Data availability statement

Data for AERO and Bureau of Land Management AIM Program vegetation indicators can be found at https://landscapedatacommons.org (McCord et al., 2023). Land treatment digital library data can be found at https://www.usgs.gov/apps/ltdl/ (Pilliod et al., 2019). Wildfire data can be found at https://doi.org/10.5066/P9Z2VVRT (Welty and Jeffries 2020).

