## Supplemental Figures for "Wildfire, restoration, and post-wildfire rehabilitation effects on wind erosion in the Great Basin"

**Table S1.** Data source and variable descriptions used for propensity score matching and analyzing the effect of rangeland treatments on aeolian sediment transport in the Great Basin. Abbreviation are AIM: Assessment, Inventory, and Monitoring; LTDL: Land Treatment Digital Library; WFTT: Wildland Fire Trends Tool; SOLUS: Soil Landscaped of the United States; DEM: Digital elevation model; PRISM: PRISM Climate Group; USDA: United States Department of Agriculture.

| **Source** | **Variable** | **Description** |
| --- | --- | --- |
| AERO | *lnQ* | Natural-log transformed *Q* prediction from AERO model |
| AIM | *PrimaryKey* | PrimaryKey ID |
|  | *plotkey* | plotkey ID |
|  | *latitude* | Latitude for AIM plot location |
|  | *longitude* | Longitude for AIM plot location |
|  | *samp_yr* | The year an AIM plot was sampled |
|  | *BareSoilCover* | Percentage of bare soil cover measured at an AIM plot |
|  | *TotalFoliarCover* | Percentage of total foliar cover of measured at an AIM plot |
|  | *AH_ForbCover* | Percentage of forb cover of measured at an AIM plot |
|  | *AH_GrassCover* | Percentage of grass cover measured at an AIM plot |
|  | *AH_ShrubCover* | Percentage of shrub cover measured at an AIM plot |
|  | *gap_gt1m* | Percentage of an AIM plot with canopy gaps greater than 1 meter |
|  | *AH_AnnualForbCover* | Percentage of annual forb cover measured at an AIM plot |
|  | *AH_AnnualGrassCover* | Percentage of annual grass cover measured at an AIM plot |
|  | *AH_PerennialForbCover* | Percentage of perennial forb cover measured at an AIM plot |
|  | *AH_PerennialGrassCover* | Percentage of perennial forb cover measured at an AIM plot |
|  | *AH_Sagebrush* | Percentage of sagebrush species cover measured at an AIM plot |
| LTDL | *Trt_Type_S* | Major treatment designation |
| SOLUS | *sand_pct_solus* | Percentage of sand predicted at an AIM plot |
| DEM | *elev_m* | Estimated elevation of AIM plot location |
| PRISM | *annual_ppt_mm* | Estimated mean annual precipitation in mm |
| USDA | *mlra* | Identity of Major Land Resource Area an AIM monitoring plot occurs in |

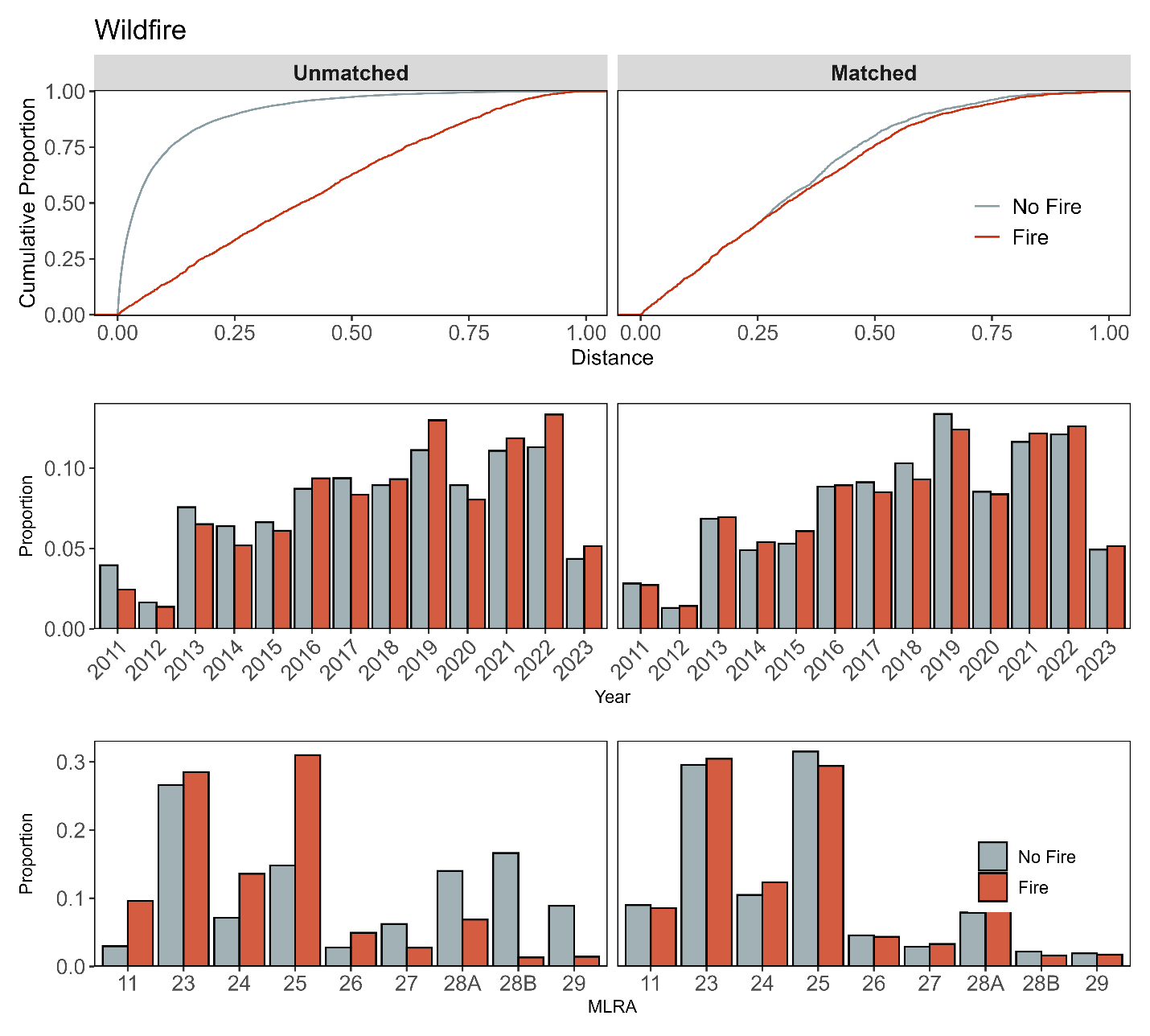

**Figure S1a**. Assessment of covariate balance using propensity score matching (PSM) for No treatment effect monitoring plots. The top plot shows the empirical cumulative distribution function (eCDF) for the distance measure of the propensity score. The middle and bottom plots display balance in categorical variables, accounting for the year of monitoring and the major land resource area (MLRA) of the plots, comparing unmatched (left column) and matched sets (right columns) of monitoring plots, both with and without a history of fire.

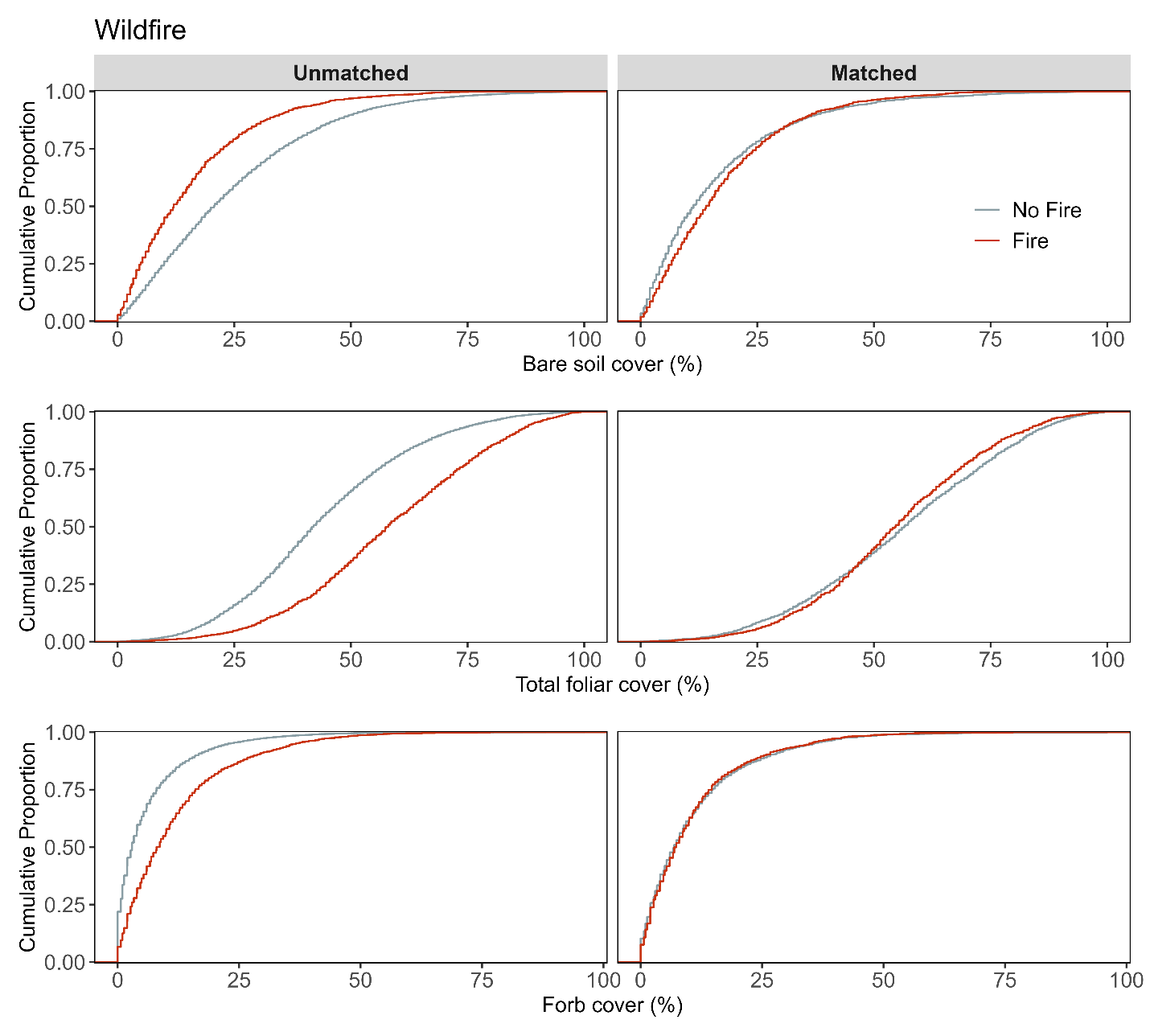

**Figure S1b.** Assessment of covariate balance using PSM for No treatment effect monitoring plots. The figure compares eCDFs for bare soil (top), total foliar cover (middle), and forb cover (bottom) percentages between plots with and without a history of fire, for both unmatched (left column) and matched sets of plots (right column).

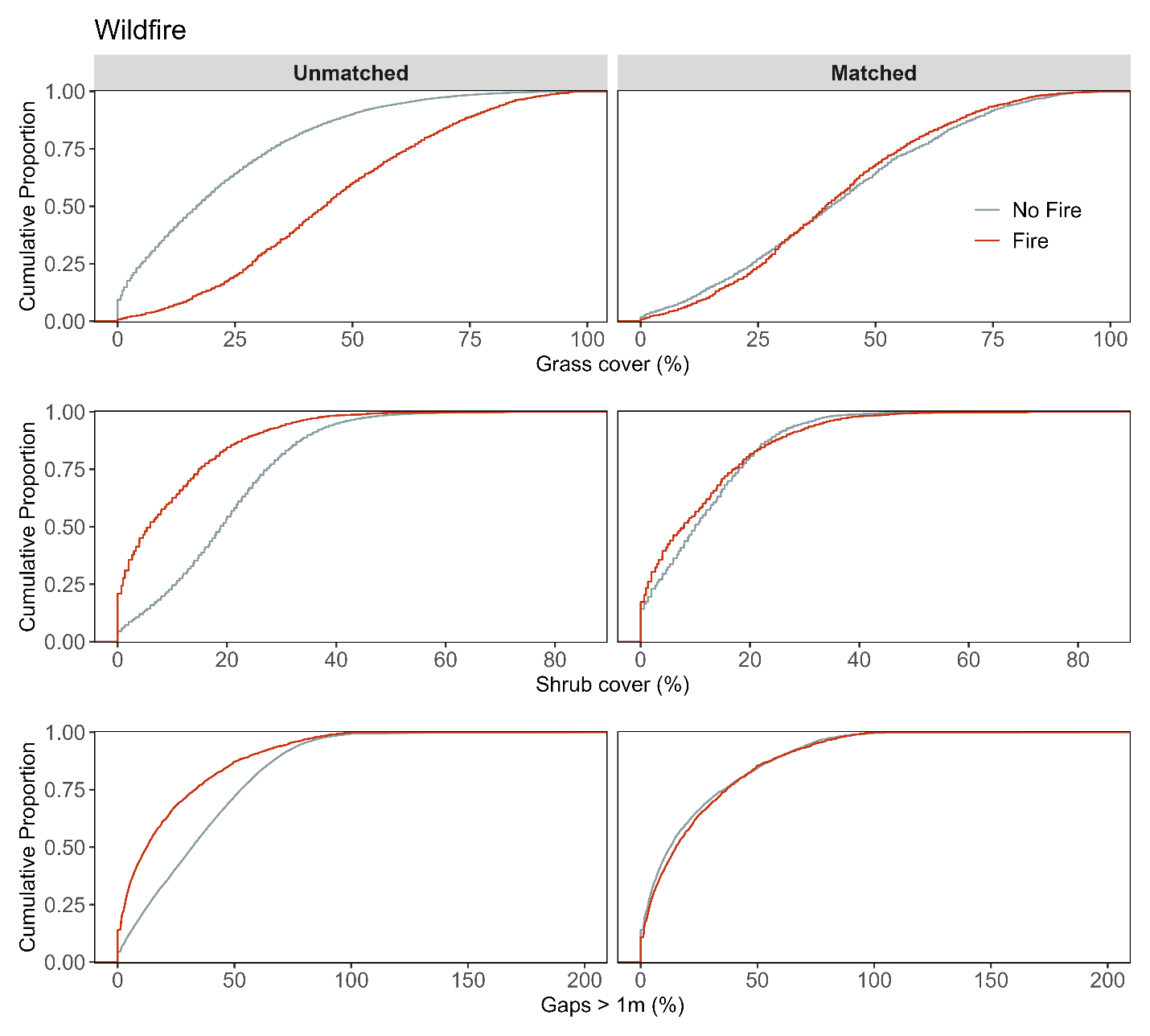

**Figure S1c.** Assessment of covariate balance using PSM for No treatment effect monitoring plots. The figure compares eCDFs for grass cover (top), shrub cover (middle), and gaps > 1m (bottom) percentages between plots with and without a history of fire, for both unmatched (left column) and matched sets of plots (right column).

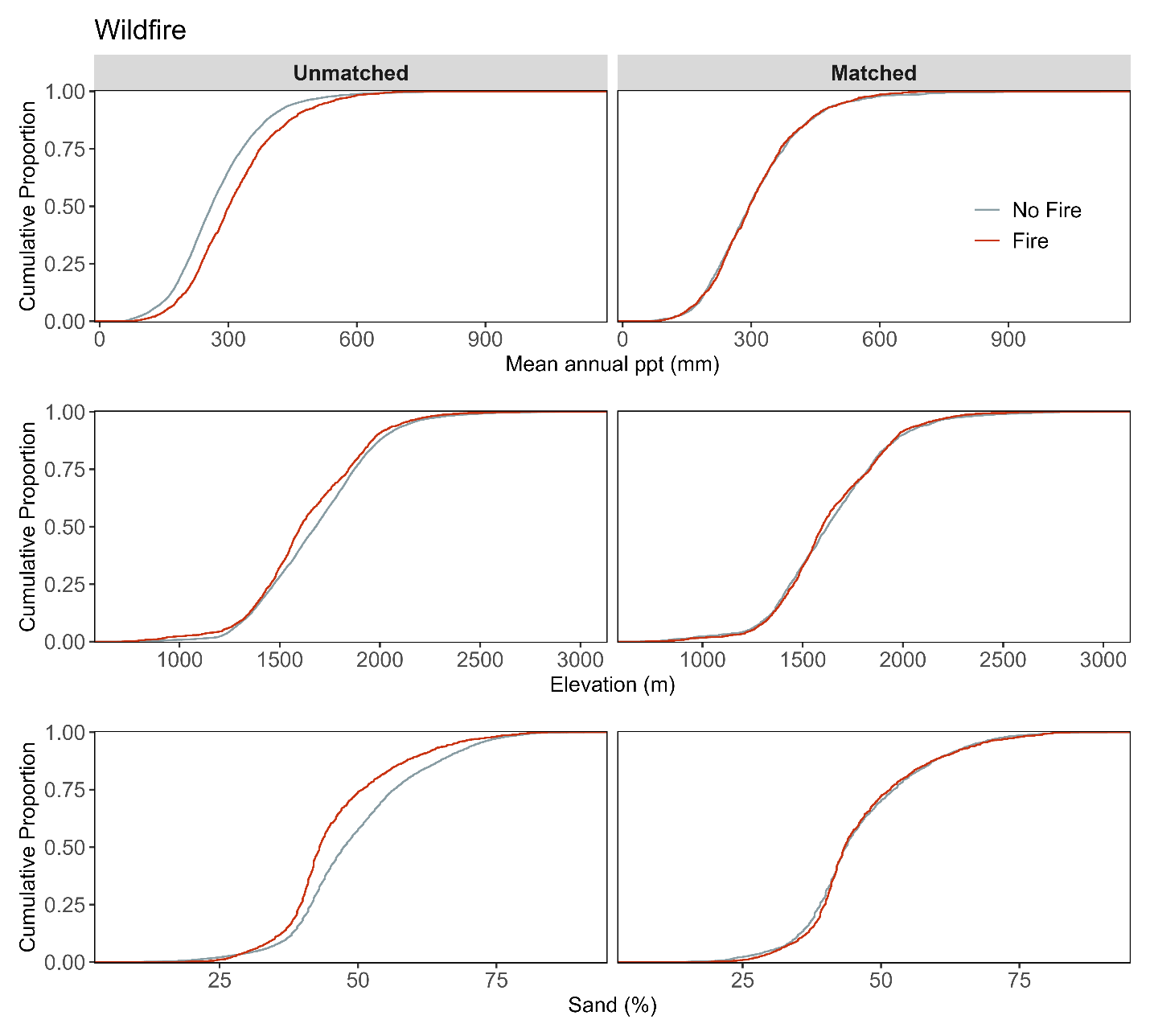

**Figure S1d**. Assessment of covariate balance using PSM for No treatment effect monitoring plots. The figure compares eCDFs for mean annual precipitation (top), elevation (middle), and sand texture (bottom) percentages between plots with and without a history of fire, for both unmatched (left column) and matched sets of plots (right column).

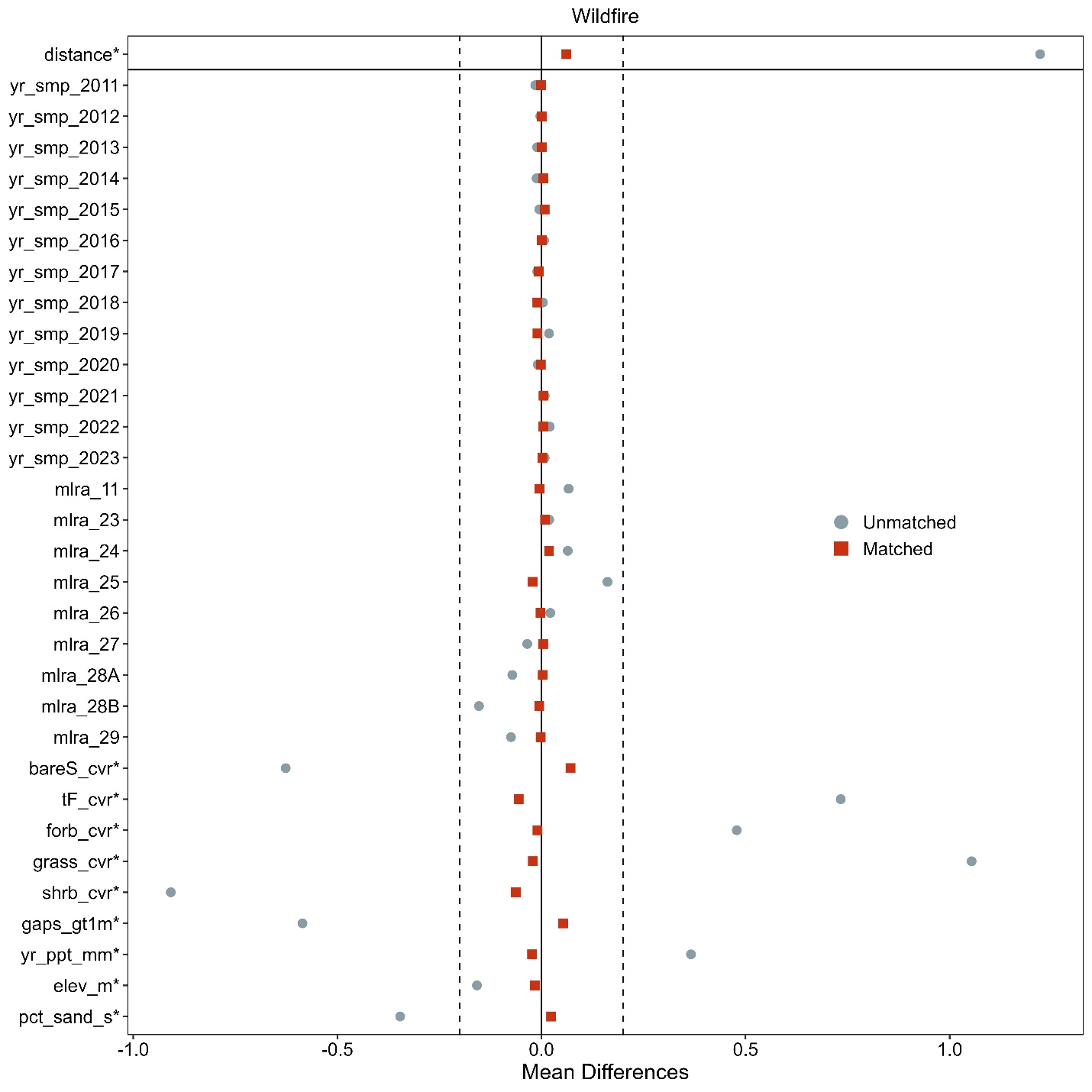

**Figure S1e**. Love-plot illustrating standardized mean differences for covariates in both the full dataset (unmatched) and the matched dataset of No treatment effect plots. Vertical lines indicate the caliper threshold (0.2 times the standard deviation of the propensity score) applied during matching.

**Supporting Information for Restoration monitoring plots and non-treated controls**

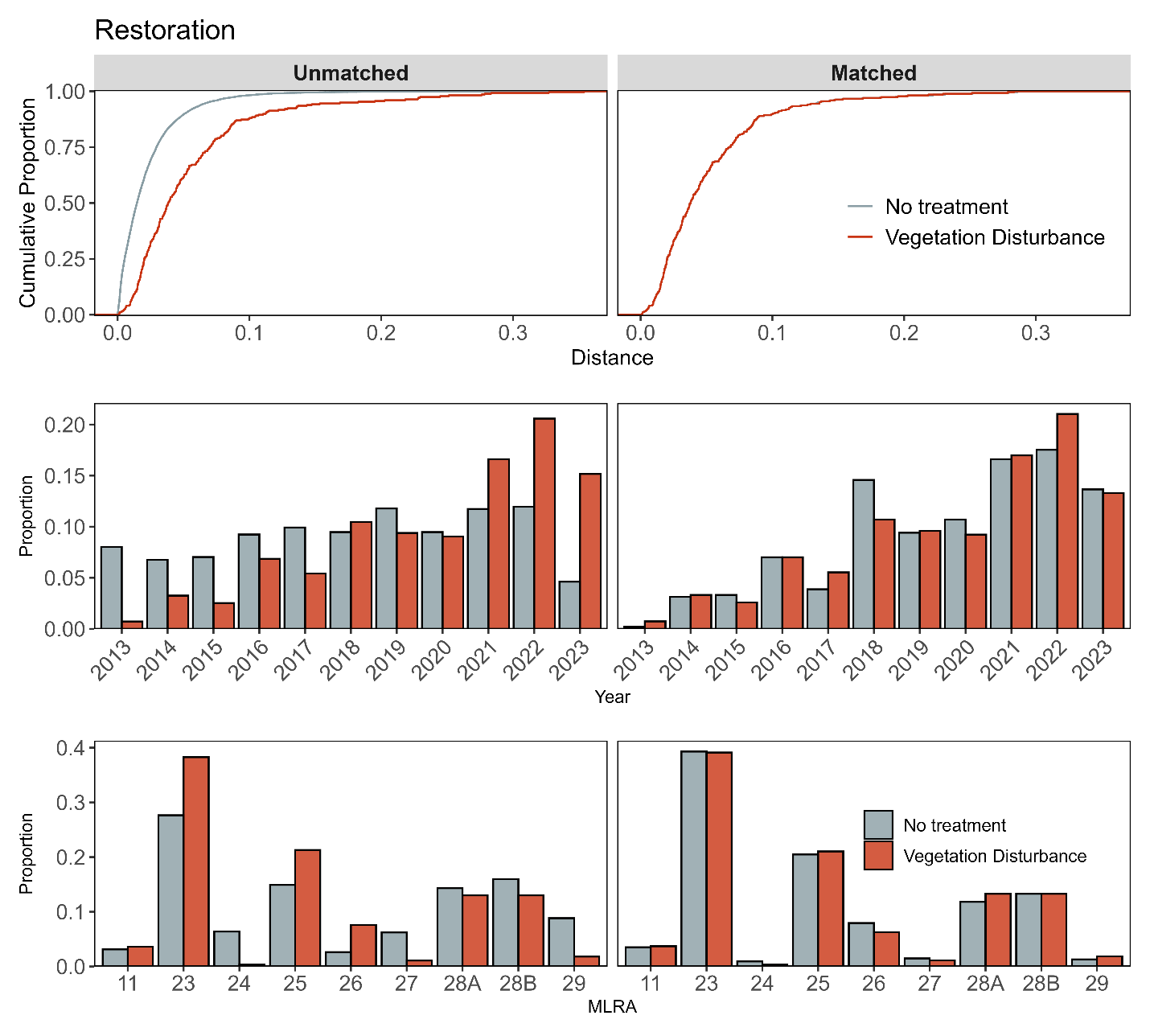

**Figure S2a.** Assessment of covariate balance using propensity score matching (PSM) for Restoration monitoring plots. The top plot shows the empirical cumulative distribution function (eCDF) for the distance measure of the propensity score. The middle and bottom plots display balance in categorical variables, accounting for the year of monitoring and the major land resource area (MLRA) of the plots, comparing unmatched (left column) and matched sets (right columns) for vegetation disturbance and non-treated control monitoring plots.

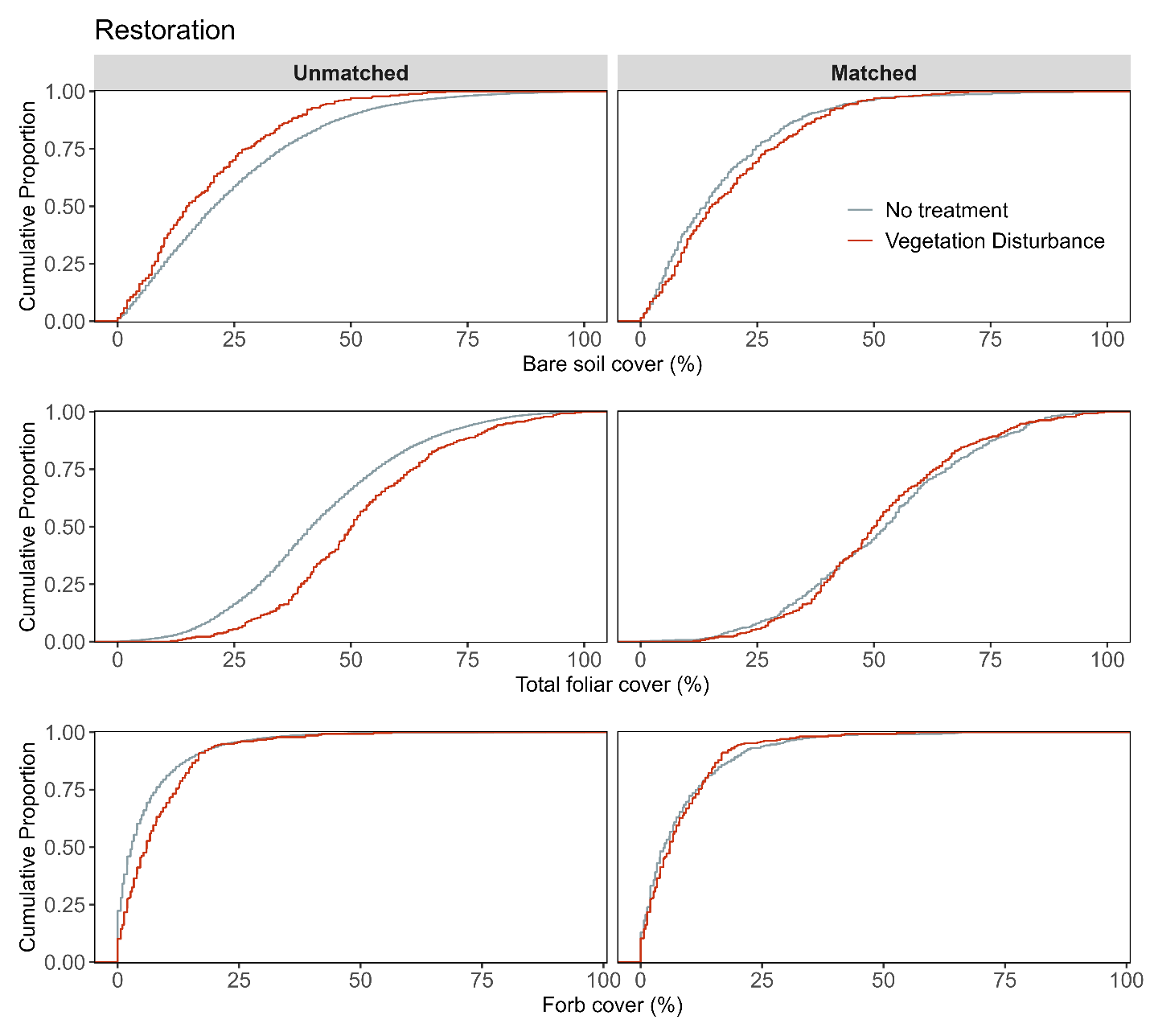

**Figure S2b.** Assessment of covariate balance using PSM for Restoration monitoring plots. The figure compares eCDFs for bare soil (top), total foliar cover (middle), and forb cover (bottom) percentages between vegetation disturbance treatments and non-treated controls, for both unmatched (left column) and matched sets of monitoring plots (right column).

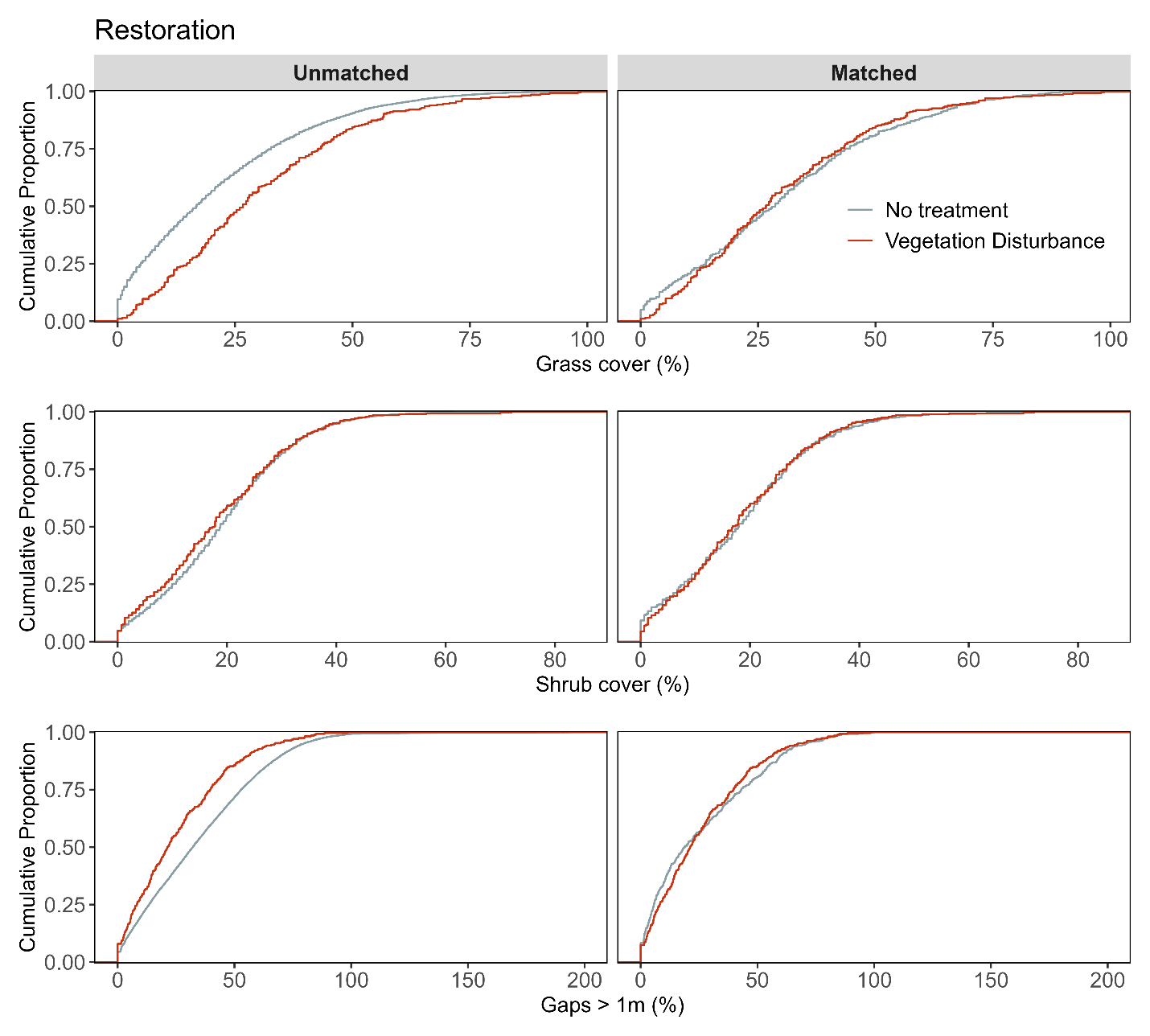

**Figure S2c**. Assessment of covariate balance using PSM for Restoration monitoring plots. The figure compares eCDFs for grass cover (top), shrub cover (middle), and gaps > 1m (bottom) percentages between vegetation disturbance treatments and non-treated controls, for both unmatched (left column) and matched sets of monitoring plots (right column).

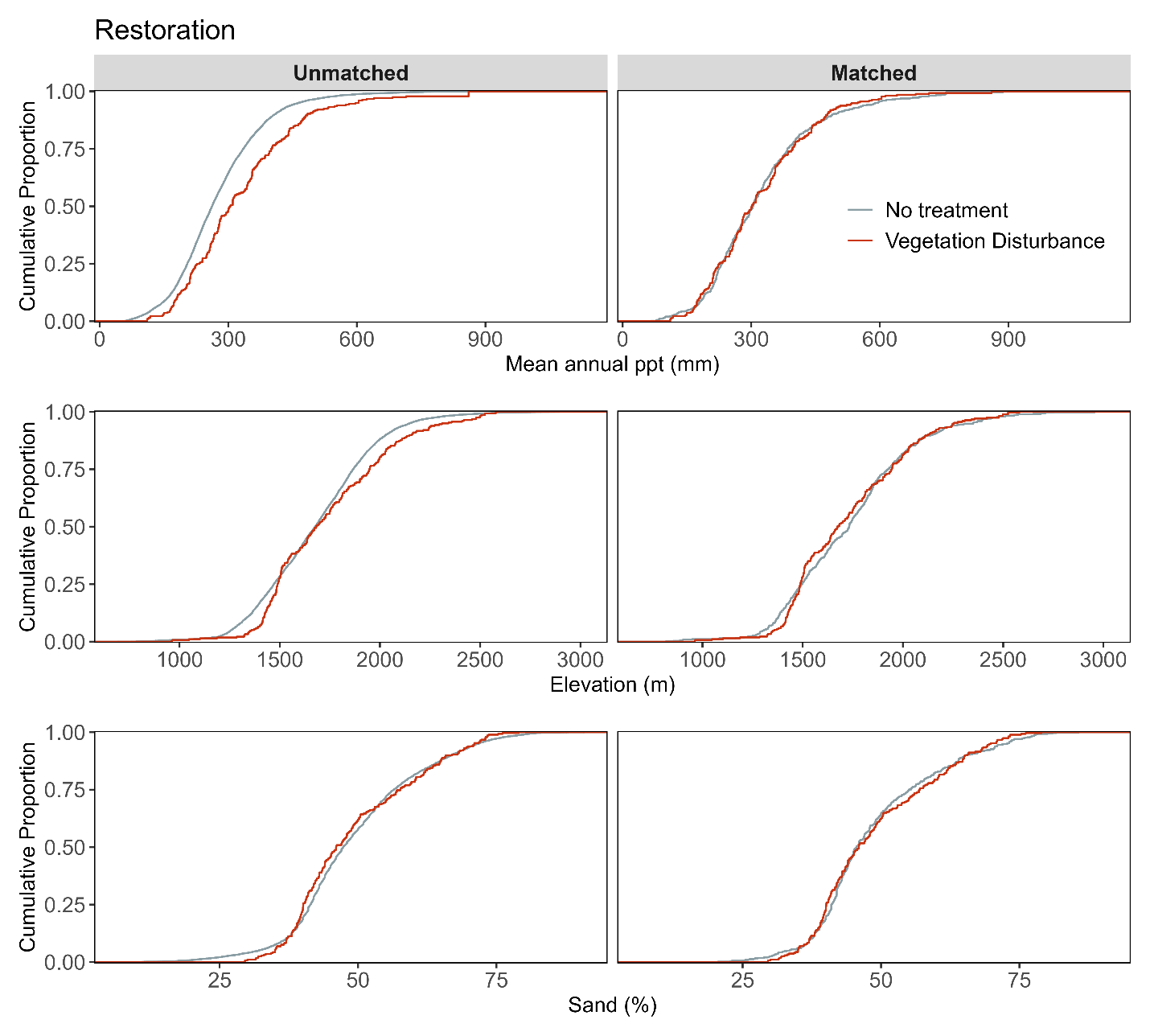
 **Figure Sd.** Assessment of covariate balance using PSM for Restoration effect monitoring plots. The figure compares eCDFs for mean annual precipitation (top), elevation (middle), and sand texture (bottom) percentages between vegetation disturbance treatments and non-treated controls, for both unmatched (left column) and matched sets of monitoring plots (right column).

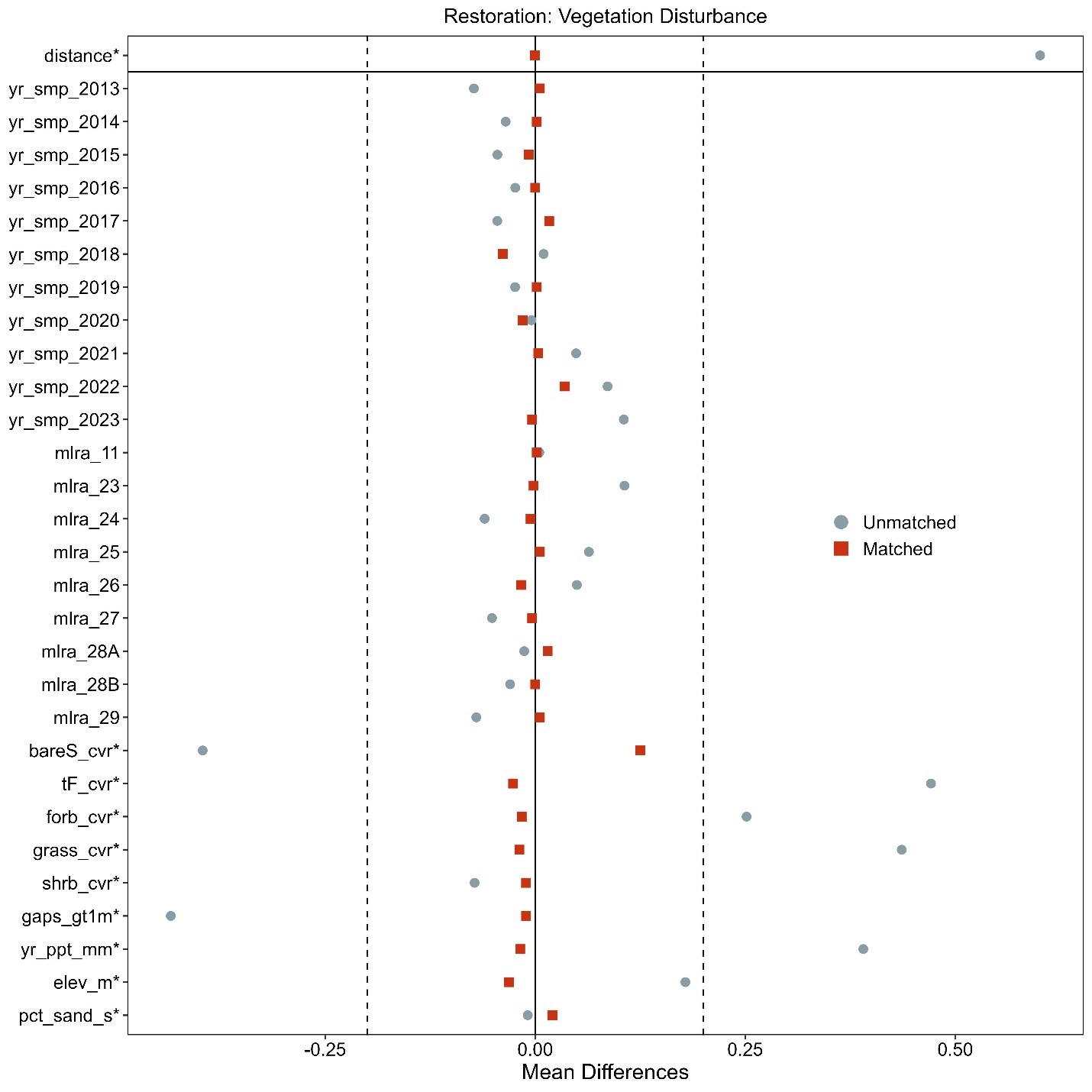

**Figure S4.2e.** Love-plot illustrating standardized mean differences for covariates in both the full dataset (unmatched) and the matched dataset for vegetation disturbance treatments and non-treated controls Restoration plots. Vertical lines indicate the caliper threshold (0.2 times the standard deviation of the propensity score) applied during matching.

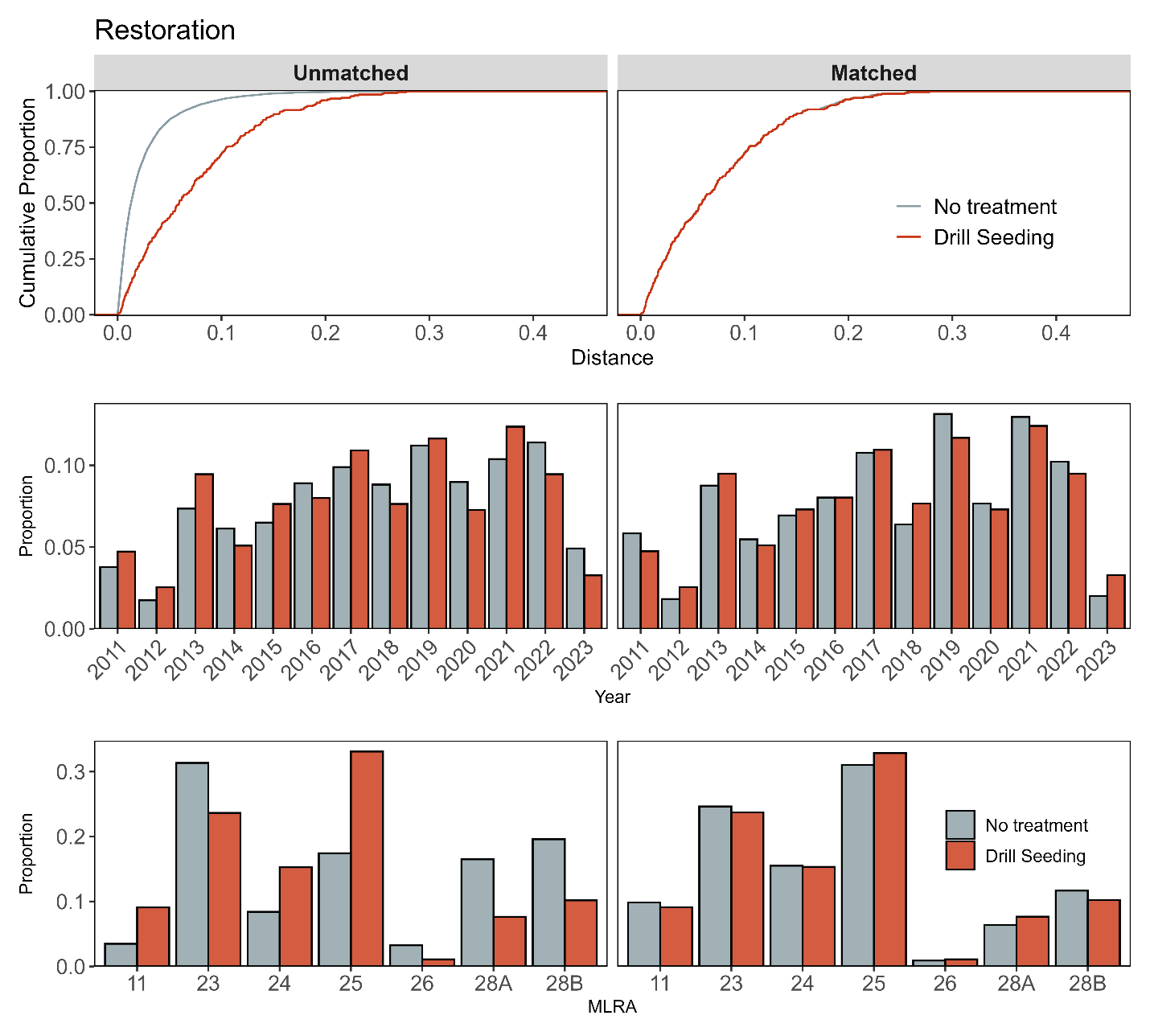

**Figure S3a.** Assessment of covariate balance using propensity score matching (PSM) for Restoration monitoring plots. The top plot shows the empirical cumulative distribution function (eCDF) for the distance measure of the propensity score. The middle and bottom plots display balance in categorical variables, accounting for the year of monitoring and the major land resource area (MLRA) of the plots, comparing unmatched (left column) and matched sets (right columns) for drill seeding treatments and non-treated control monitoring plots.

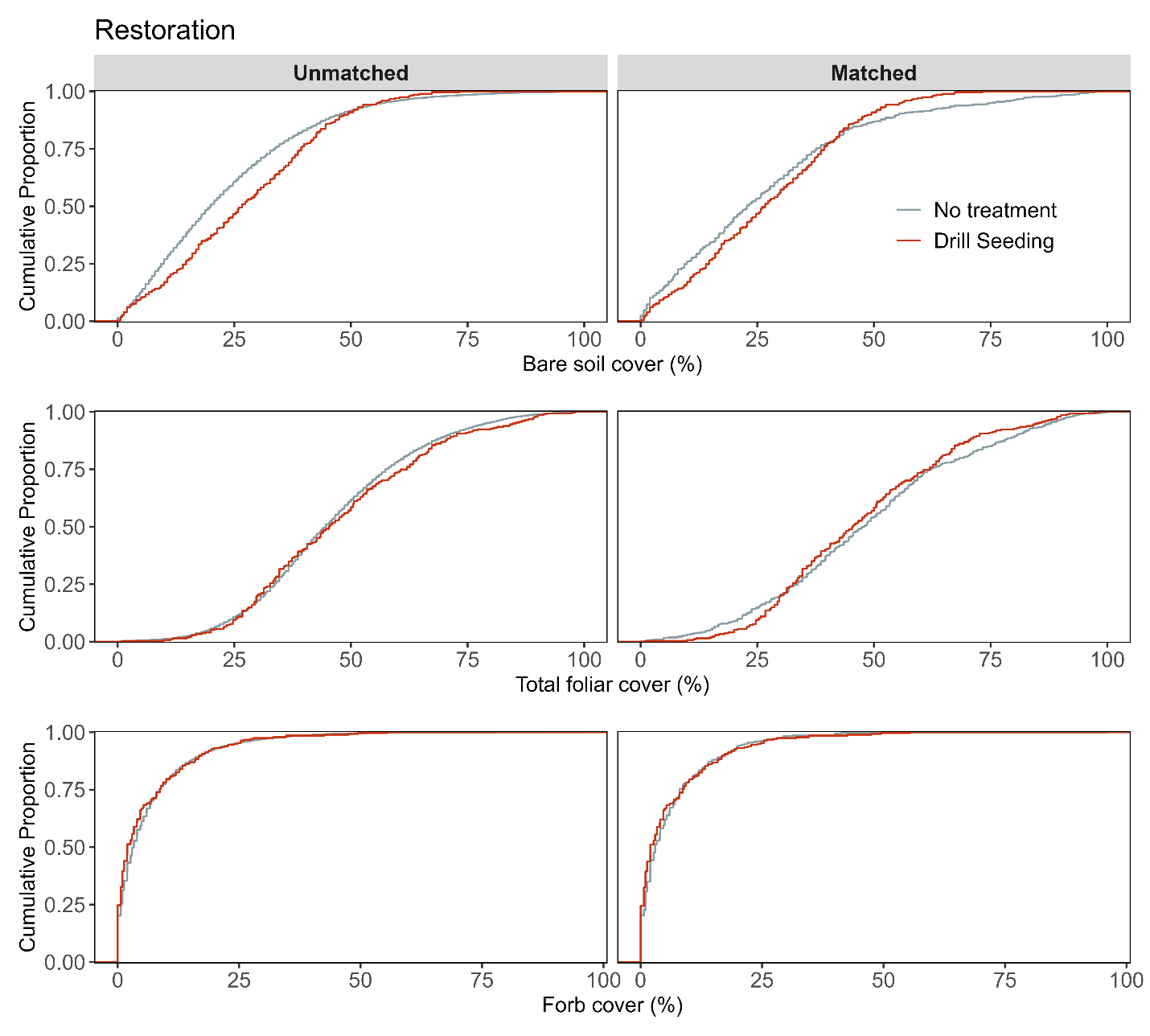

**Figure S3b.** Assessment of covariate balance using PSM for Restoration monitoring plots. The figure compares eCDFs for bare soil (top), total foliar cover (middle), and forb cover (bottom) percentages between drill seeding treatments and non-treated controls, for both unmatched (left column) and matched sets of monitoring plots (right column).

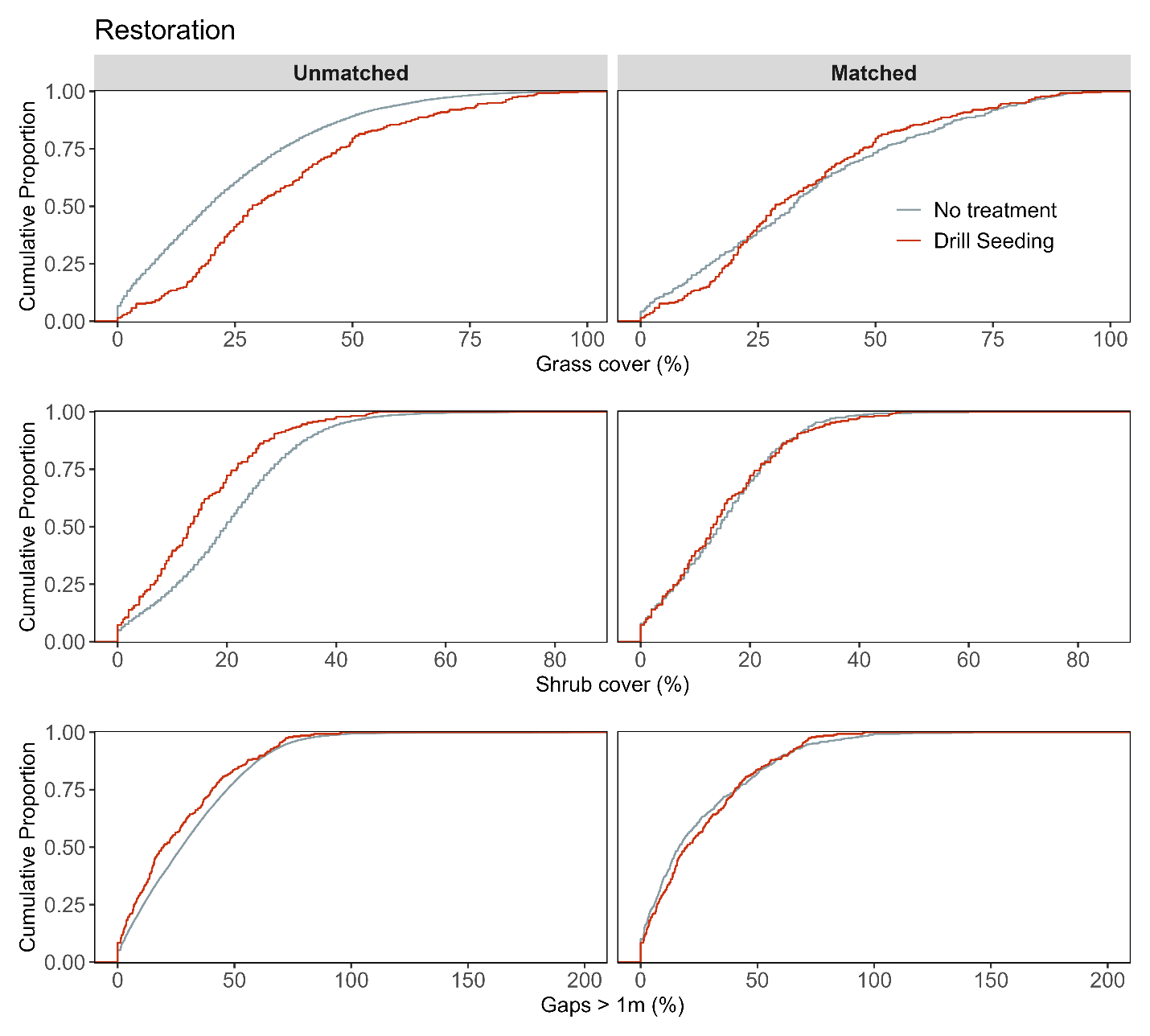

**Figure S3c.** Assessment of covariate balance using PSM for Restoration monitoring plots. The figure compares eCDFs for grass cover (top), shrub cover (middle), and gaps > 1m (bottom) percentages between drill seeding treatments and non-treated controls, for both unmatched (left column) and matched sets of monitoring plots (right column).

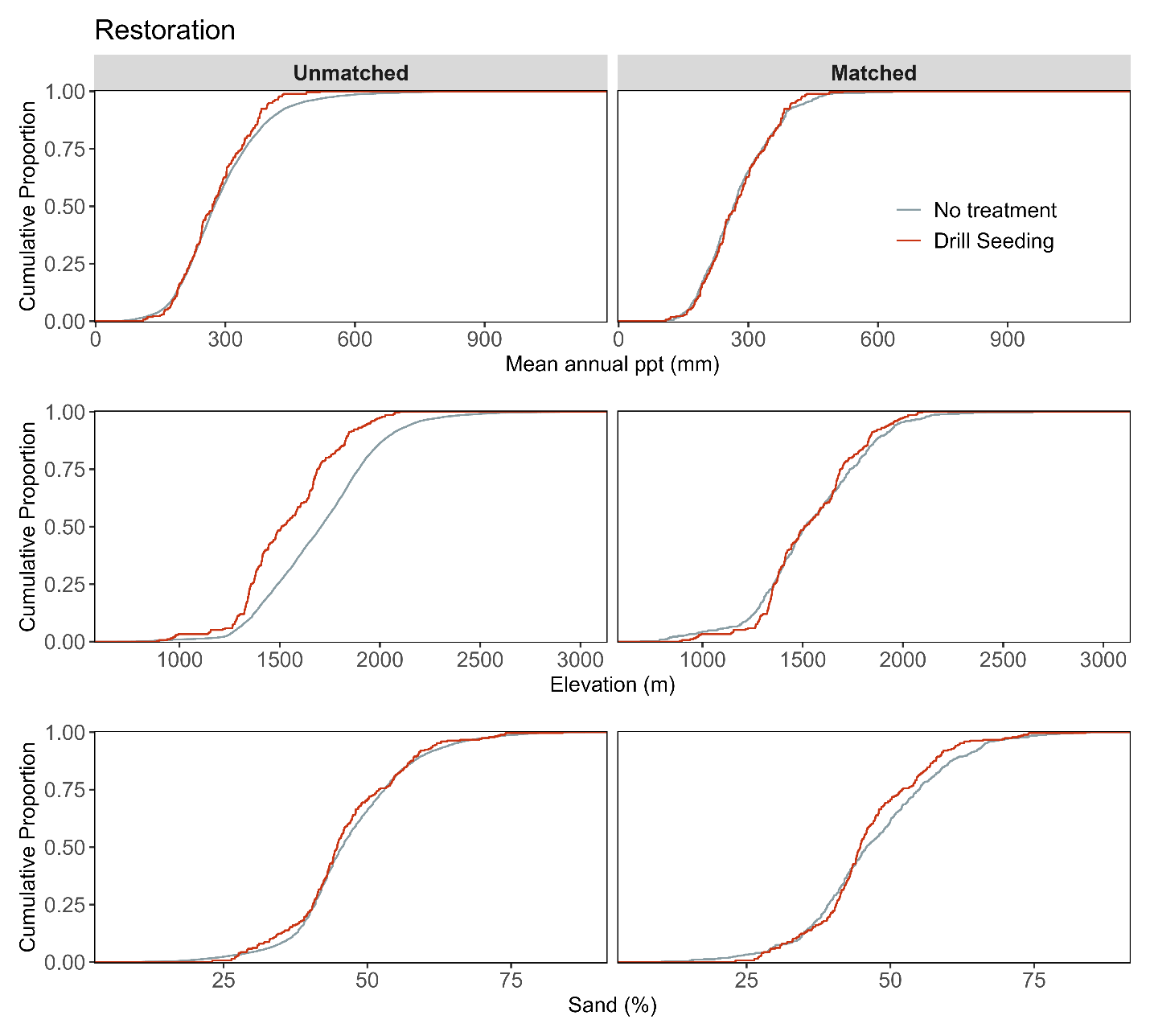

**Figure S3d.** Assessment of covariate balance using PSM for Restoration monitoring plots. The figure compares eCDFs for mean annual precipitation (top), elevation (middle), and sand texture (bottom) percentages between drill seeding treatments and non-treated controls, for both unmatched (left column) and matched sets of monitoring plots (right column).

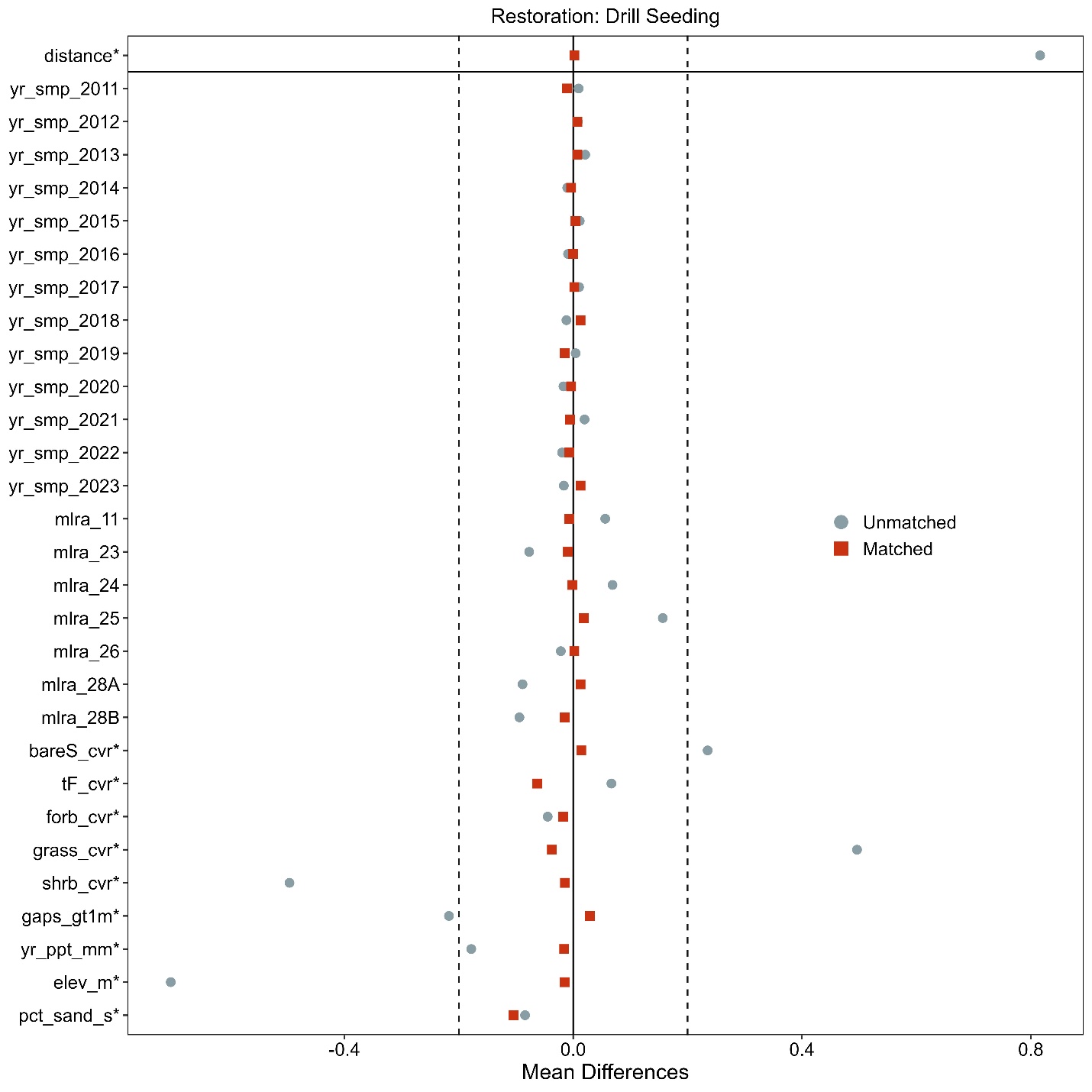

**Figure S3e.** Love-plot illustrating standardized mean differences for covariates in both the full dataset (unmatched) and the matched dataset for drill seeding treatments and non-treated controls Restoration plots. Vertical lines indicate the caliper threshold (0.2 times the standard deviation of the propensity score) applied during matching.

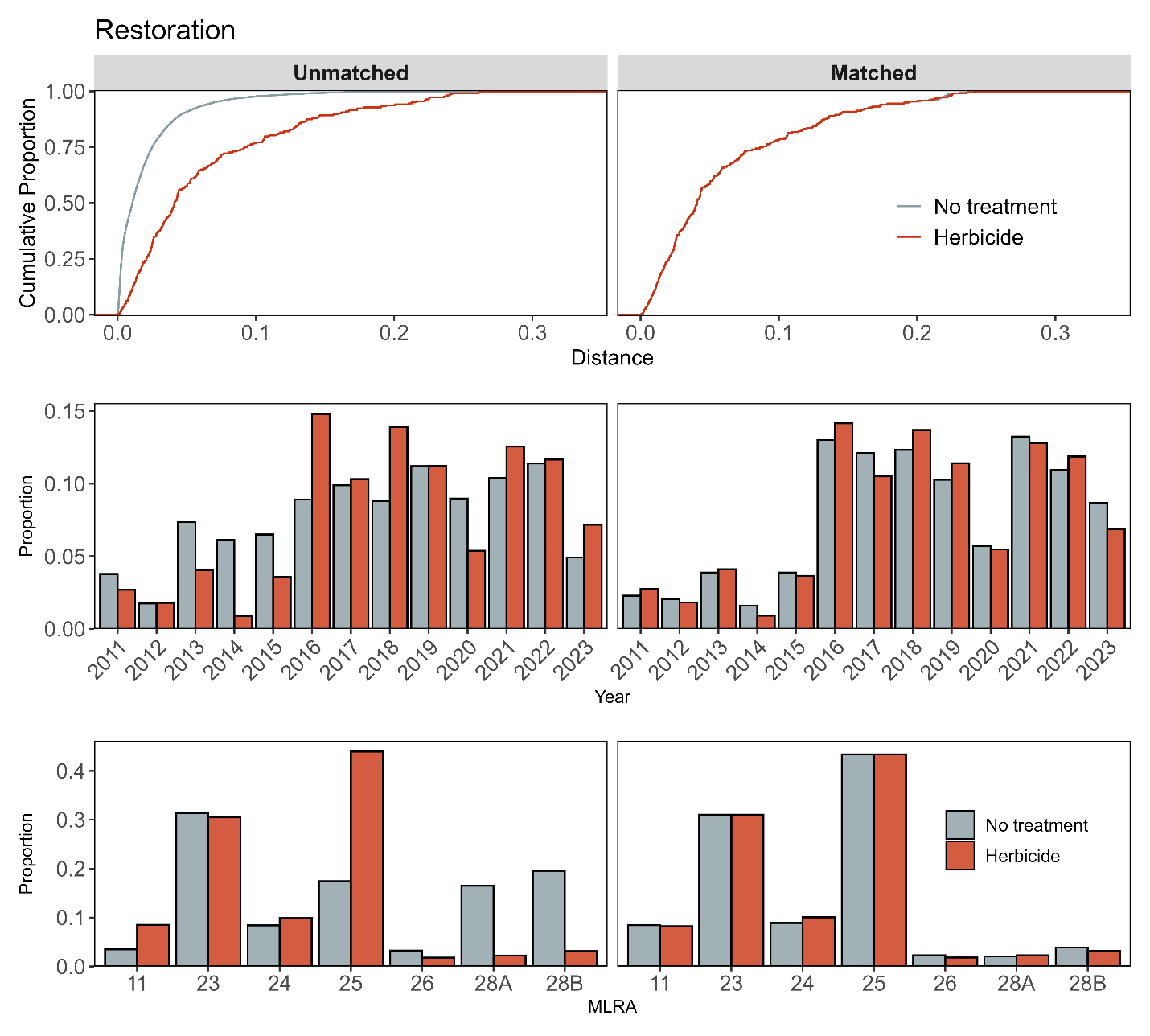

**Figure S4a**. Assessment of covariate balance using propensity score matching (PSM) for Restoration monitoring plots. The top plot shows the empirical cumulative distribution function (eCDF) for the distance measure of the propensity score. The middle and bottom plots display balance in categorical variables, accounting for the year of monitoring and the major land resource area (MLRA) of the plots, comparing unmatched (left column) and matched sets (right columns) for herbicide and non-treated control monitoring plots.

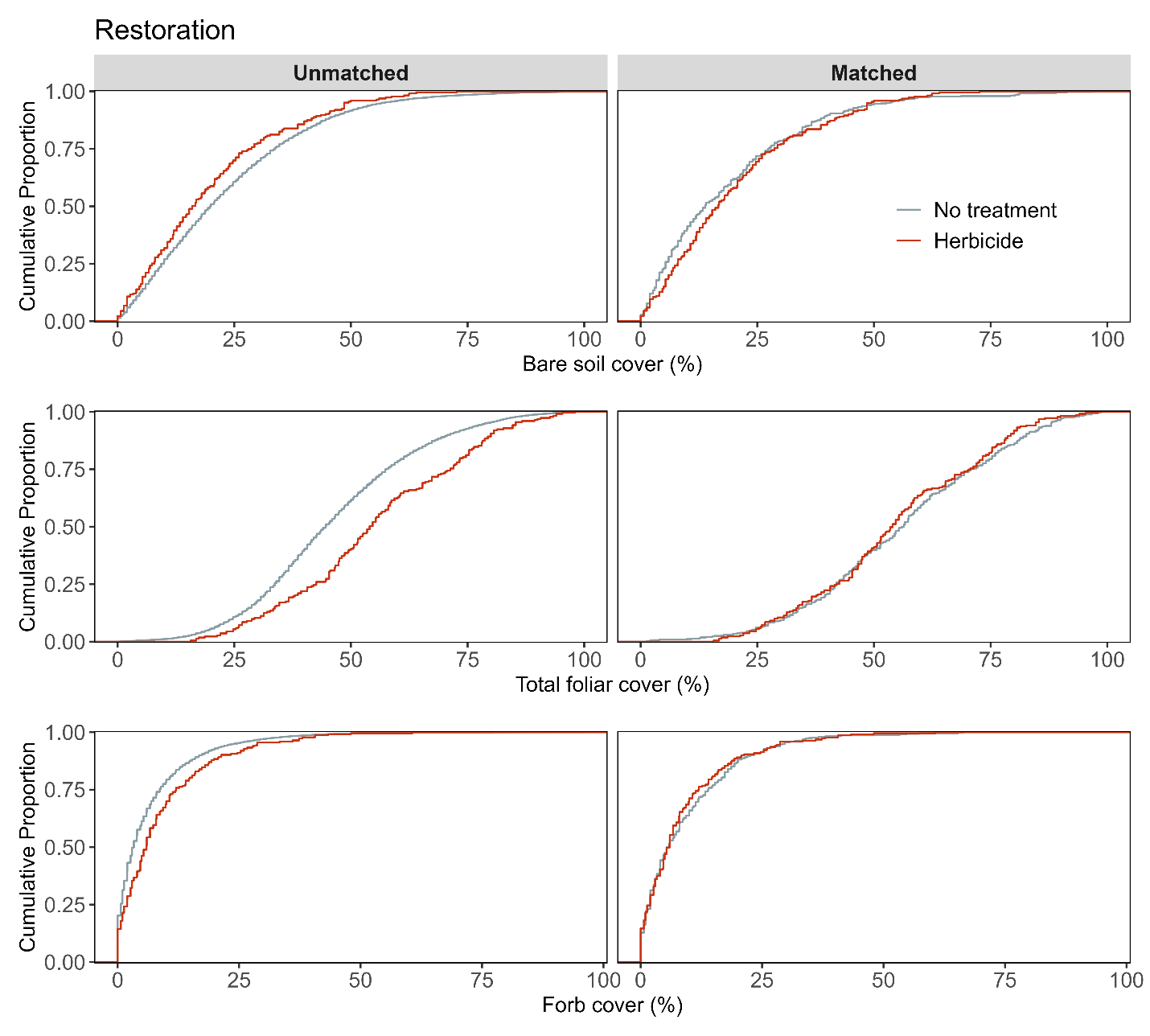

**Figure S4b.** Assessment of covariate balance using PSM for Restoration monitoring plots. The figure compares eCDFs for bare soil (top), total foliar cover (middle), and forb cover (bottom) percentages between herbicide and non-treated controls, for both unmatched (left column) and matched sets of monitoring plots (right column).

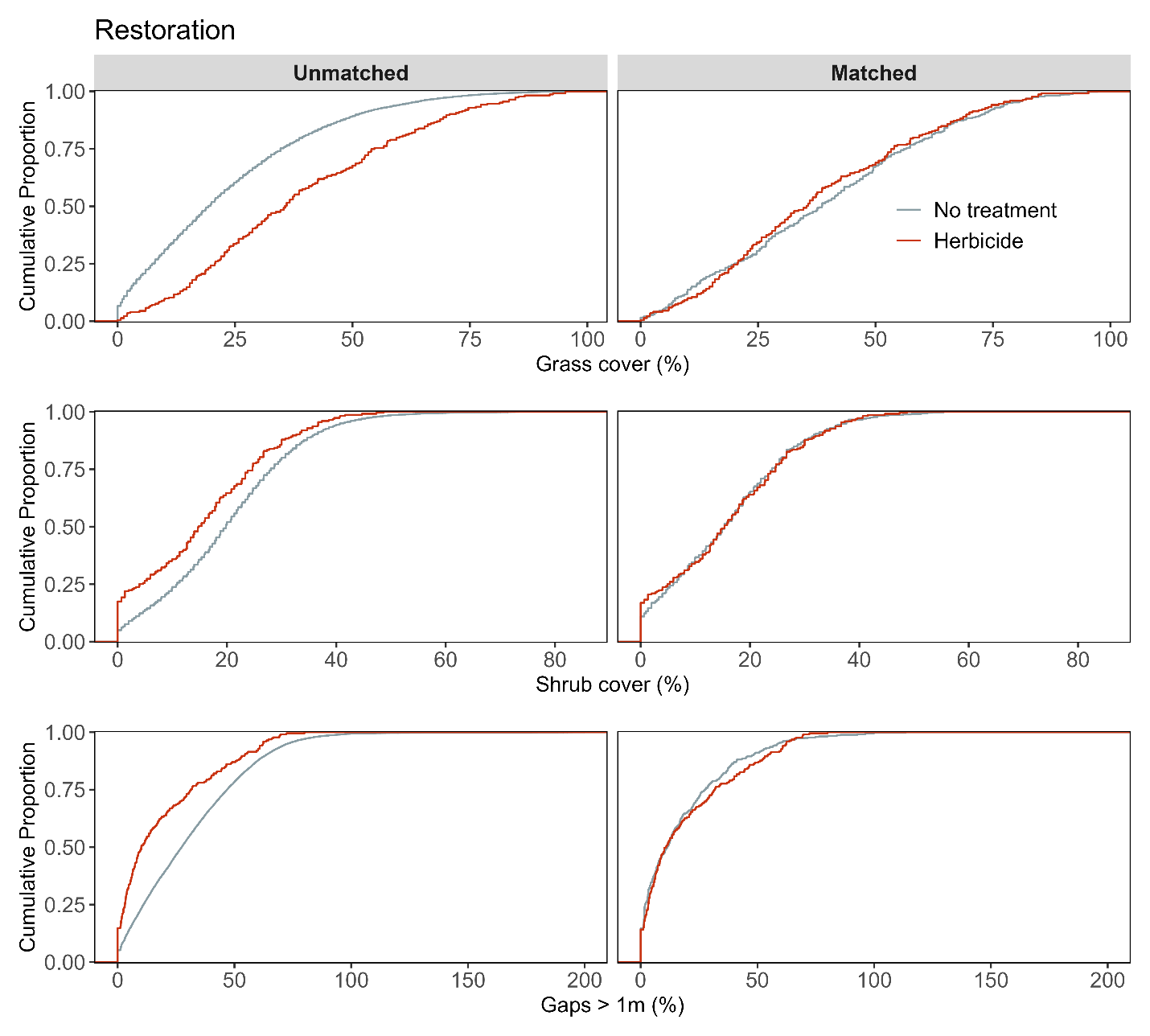

**Figure S4c.** Assessment of covariate balance using PSM for Restoration monitoring plots. The figure compares eCDFs for grass cover (top), shrub cover (middle), and gaps > 1m (bottom) percentages between herbicide and non-treated controls, for both unmatched (left column) and matched sets of monitoring plots (right column).

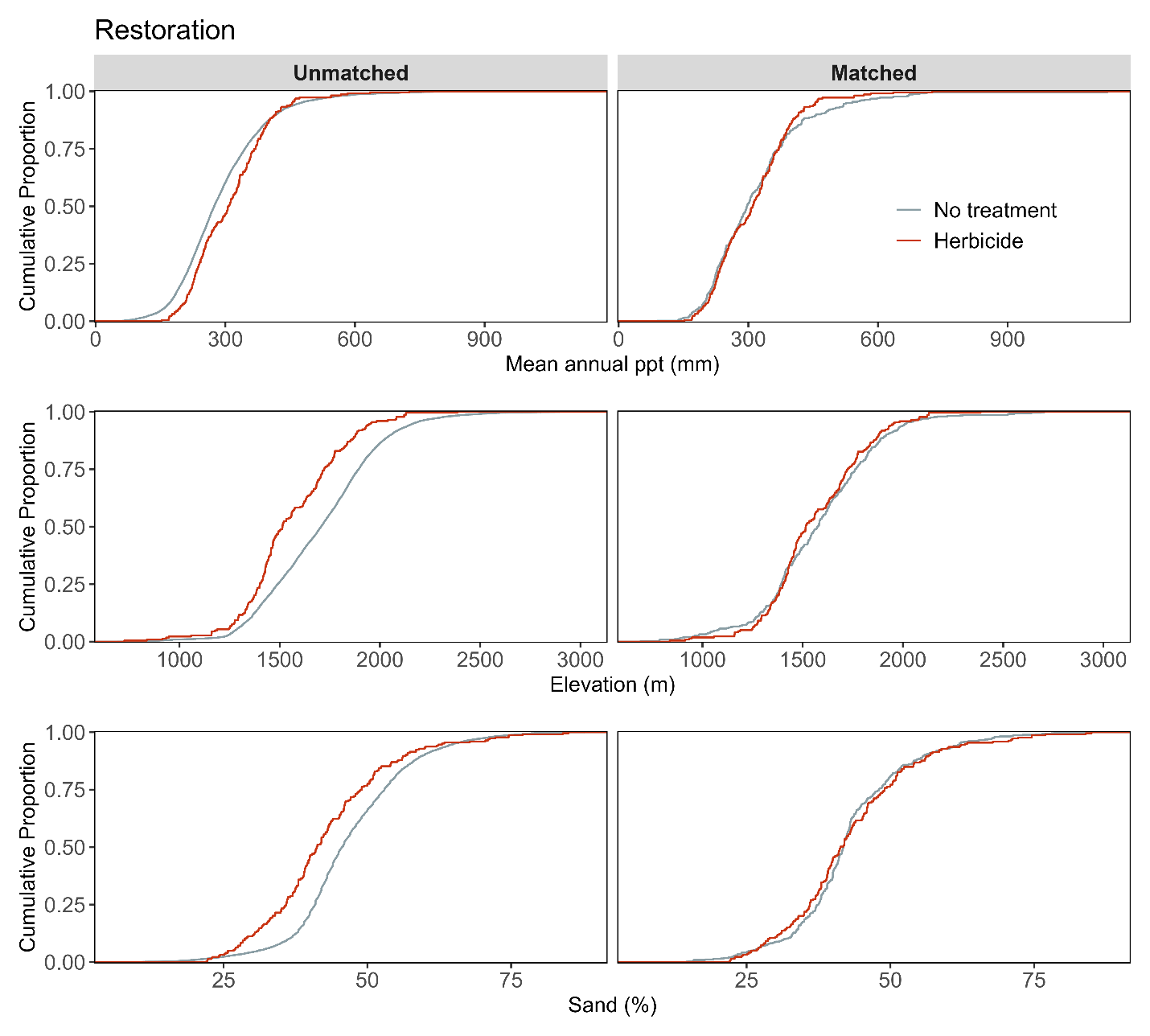

**Figure S4d.** Assessment of covariate balance using PSM for Restoration monitoring plots. The figure compares eCDFs for mean annual precipitation (top), elevation (middle), and sand texture (bottom) percentages between herbicide and non-treated controls, for both unmatched (left column) and matched sets of monitoring plots (right column).

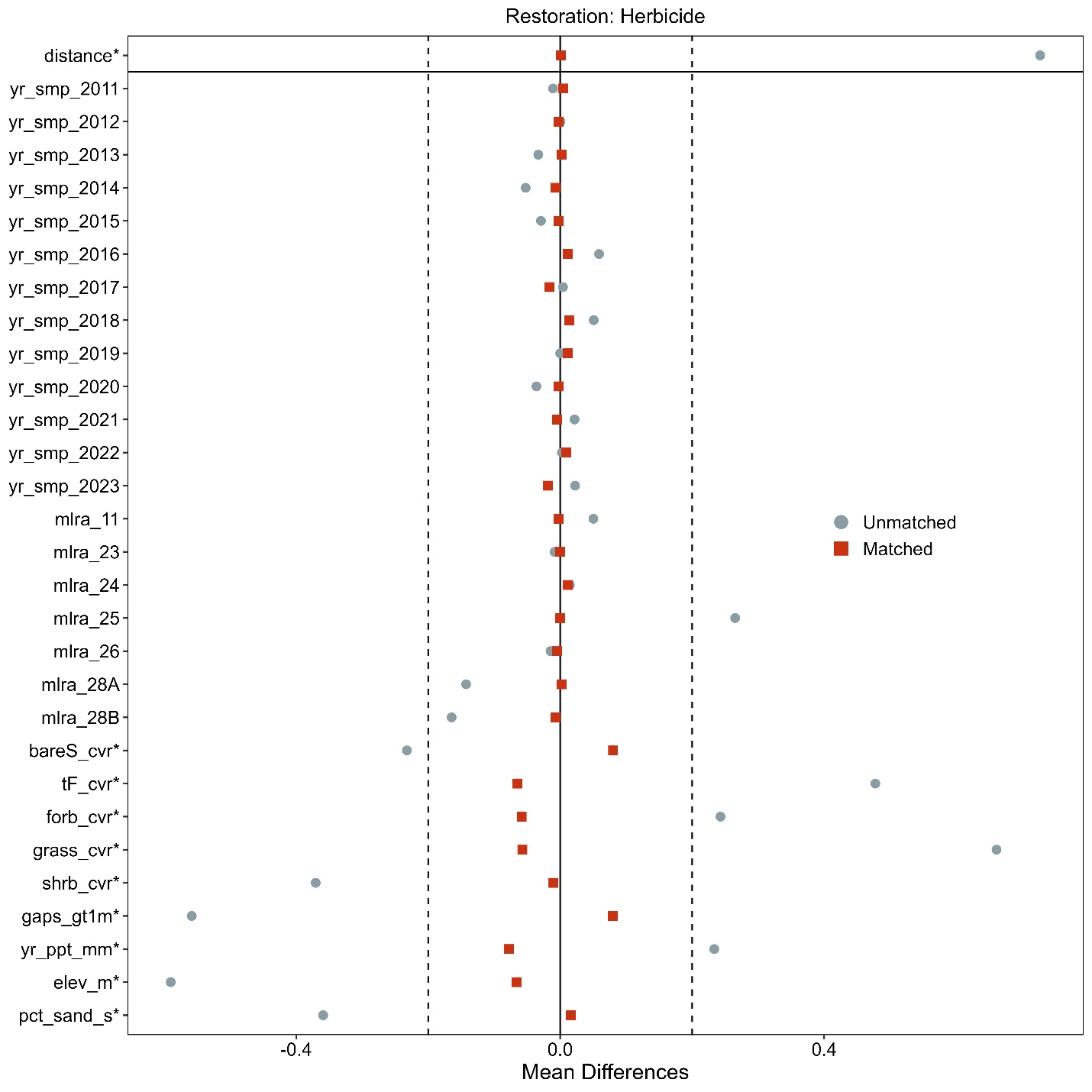

**Figure S4e**. Love-plot illustrating standardized mean differences for covariates in both the full dataset (unmatched) and the matched dataset for herbicide and non-treated controls Restoration plots. Vertical lines indicate the caliper threshold (0.2 times the standard deviation of the propensity score) applied during matching.

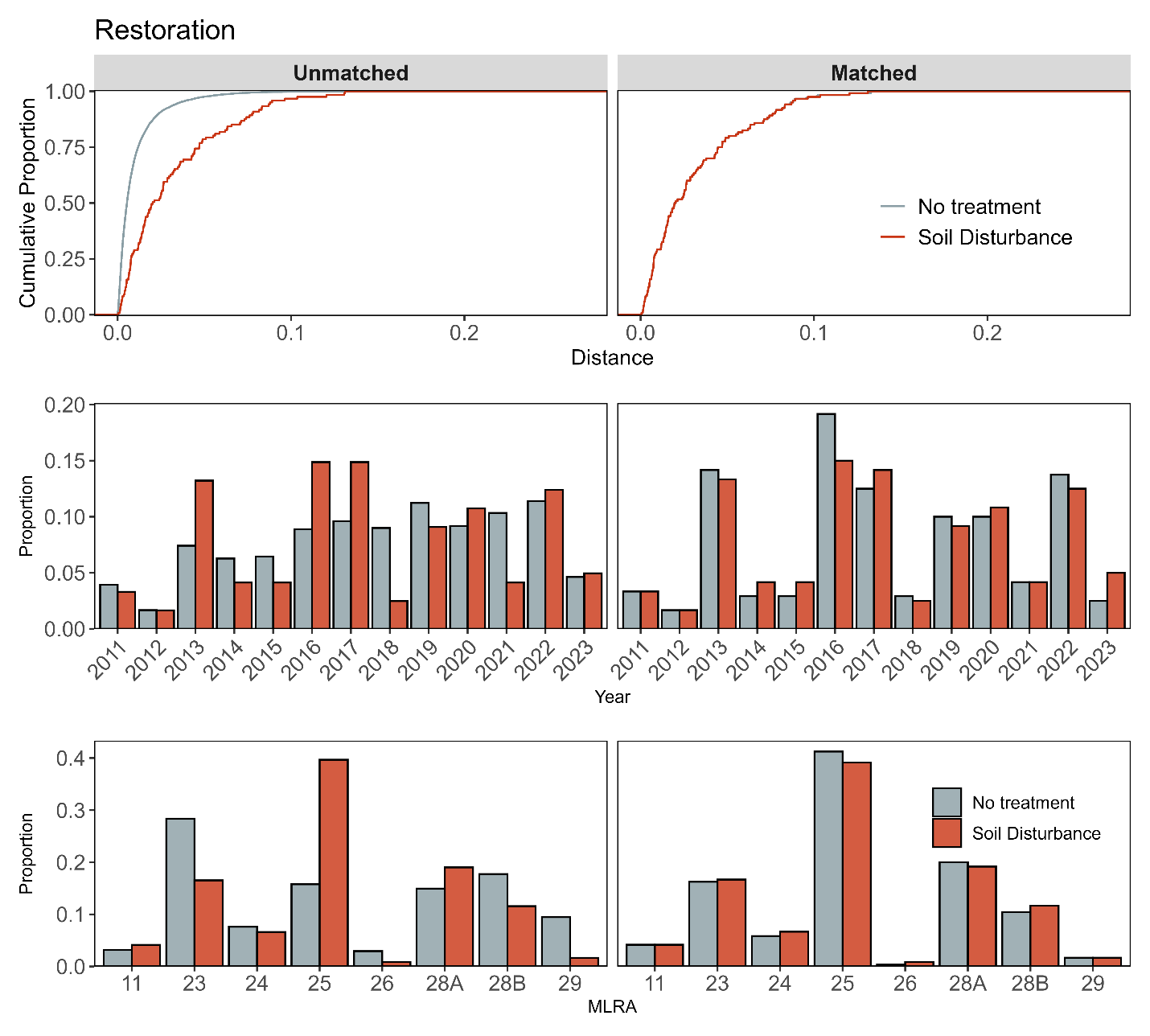

**Figure S5a**. Assessment of covariate balance using propensity score matching (PSM) for Restoration monitoring plots. The top plot shows the empirical cumulative distribution function (eCDF) for the distance measure of the propensity score. The middle and bottom plots display balance in categorical variables, accounting for the year of monitoring and the major land resource area (MLRA) of the plots, comparing unmatched (left column) and matched sets (right columns) for soil disturbance and non-treated control monitoring plots.

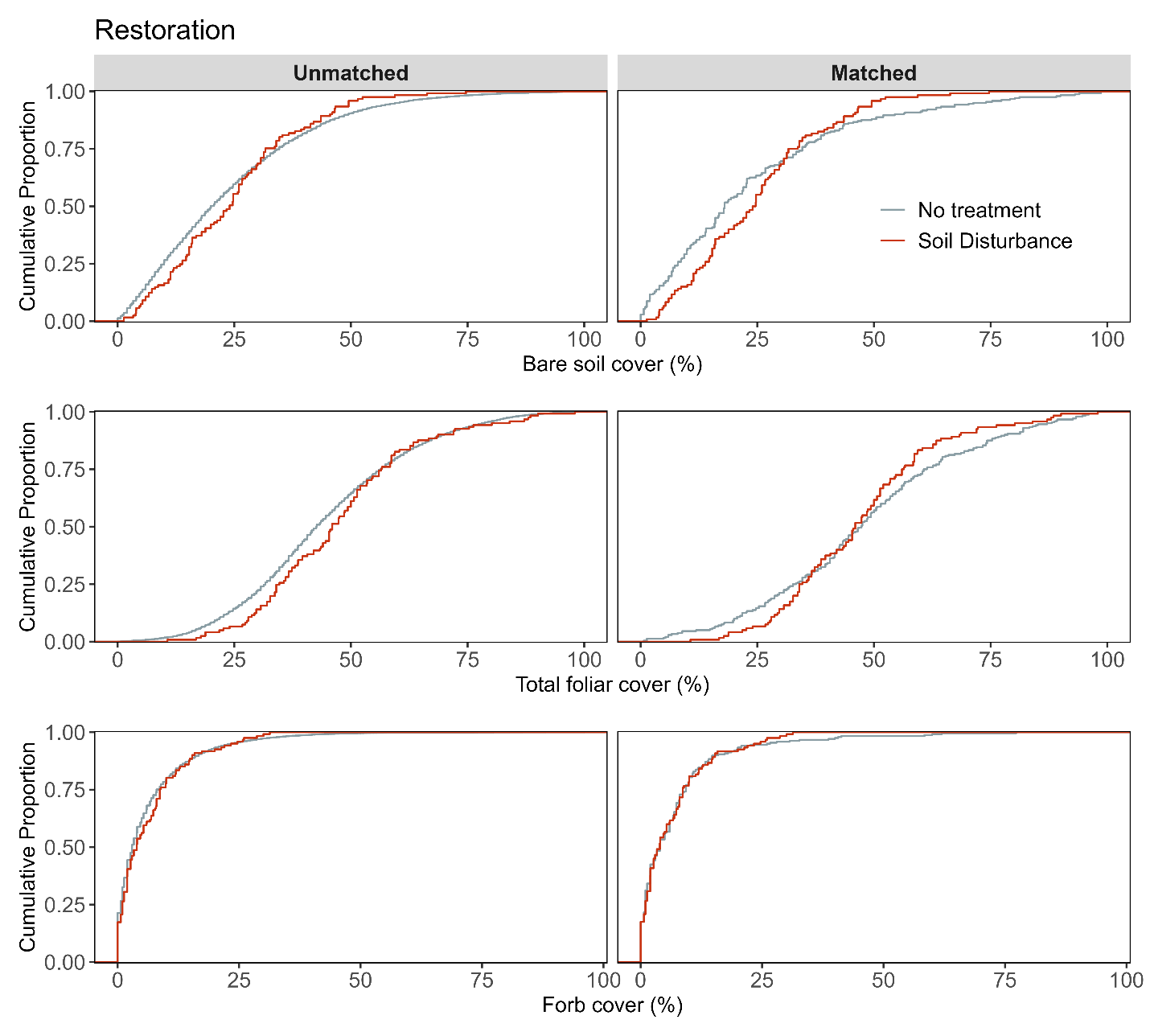

**Figure S5b.** Assessment of covariate balance using PSM for Restoration monitoring plots. The figure compares eCDFs for bare soil (top), total foliar cover (middle), and forb cover (bottom) percentages between soil disturbance and non-treated controls, for both unmatched (left column) and matched sets of monitoring plots (right column).

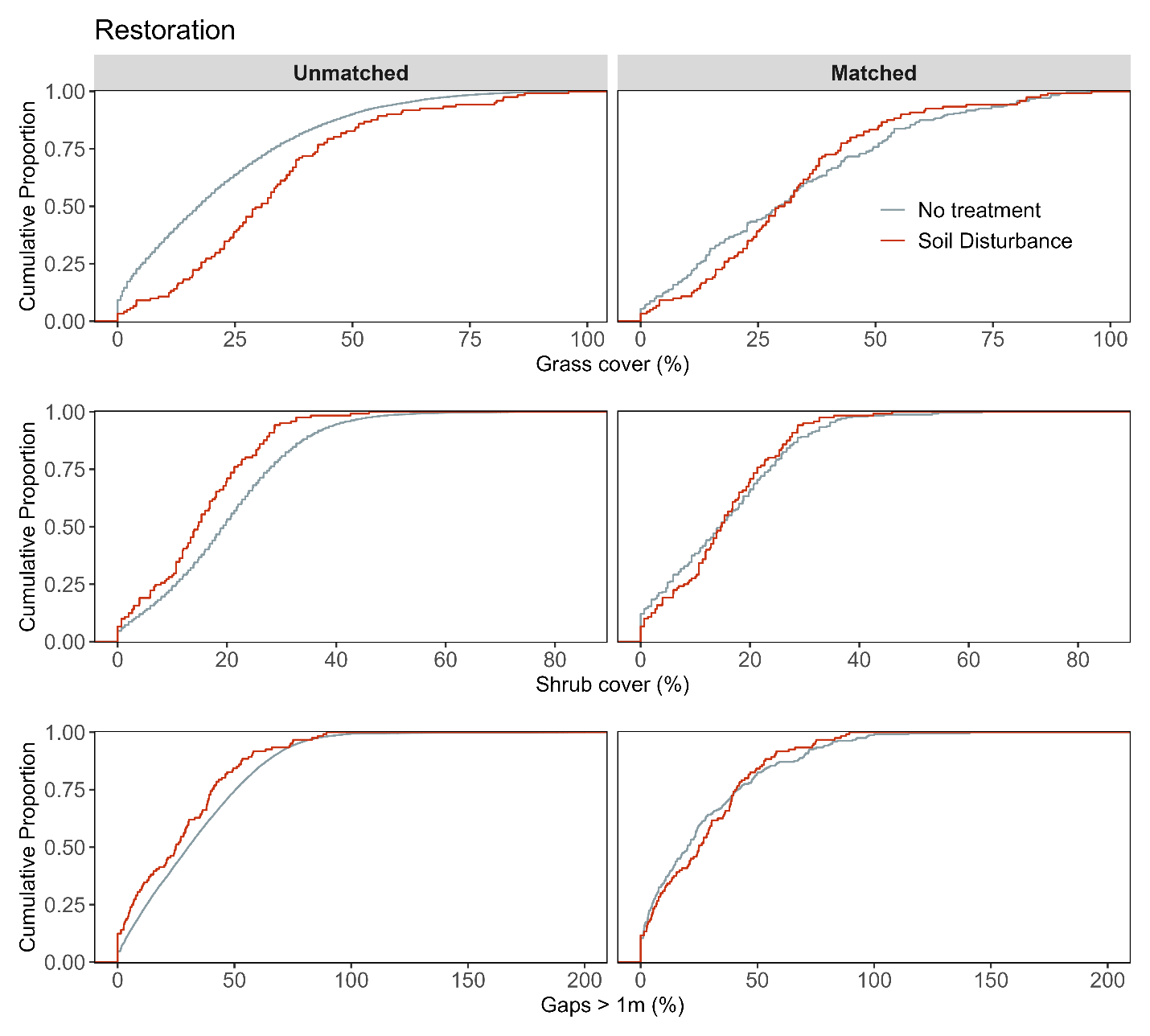

**Figure S5c**. Assessment of covariate balance using PSM for Restoration monitoring plots. The figure compares eCDFs for grass cover (top), shrub cover (middle), and gaps > 1m (bottom) percentages between soil disturbance and non-treated controls, for both unmatched (left column) and matched sets of monitoring plots (right column).

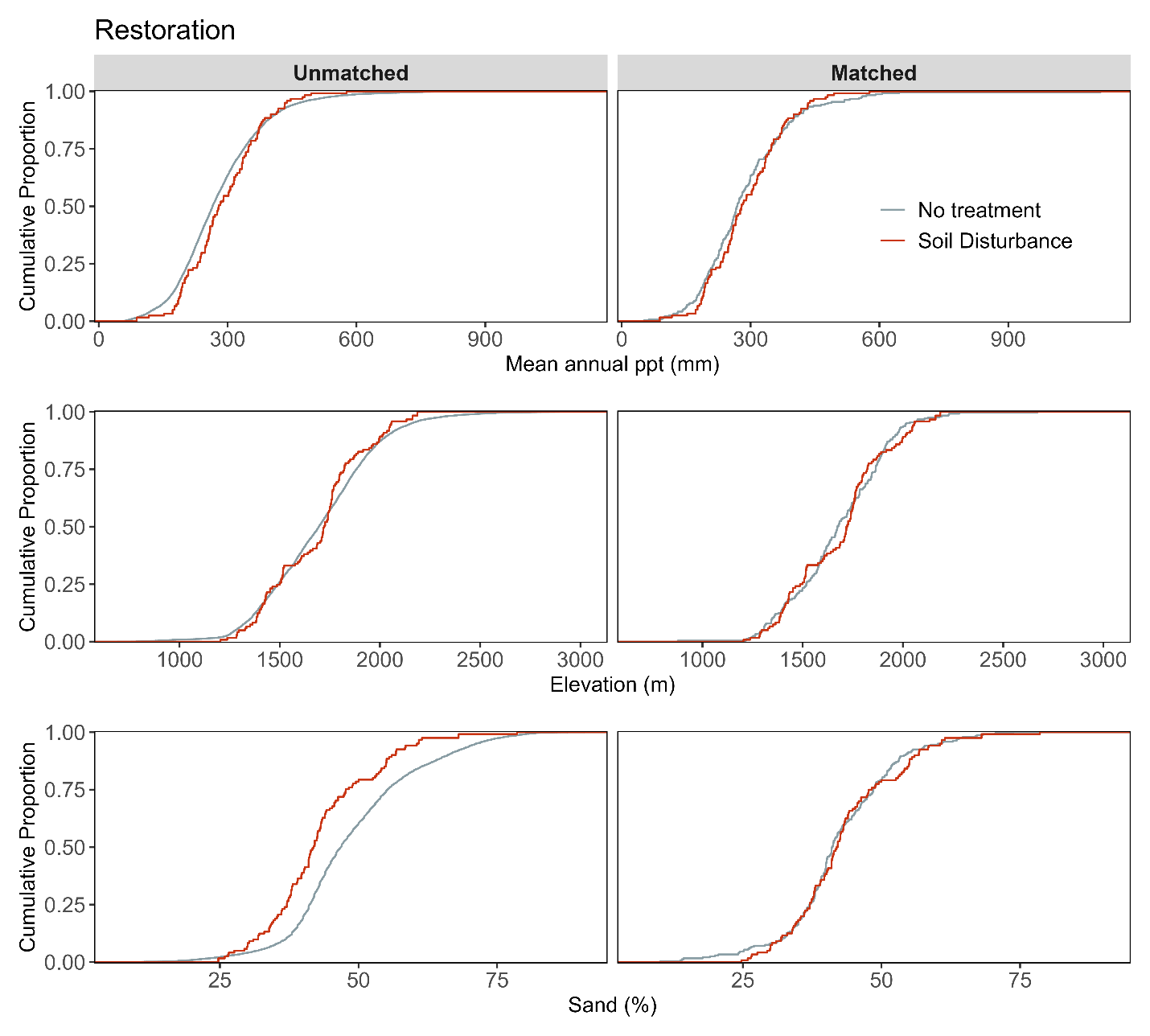
 **Figure S5d.** Assessment of covariate balance using PSM for Restoration monitoring plots. The figure compares eCDFs for mean annual precipitation (top), elevation (middle), and sand texture (bottom) percentages between soil disturbance and non-treated controls, for both unmatched (left column) and matched sets of monitoring plots (right column).

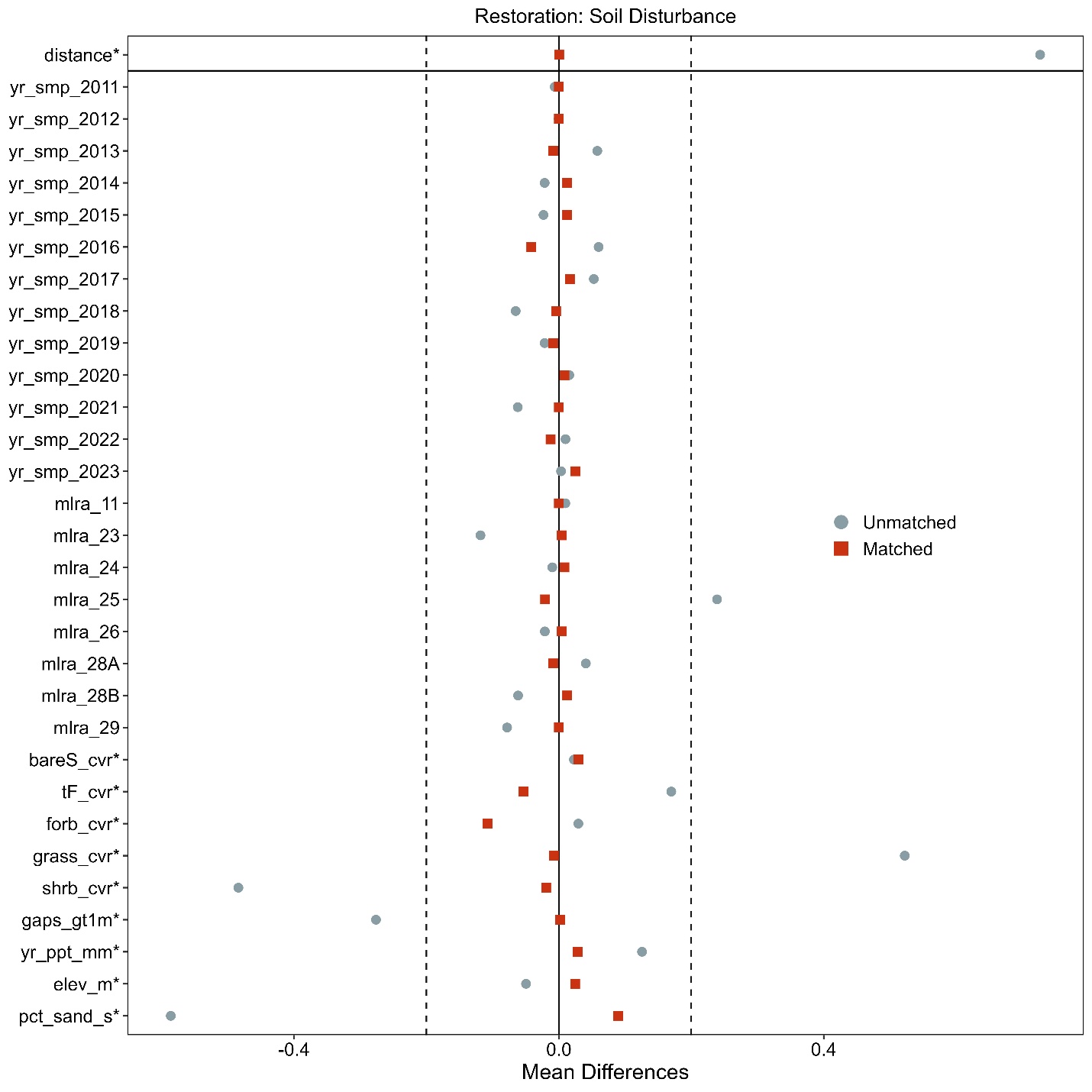

**Figure S5e.** Love-plot illustrating standardized mean differences for covariates in both the full dataset (unmatched) and the matched dataset for soil disturbance and non-treated controls Restoration plots. Vertical lines indicate the caliper threshold (0.2 times the standard deviation of the propensity score) applied during matching.

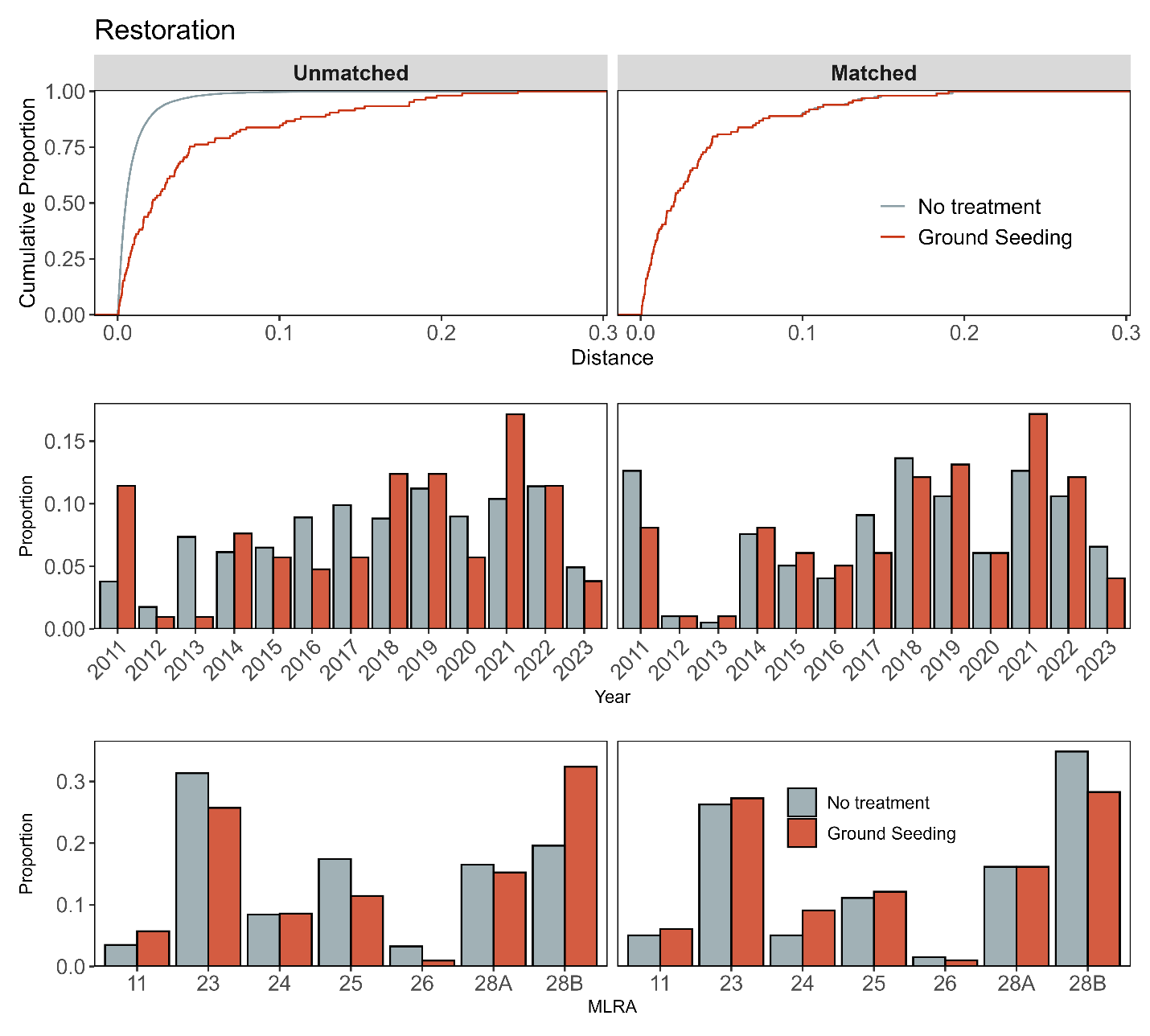
 **Figure S6a.** Assessment of covariate balance using propensity score matching (PSM) for Restoration monitoring plots. The top plot shows the empirical cumulative distribution function (eCDF) for the distance measure of the propensity score. The middle and bottom plots display balance in categorical variables, accounting for the year of monitoring and the major land resource area (MLRA) of the plots, comparing unmatched (left column) and matched sets (right columns) for ground seeding and non-treated control monitoring plots.

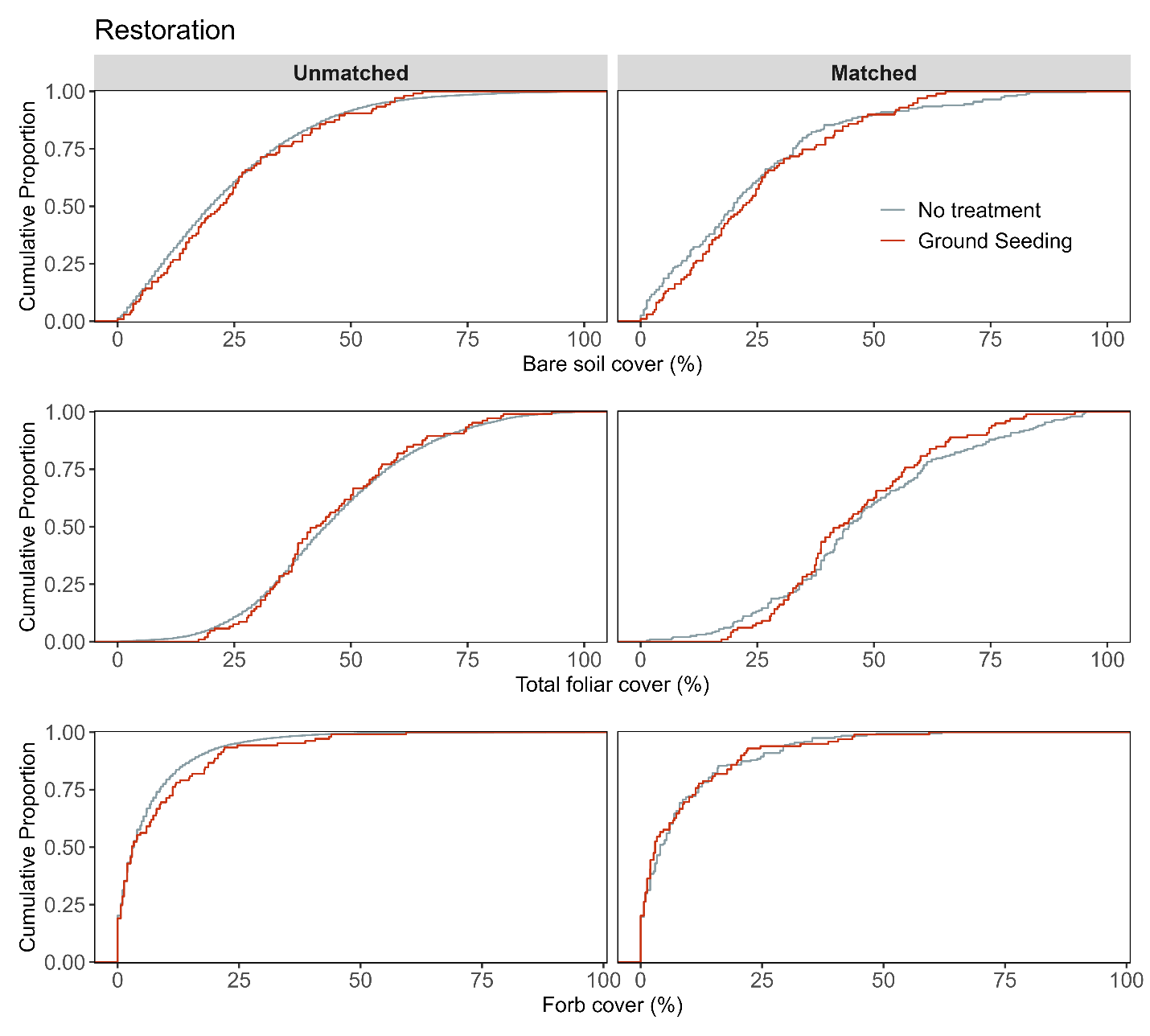

**Figure S6b**. Assessment of covariate balance using PSM for Restoration monitoring plots. The figure compares eCDFs for bare soil (top), total foliar cover (middle), and forb cover (bottom) percentages between ground seeding and non-treated controls, for both unmatched (left column) and matched sets of monitoring plots (right column).

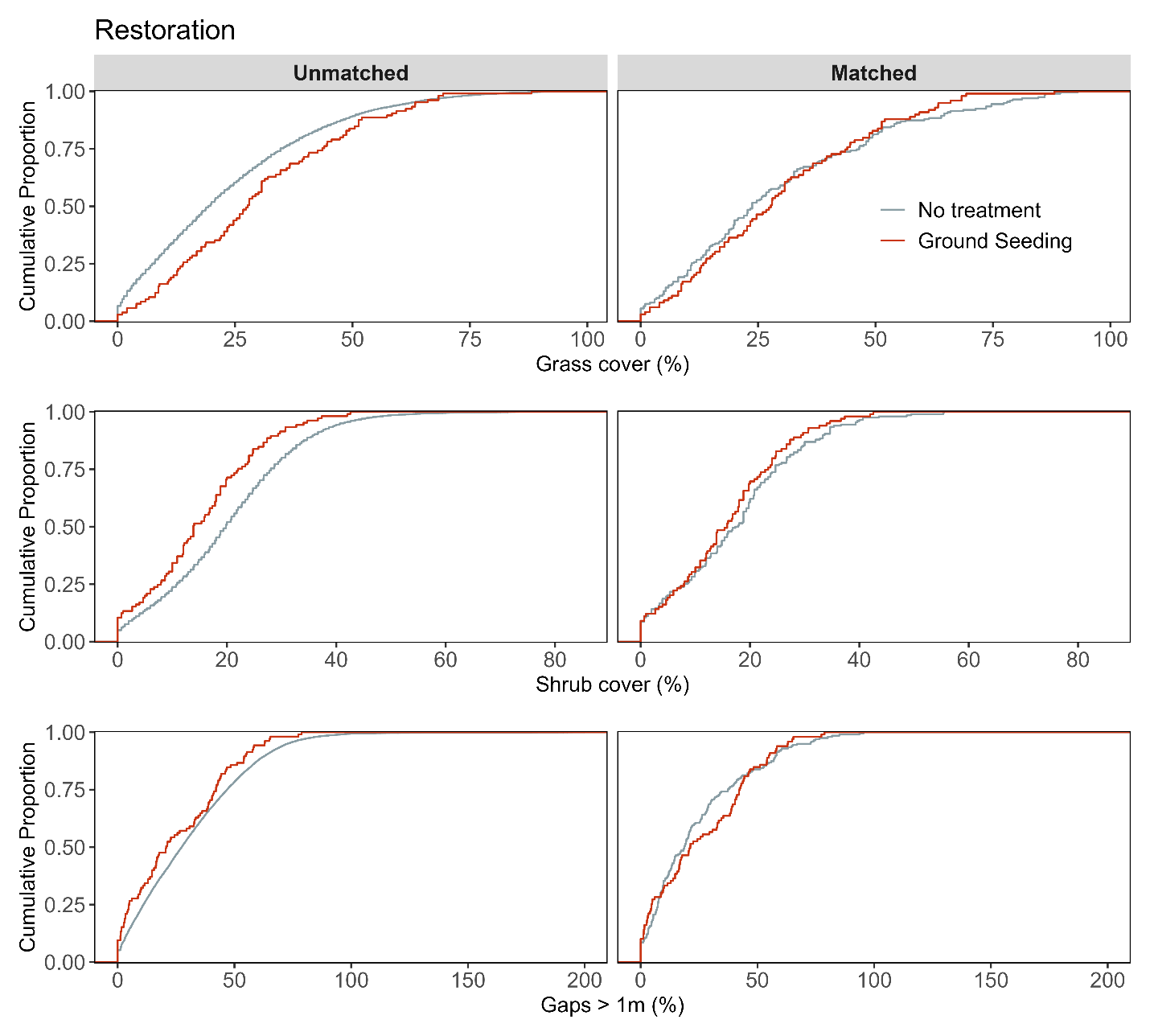
 **Figure S6c**. Assessment of covariate balance using PSM for Restoration monitoring plots. The figure compares eCDFs for grass cover (top), shrub cover (middle), and gaps > 1m (bottom) percentages between ground seeding and non-treated controls, for both unmatched (left column) and matched sets of monitoring plots (right column).

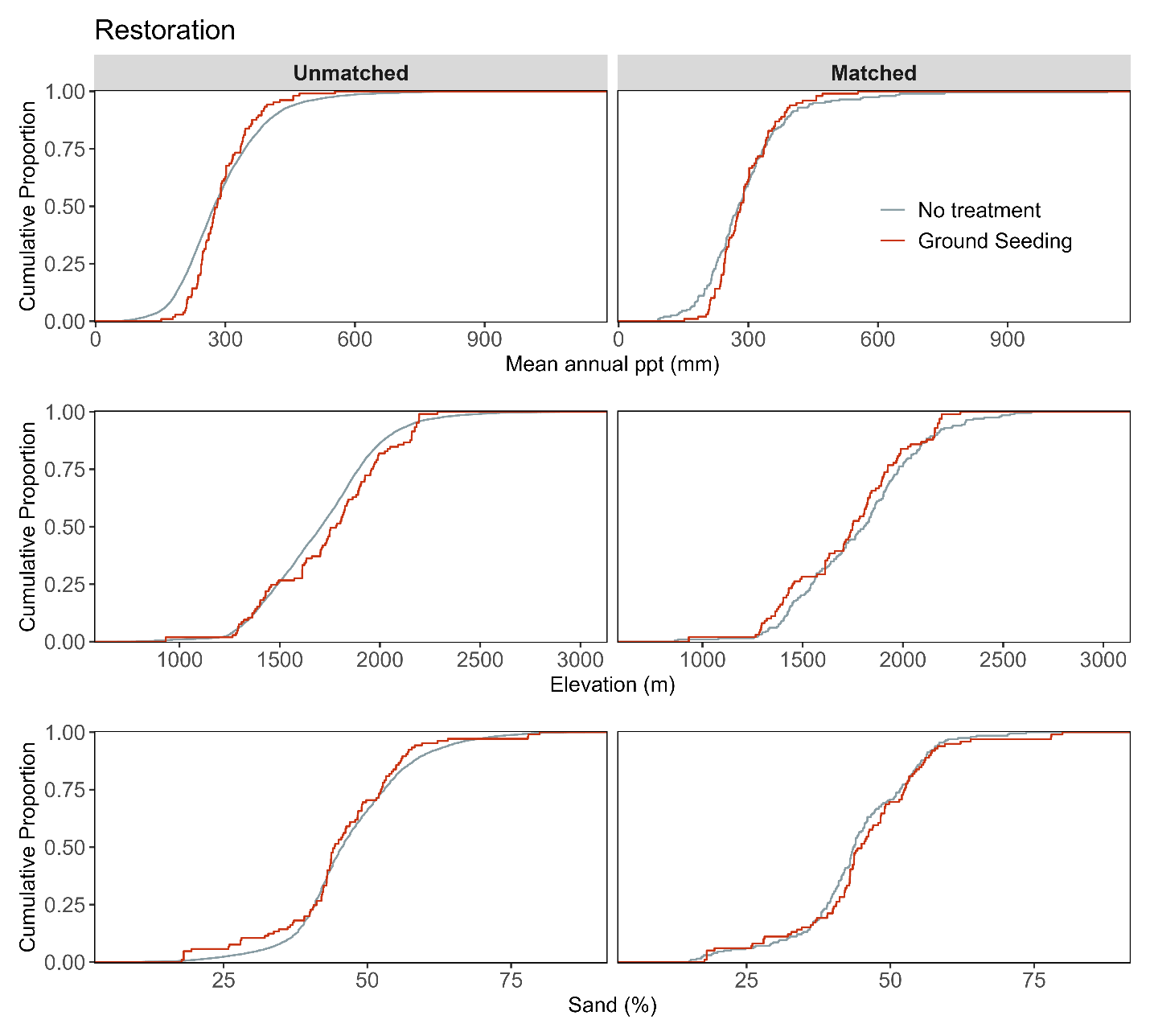

**Figure S6d.** Assessment of covariate balance using PSM for Restoration monitoring plots. The figure compares eCDFs for mean annual precipitation (top), elevation (middle), and sand texture (bottom) percentages between ground seeding and non-treated controls, for both unmatched (left column) and matched sets of monitoring plots (right column).

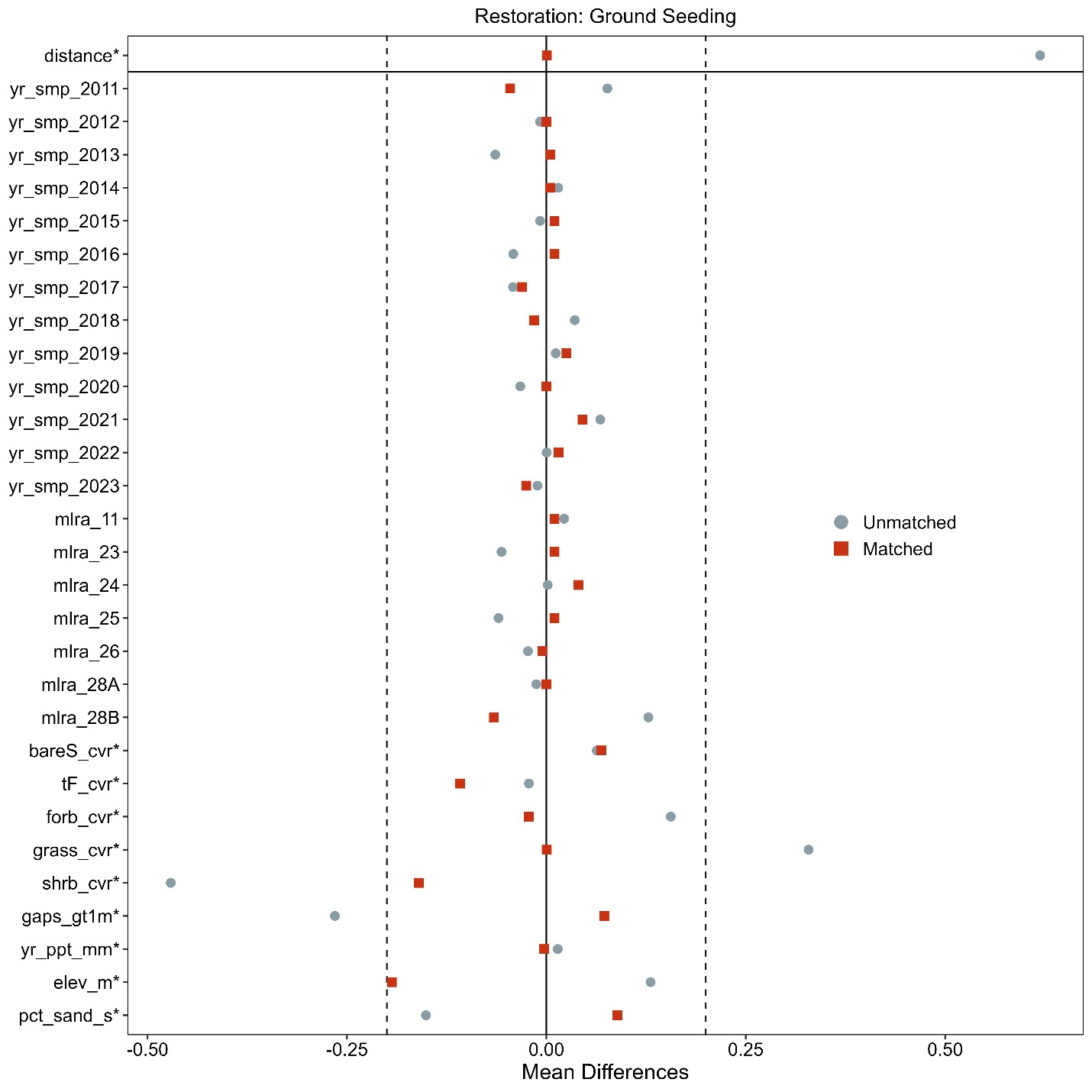

**Figure S4.6e.** Love-plot illustrating standardized mean differences for covariates in both the full dataset (unmatched) and the matched dataset for ground seeding and non-treated controls Restoration plots. Vertical lines indicate the caliper threshold (0.2 times the standard deviation of the propensity score) applied during matching.

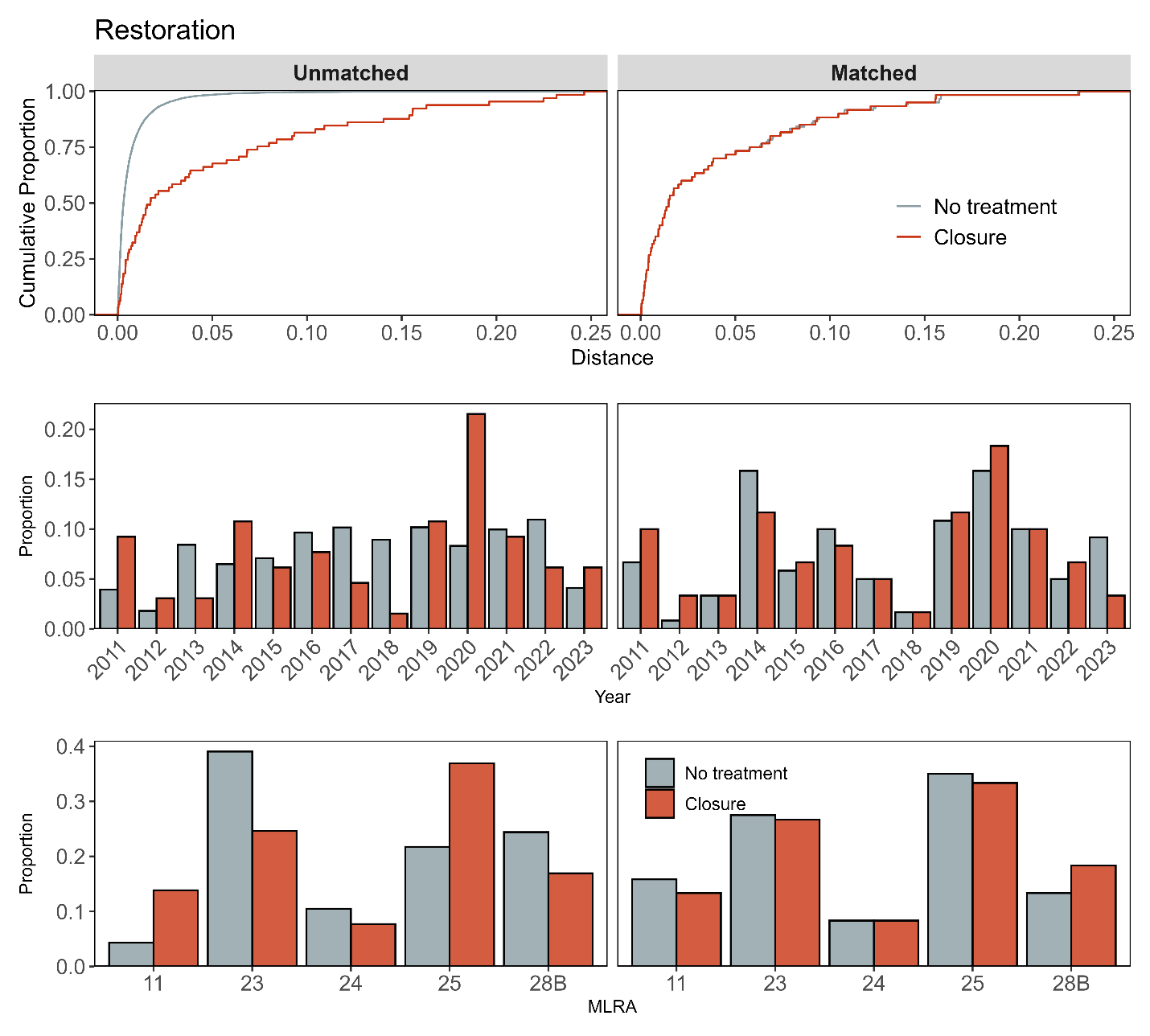

**Figure S7a**. Assessment of covariate balance using propensity score matching (PSM) for Restoration monitoring plots. The top plot shows the empirical cumulative distribution function (eCDF) for the distance measure of the propensity score. The middle and bottom plots display balance in categorical variables, accounting for the year of monitoring and the major land resource area (MLRA) of the plots, comparing unmatched (left column) and matched sets (right columns) for closure and non-treated control monitoring plots.

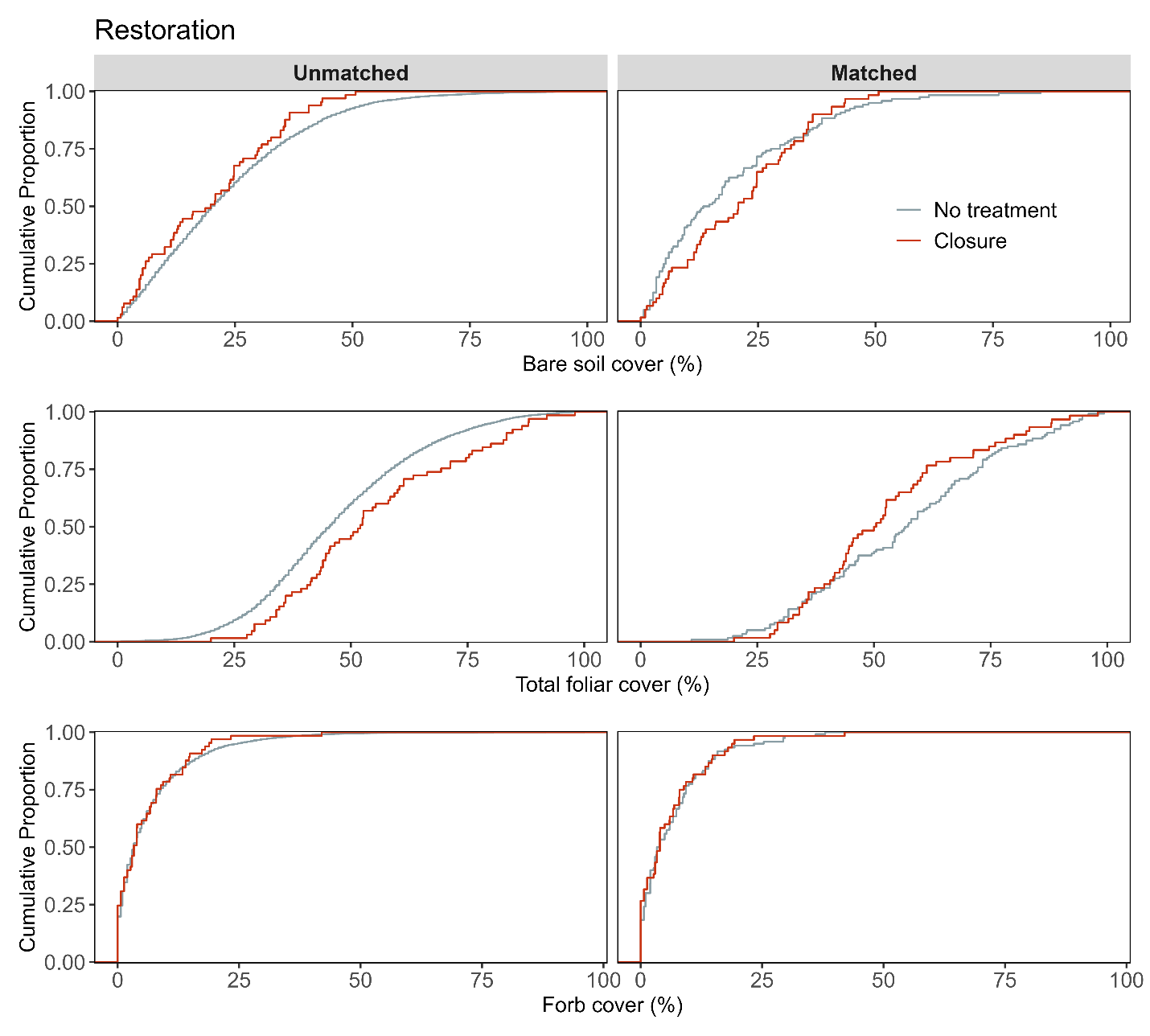

**Figure S7b.** Assessment of covariate balance using PSM for Restoration monitoring plots. The figure compares eCDFs for bare soil (top), total foliar cover (middle), and forb cover (bottom) percentages between closure and non-treated controls, for both unmatched (left column) and matched sets of monitoring plots (right column).

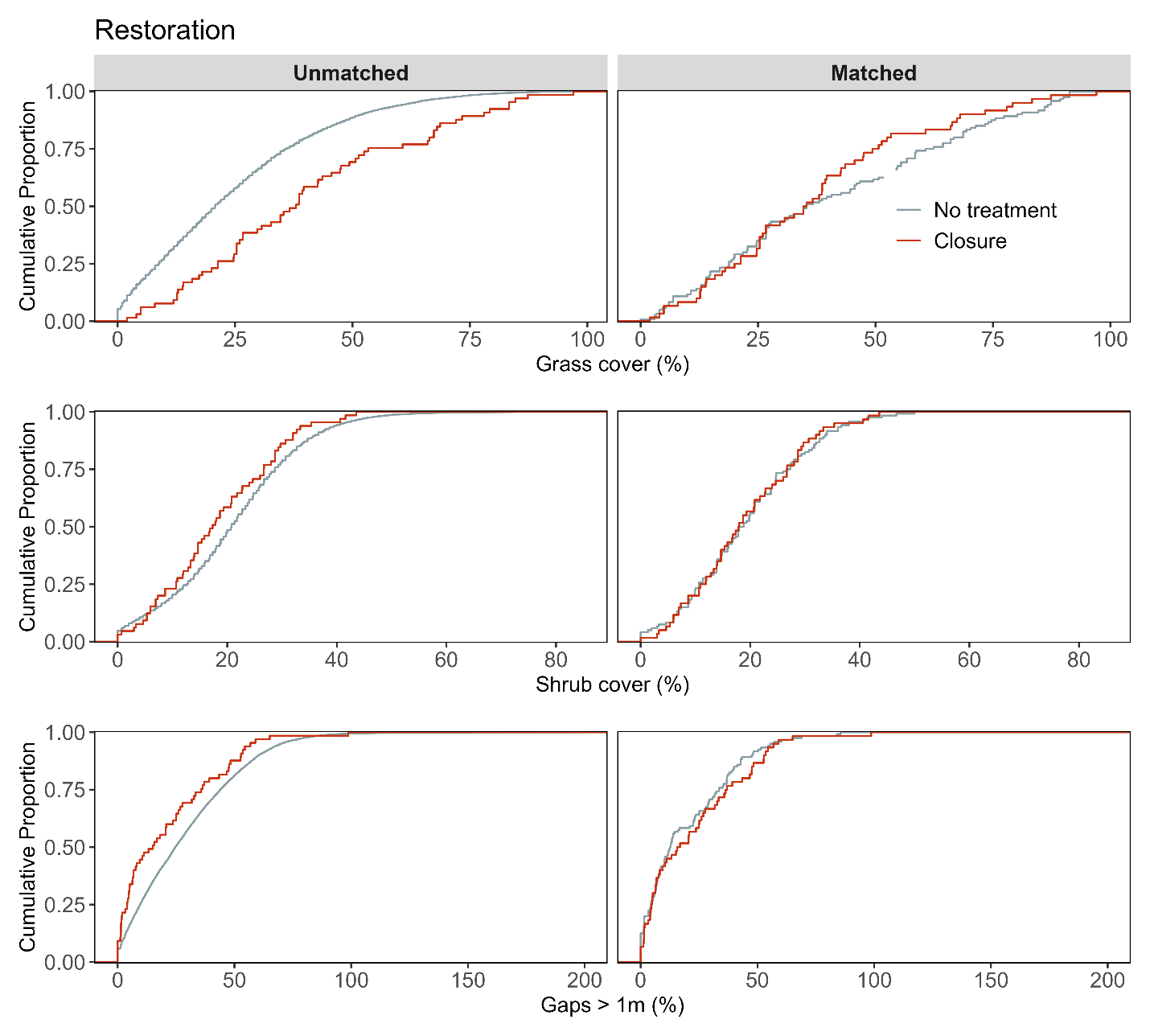

**Figure S7c.** Assessment of covariate balance using PSM for Restoration monitoring plots. The figure compares eCDFs for grass cover (top), shrub cover (middle), and gaps > 1m (bottom) percentages between closure and non-treated controls, for both unmatched (left column) and matched sets of monitoring plots (right column).

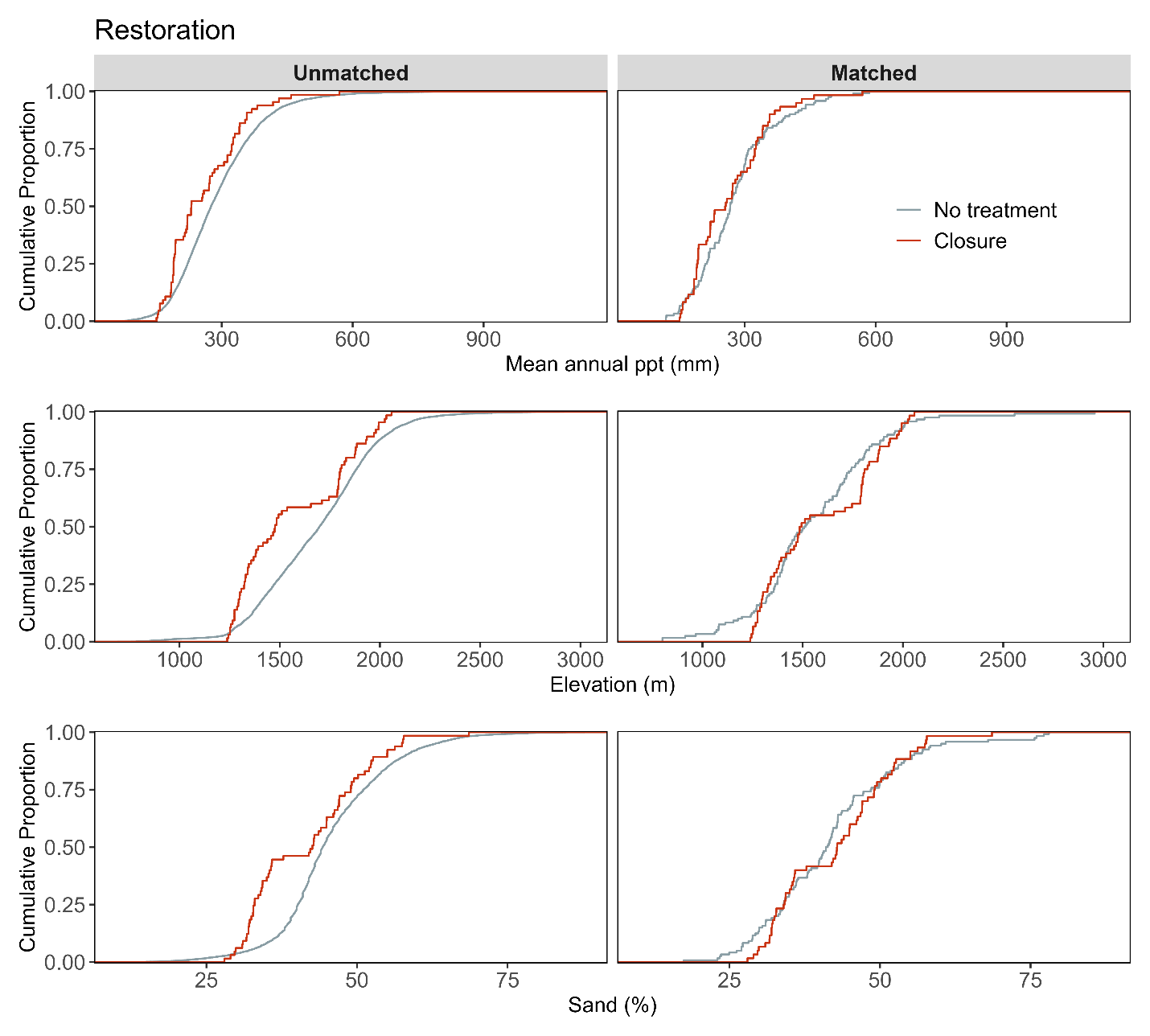

**Figure S7d.** Assessment of covariate balance using PSM for Restoration monitoring plots. The figure compares eCDFs for mean annual precipitation (top), elevation (middle), and sand texture (bottom) percentages between closure and non-treated controls, for both unmatched (left column) and matched sets of monitoring plots (right column).

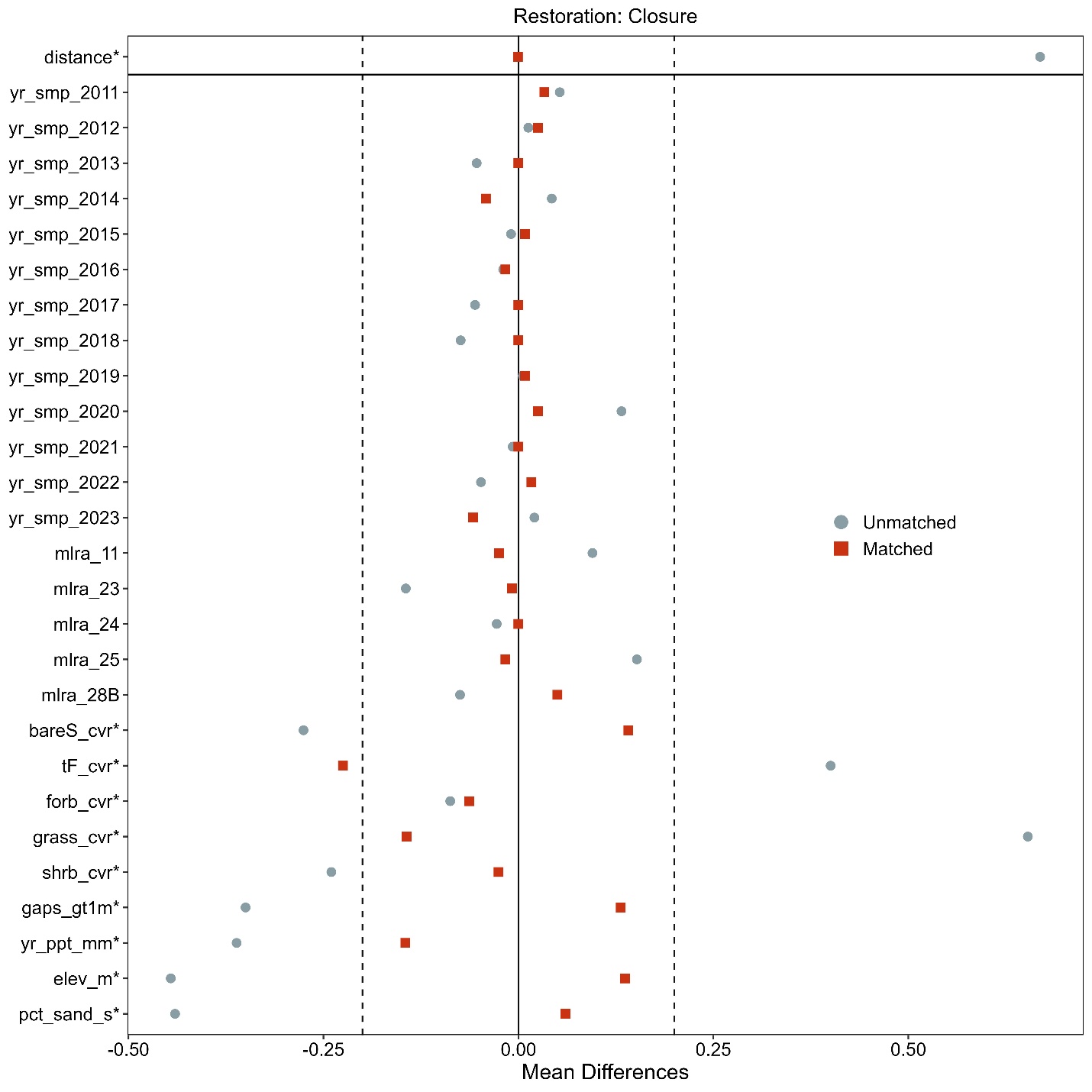

**Figure S7e**. Love-plot illustrating standardized mean differences for covariates in both the full dataset (unmatched) and the matched dataset for closure and non-treated controls Restoration plots. Vertical lines indicate the caliper threshold (0.2 times the standard deviation of the propensity score) applied during matching.

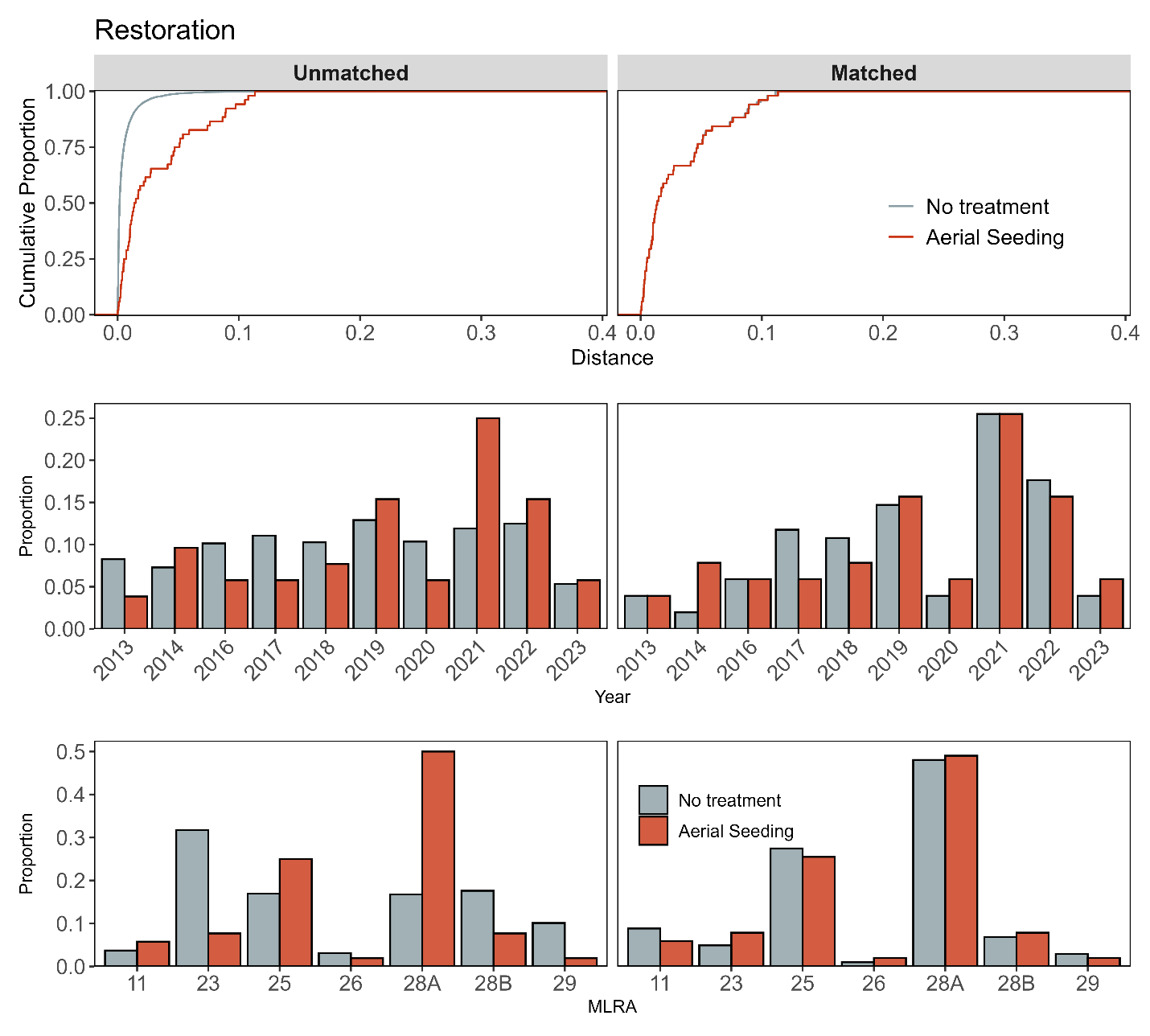

**Figure S8a**. Assessment of covariate balance using propensity score matching (PSM) for Restoration monitoring plots. The top plot shows the empirical cumulative distribution function (eCDF) for the distance measure of the propensity score. The middle and bottom plots display balance in categorical variables, accounting for the year of monitoring and the major land resource area (MLRA) of the plots, comparing unmatched (left column) and matched sets (right columns) for aerial seeding and non-treated control monitoring plots.

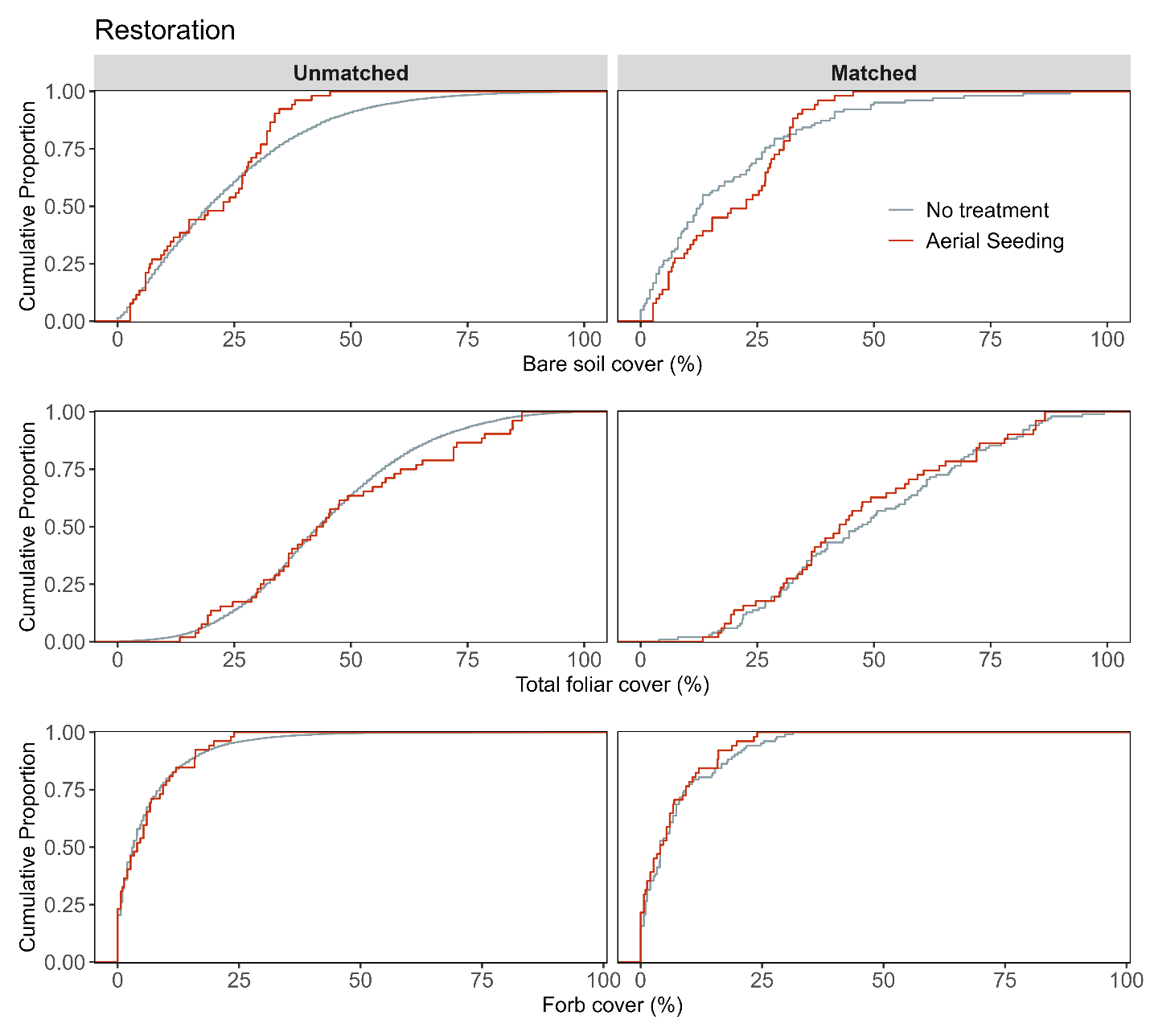

**Figure S8b**. Assessment of covariate balance using PSM for Restoration monitoring plots. The figure compares eCDFs for bare soil (top), total foliar cover (middle), and forb cover (bottom) percentages between aerial seeding and non-treated controls, for both unmatched (left column) and matched sets of monitoring plots (right column).

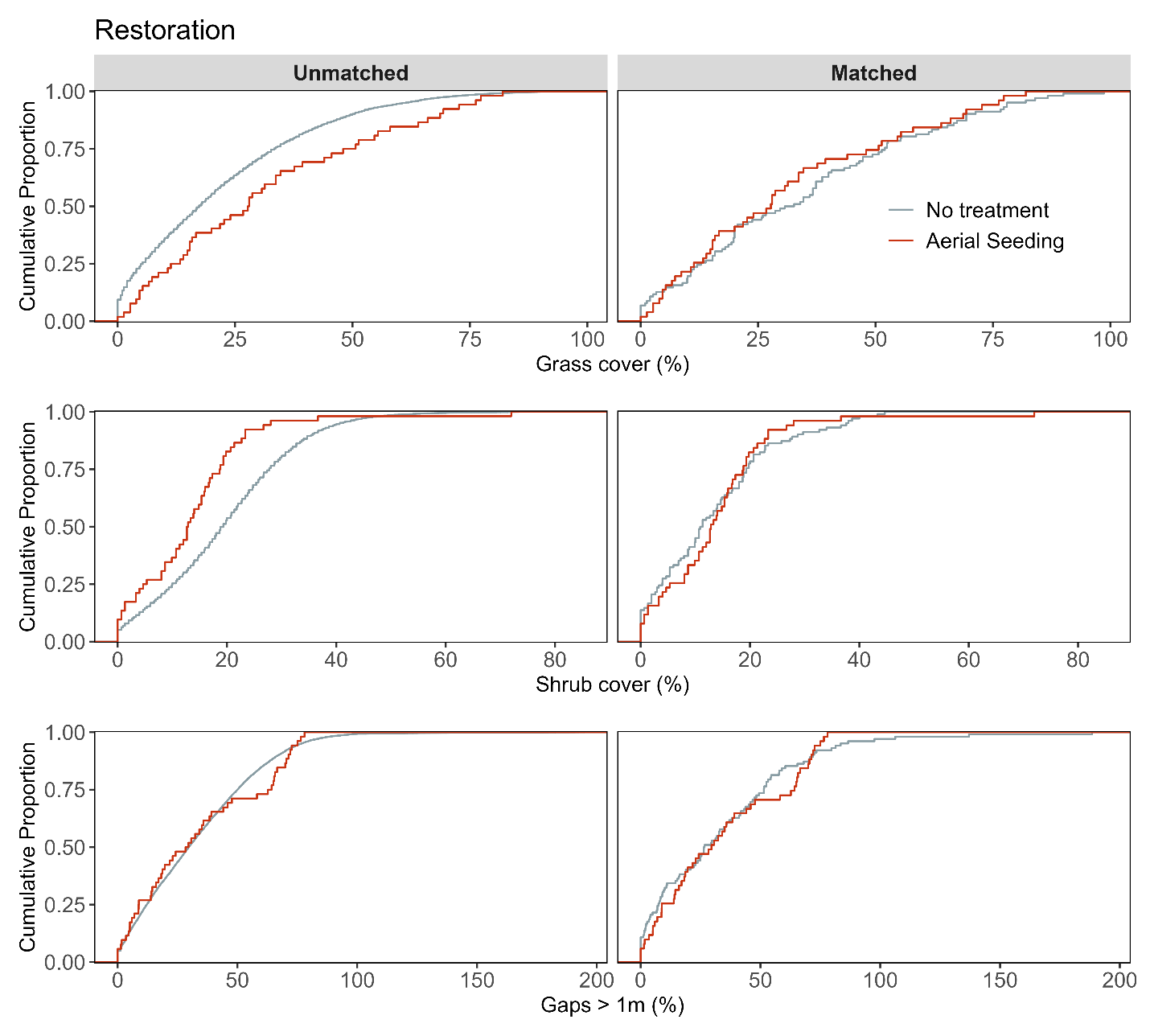

**Figure S8c**. Assessment of covariate balance using PSM for Restoration monitoring plots. The figure compares eCDFs for grass cover (top), shrub cover (middle), and gaps > 1m (bottom) percentages between aerial seeding and non-treated controls, for both unmatched (left column) and matched sets of monitoring plots (right column).

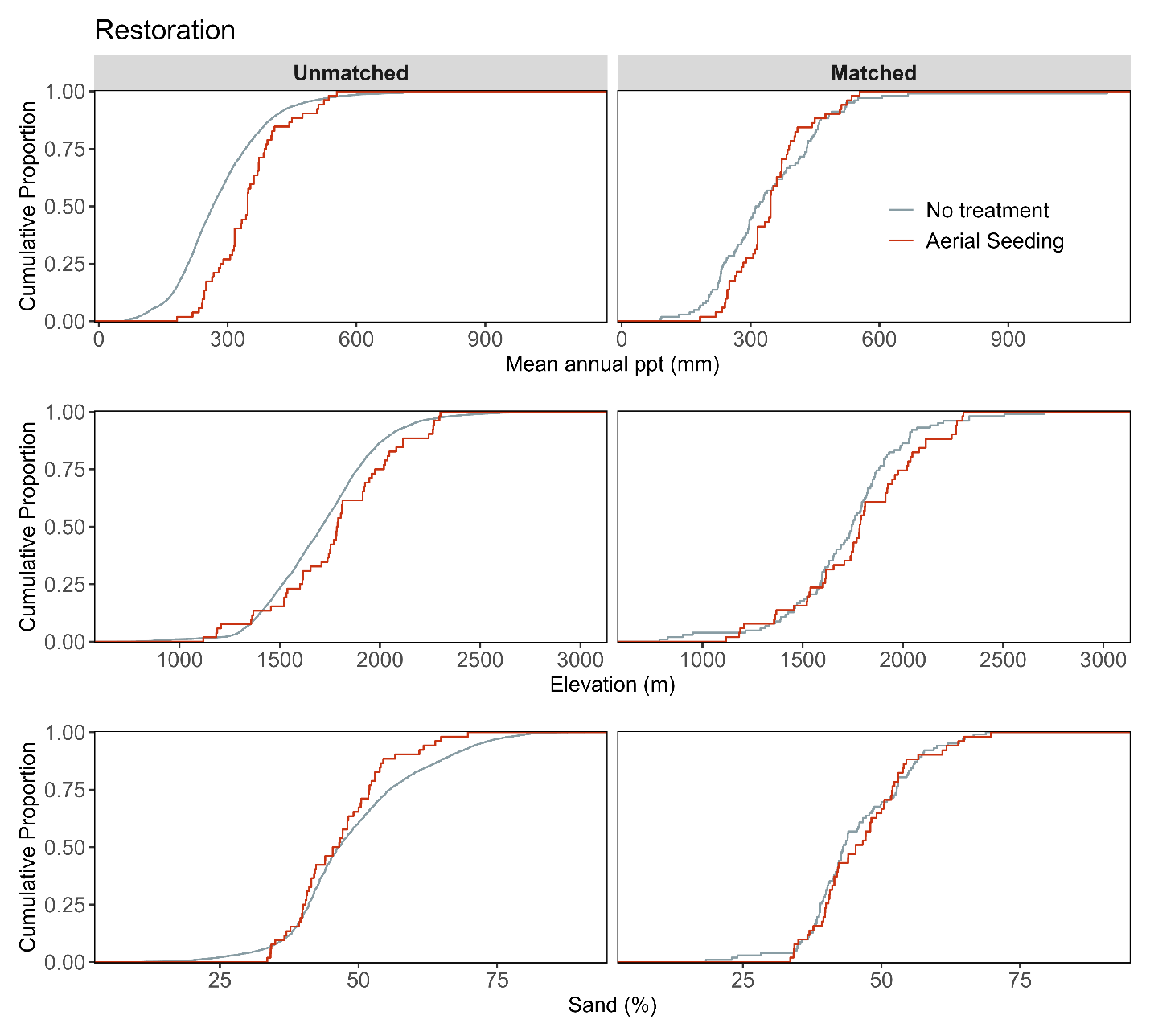

**Figure S8d**. Assessment of covariate balance using PSM for Restoration monitoring plots. The figure compares eCDFs for mean annual precipitation (top), elevation (middle), and sand texture (bottom) percentages between aerial seeding and non-treated controls, for both unmatched (left column) and matched sets of monitoring plots (right column).

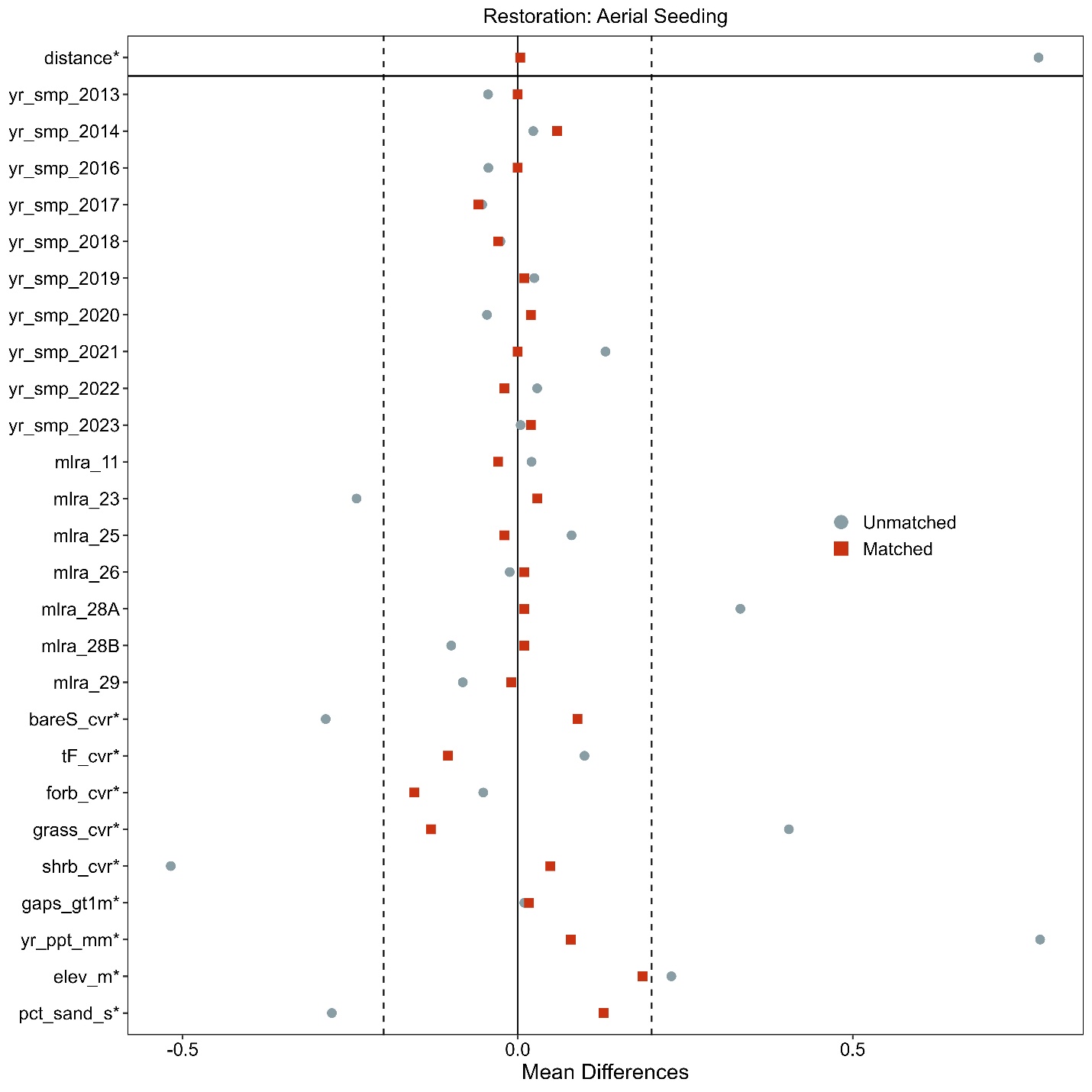

**Figure S8e.** Love-plot illustrating standardized mean differences for covariates in both the full dataset (unmatched) and the matched dataset for aerial seeding and non-treated controls Restoration plots. Vertical lines indicate the caliper threshold (0.2 times the standard deviation of the propensity score) applied during matching.

**Supporting information for Post-fire rehabilitation plots and non-treated controls**

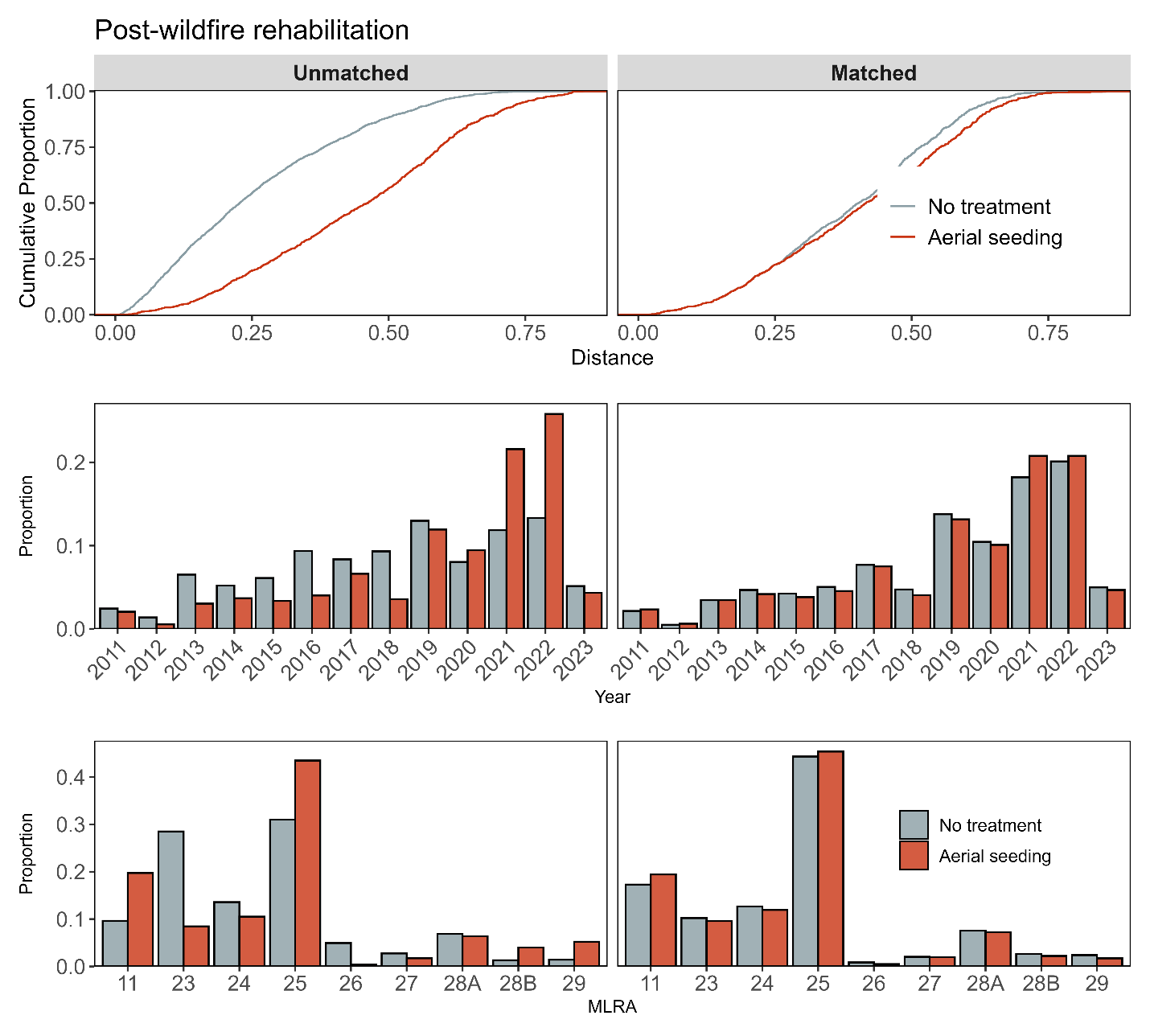

**Figure S9a.** Assessment of covariate balance using propensity score matching (PSM) for Post-fire rehabilitation monitoring plots. The top plot shows the empirical cumulative distribution function (eCDF) for the distance measure of the propensity score. The middle and bottom plots display balance in categorical variables, accounting for the year of monitoring and the major land resource area (MLRA) of the plots, comparing unmatched (left column) and matched sets (right columns) for aerial seeding and non-treated control monitoring plots.

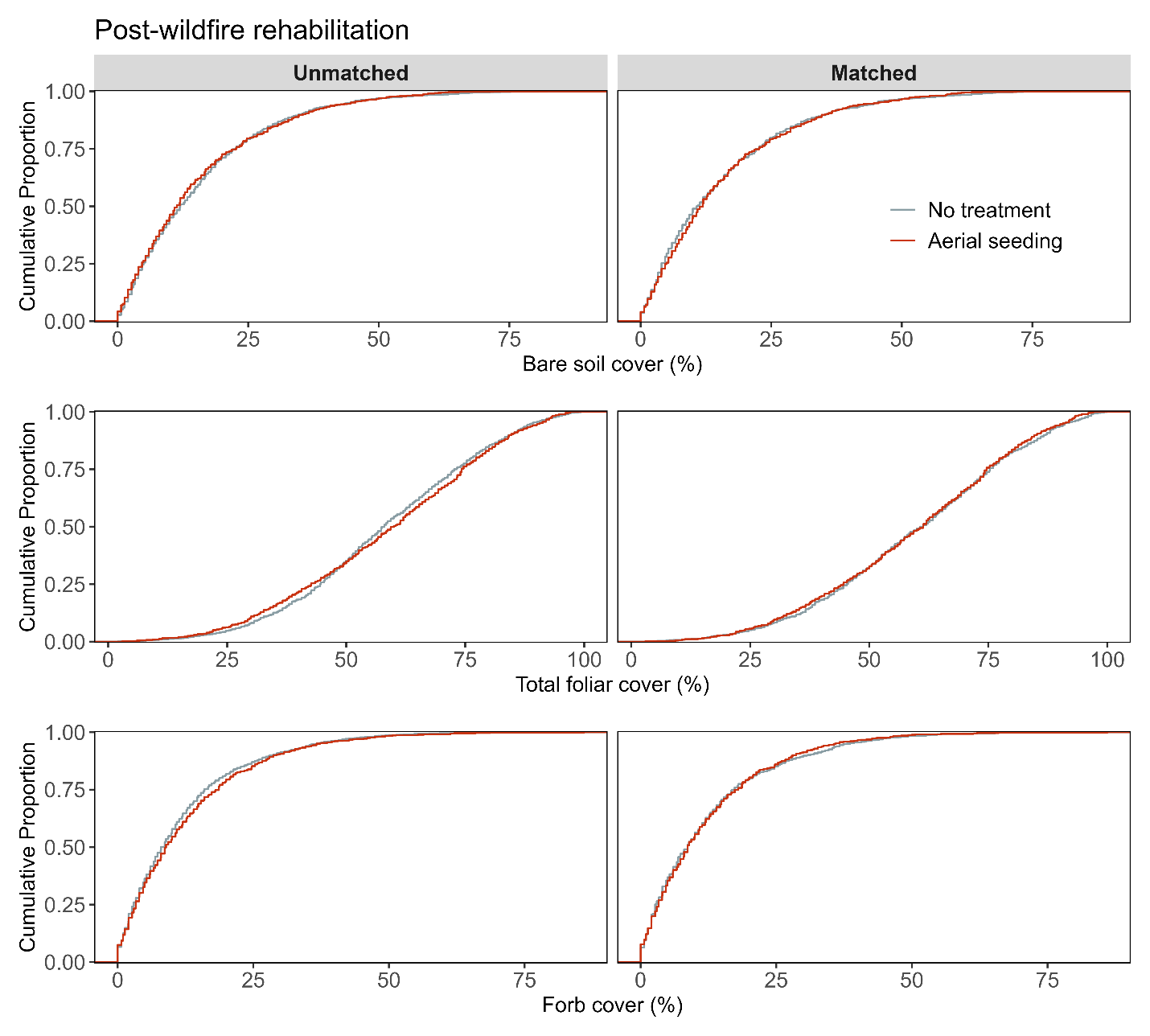

**Figure S9b**. Assessment of covariate balance using PSM for Post-fire rehabilitation monitoring plots. The figure compares eCDFs for bare soil (top), total foliar cover (middle), and forb cover (bottom) percentages between aerial seeding and non-treated controls, for both unmatched (left column) and matched sets of monitoring plots (right column).

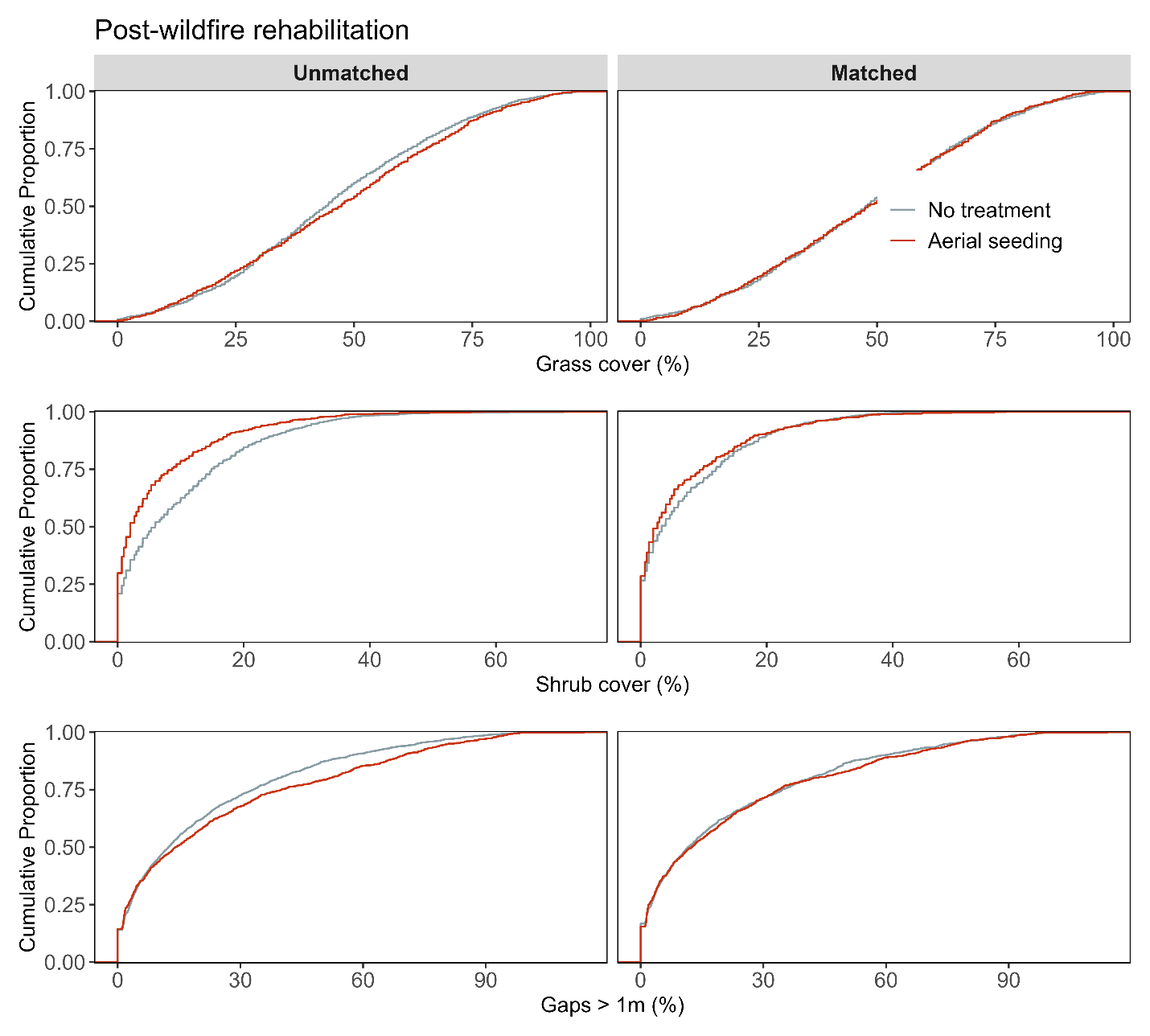

**Figure S9c**. Assessment of covariate balance using PSM for Post-fire rehabilitation monitoring plots. The figure compares eCDFs for grass cover (top), shrub cover (middle), and gaps > 1m (bottom) percentages between aerial seeding and non-treated controls, for both unmatched (left column) and matched sets of monitoring plots (right column).

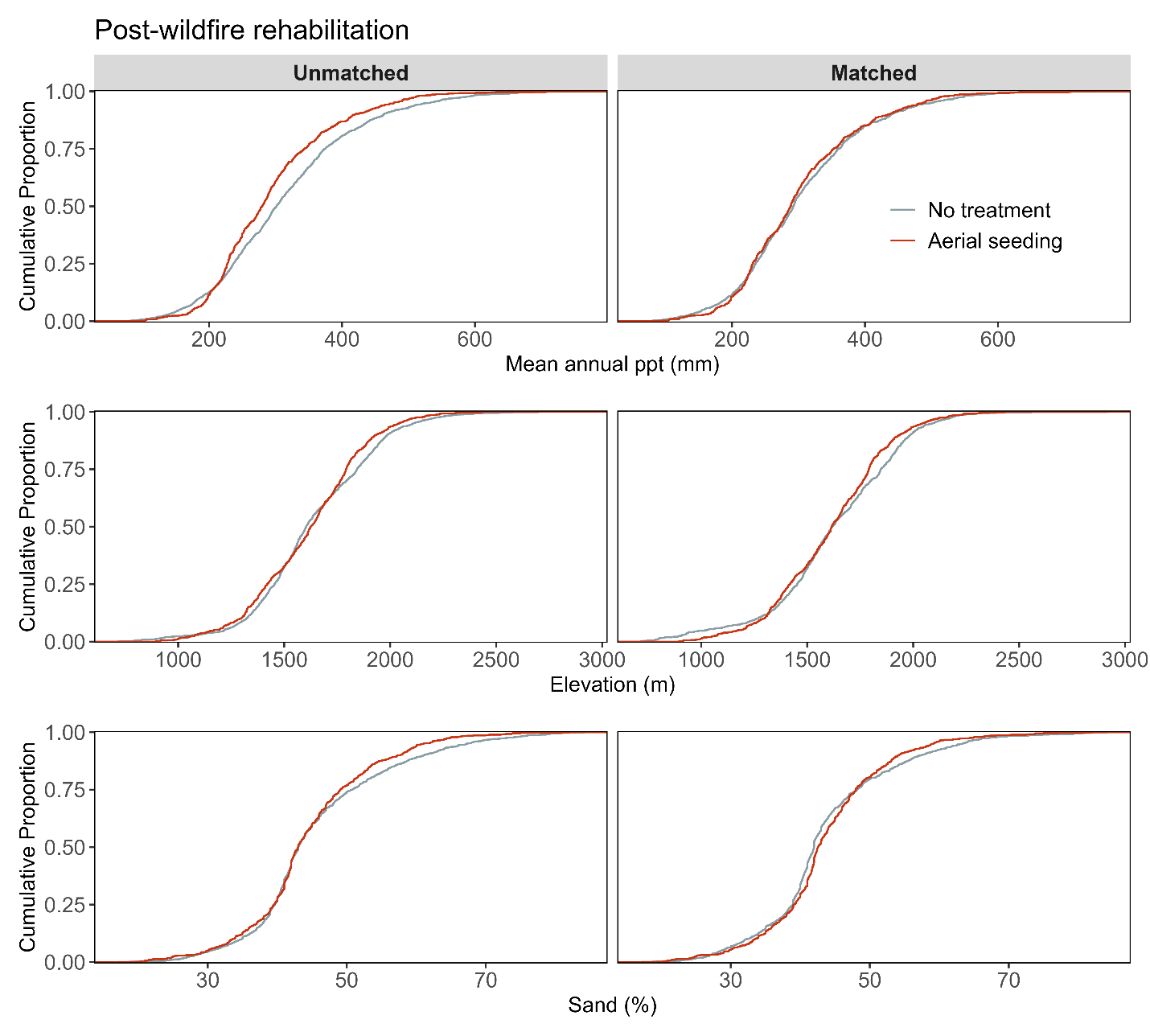

**Figure S9d**. Assessment of covariate balance using PSM for Post-fire rehabilitation monitoring plots. The figure compares eCDFs for mean annual precipitation (top), elevation (middle), and sand texture (bottom) percentages between aerial seeding and non-treated controls, for both unmatched (left column) and matched sets of monitoring plots (right column).

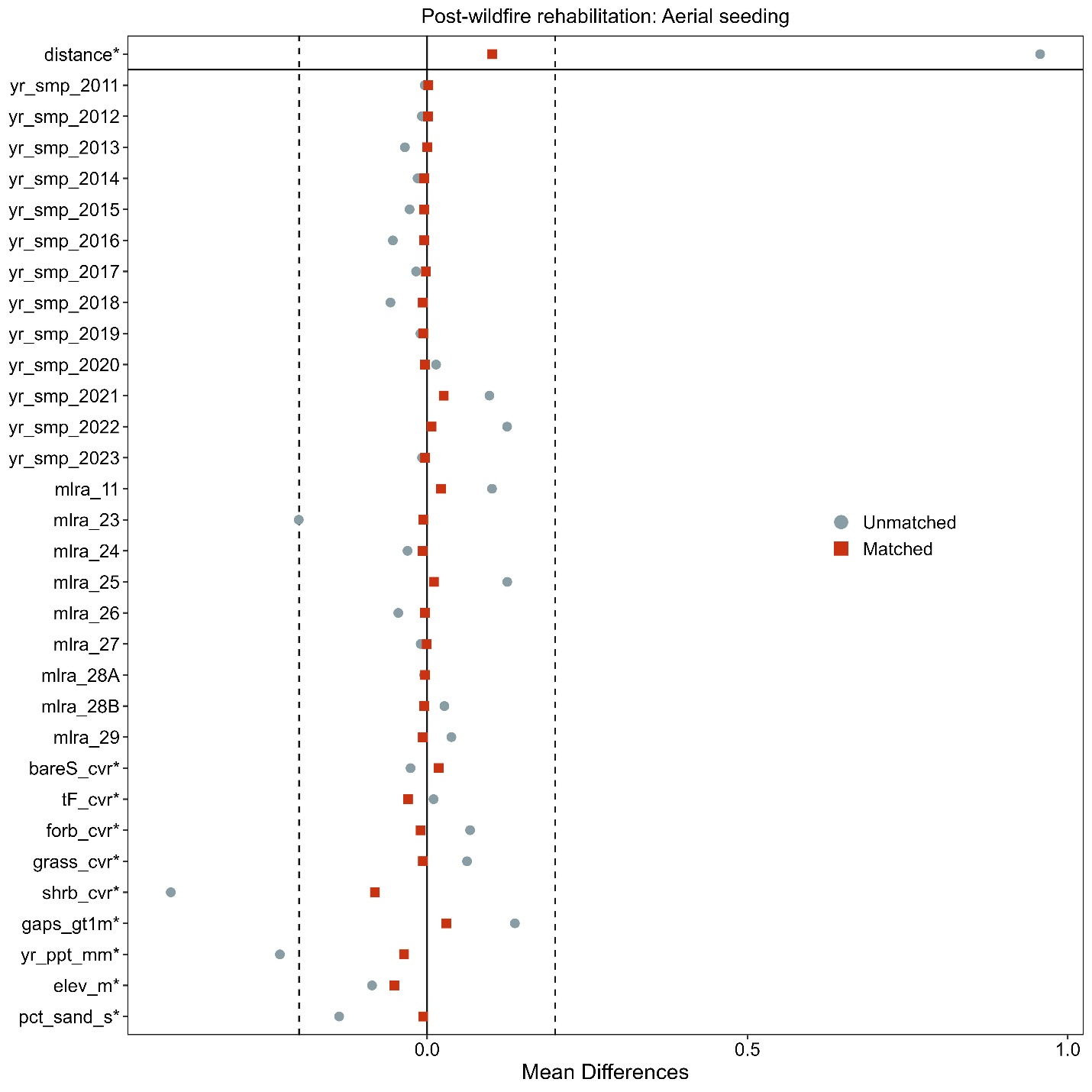

**Figure S9e.** Love-plot illustrating standardized mean differences for covariates in both the full dataset (unmatched) and the matched dataset for aerial seeding and non-treated controls for Post-fire restoration plots. Vertical lines indicate the caliper threshold (0.2 times the standard deviation of the propensity score) applied during matching.

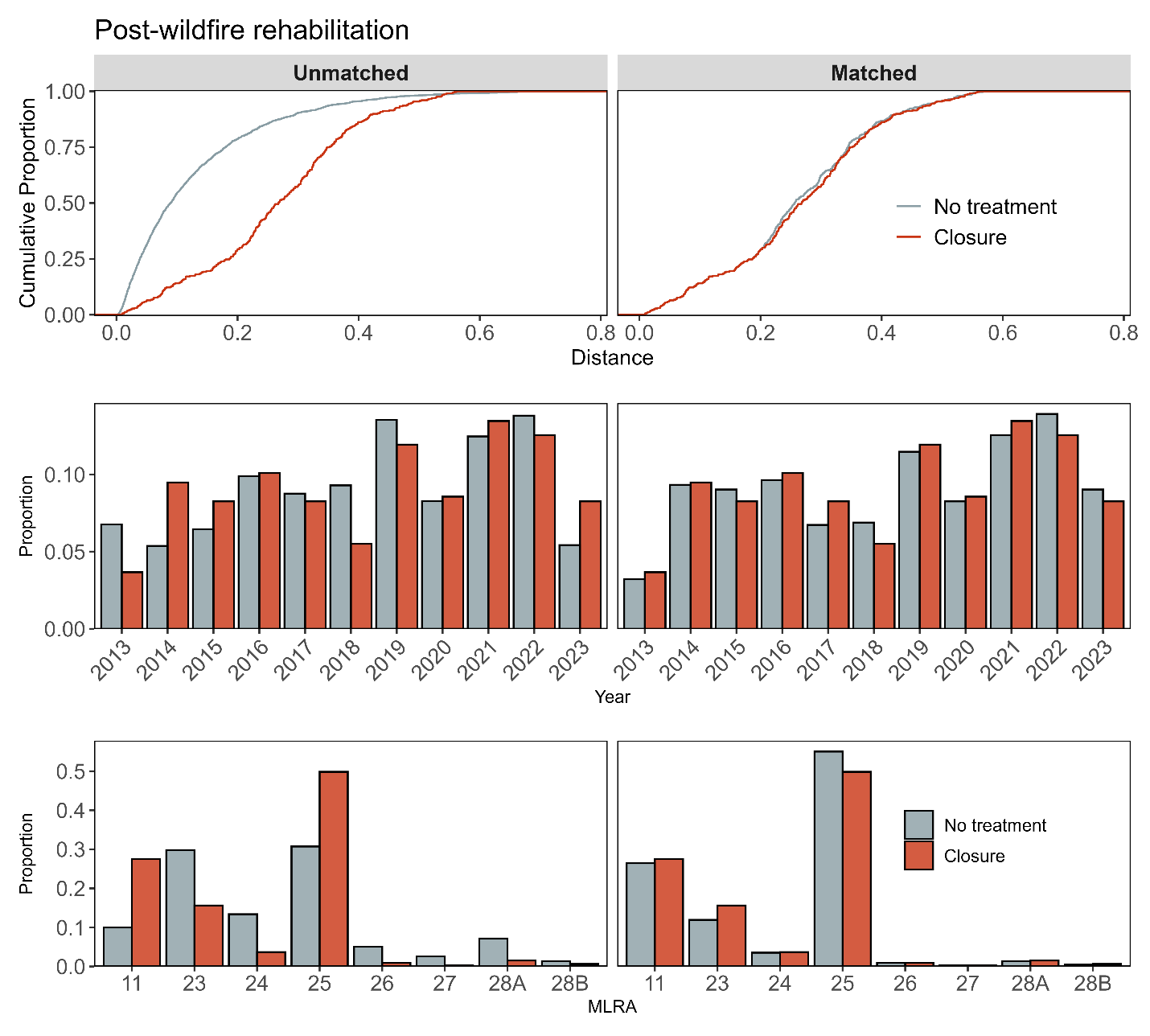

**Figure S10a.** Assessment of covariate balance using propensity score matching (PSM) for Post-fire rehabilitation monitoring plots. The top plot shows the empirical cumulative distribution function (eCDF) for the distance measure of the propensity score. The middle and bottom plots display balance in categorical variables, accounting for the year of monitoring and the major land resource area (MLRA) of the plots, comparing unmatched (left column) and matched sets (right columns) for closure and non-treated control monitoring plots.

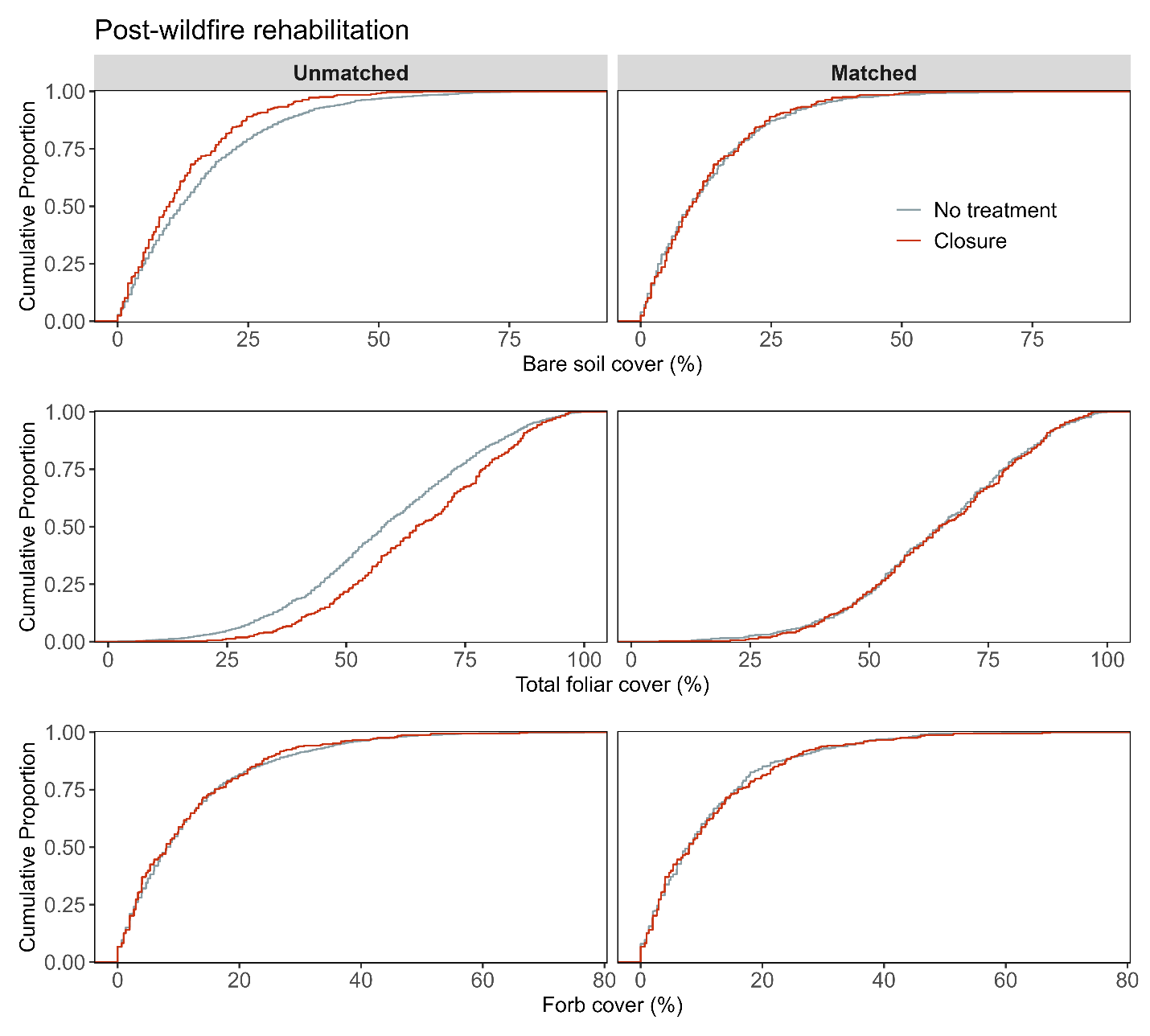

**Figure S10b**. Assessment of covariate balance using PSM for Post-fire rehabilitation monitoring plots. The figure compares eCDFs for bare soil (top), total foliar cover (middle), and forb cover (bottom) percentages between closure and non-treated controls, for both unmatched (left column) and matched sets of monitoring plots (right column).

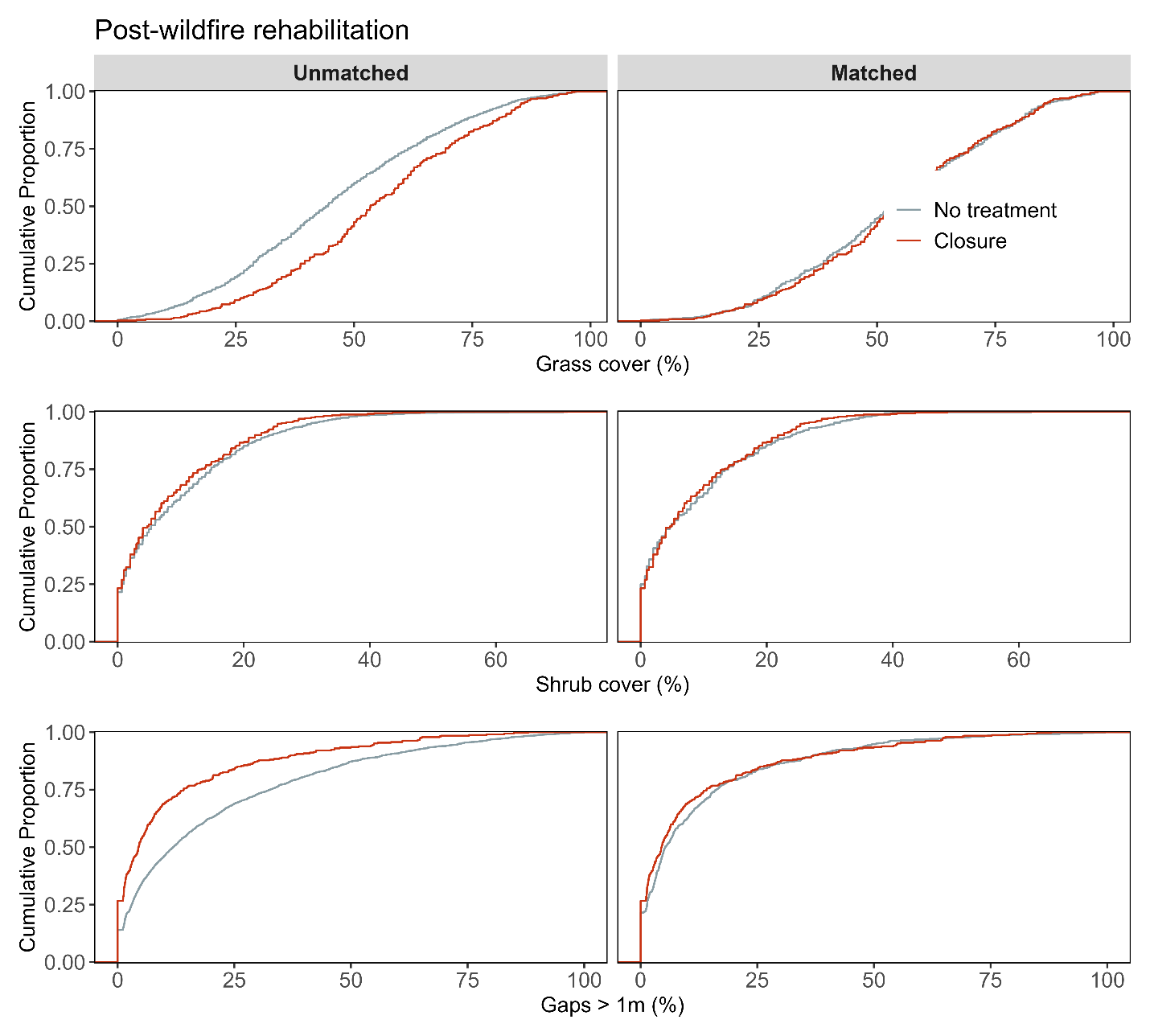

**Figure S10c**. Assessment of covariate balance using PSM for Post-fire rehabilitation monitoring plots. The figure compares eCDFs for grass cover (top), shrub cover (middle), and gaps > 1m (bottom) percentages between closure and non-treated controls, for both unmatched (left column) and matched sets of monitoring plots (right column).

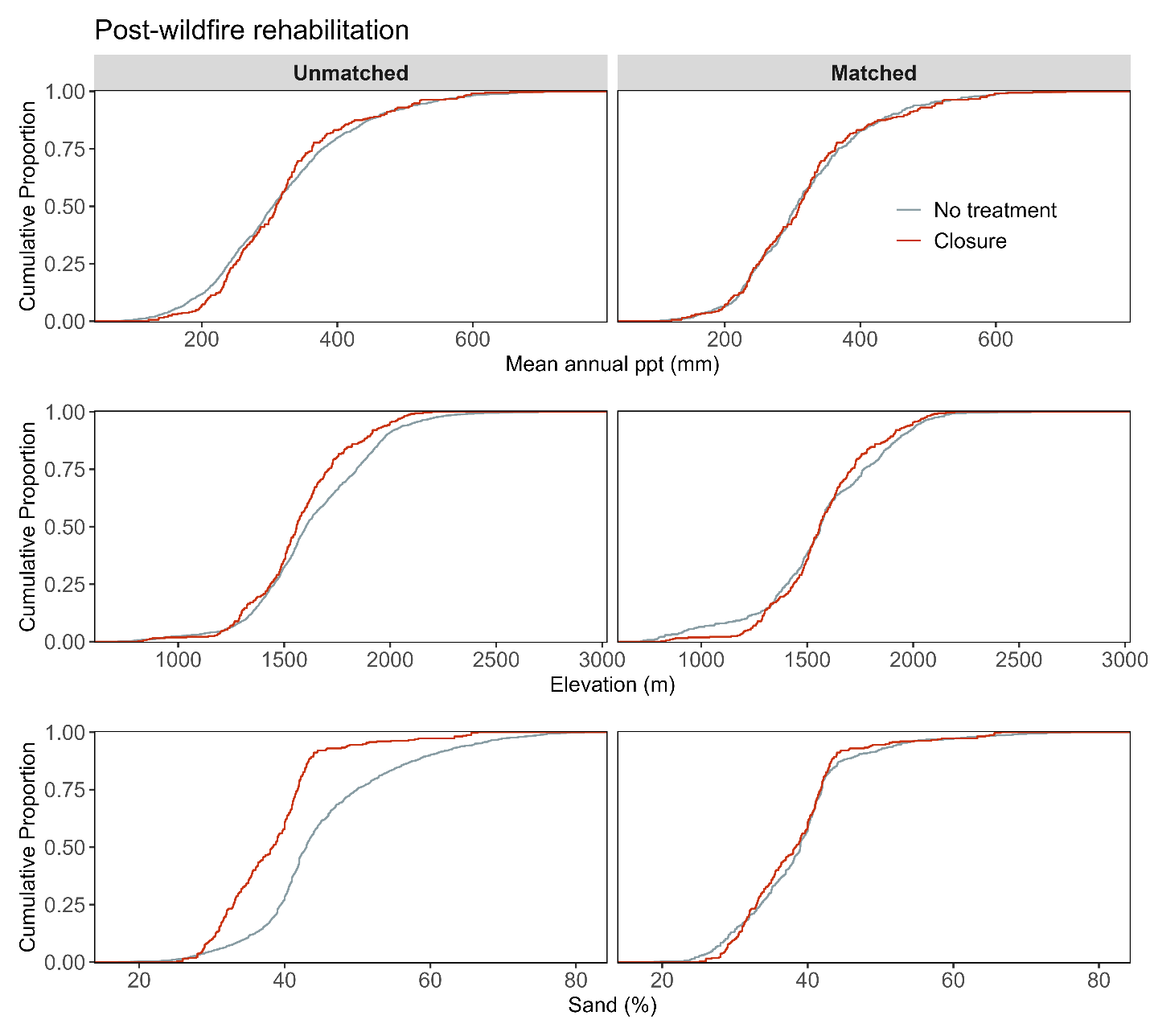

**Figure S10d**. Assessment of covariate balance using PSM for Post-fire rehabilitation monitoring plots. The figure compares eCDFs for mean annual precipitation (top), elevation (middle), and sand texture (bottom) percentages between closure and non-treated controls, for both unmatched (left column) and matched sets of monitoring plots (right column).

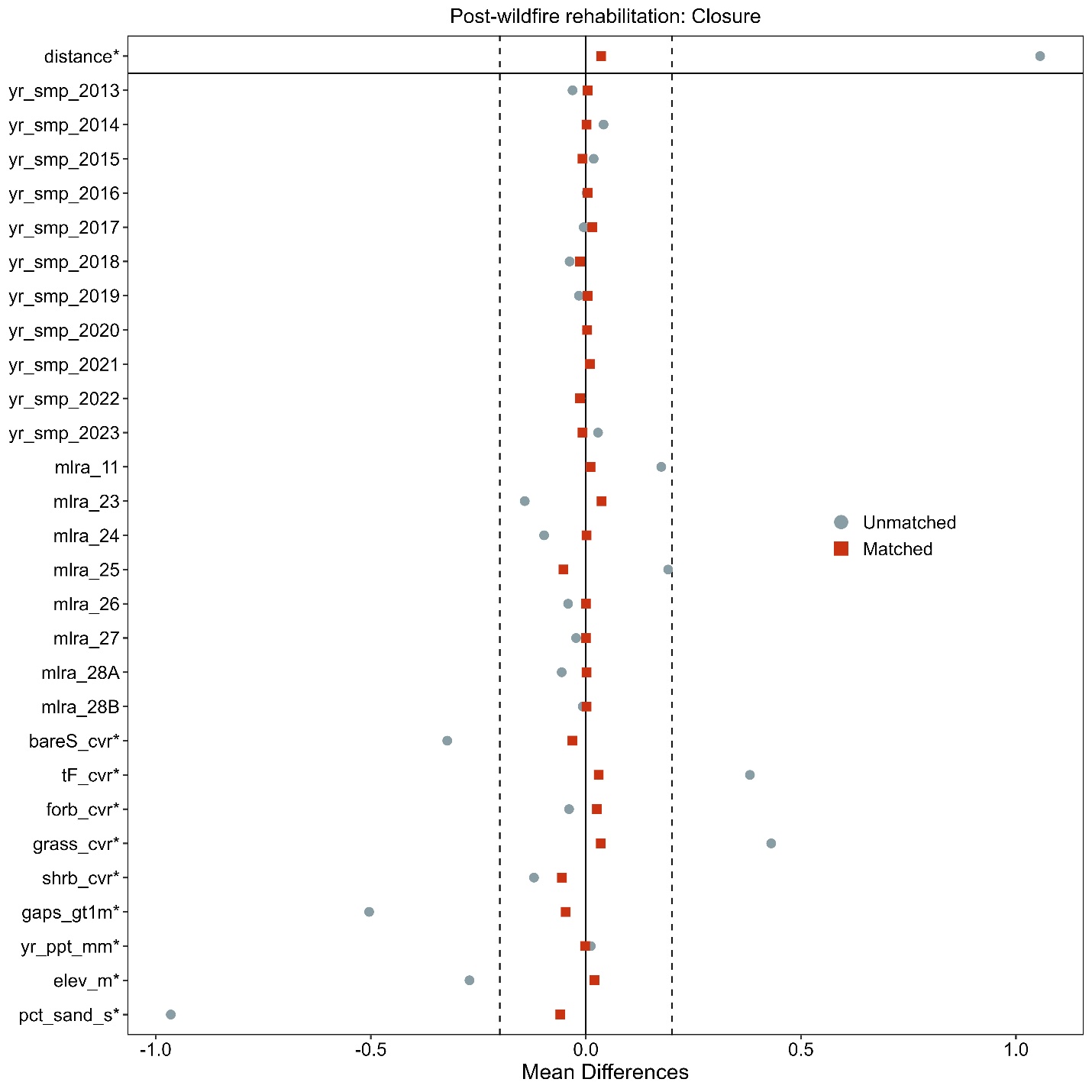

**Figure S10e**. Love-plot illustrating standardized mean differences for covariates in both the full dataset (unmatched) and the matched dataset for closure and non-treated controls Post-fire rehabilitation plots. Vertical lines indicate the caliper threshold (0.2 times the standard deviation of the propensity score) applied during matching.

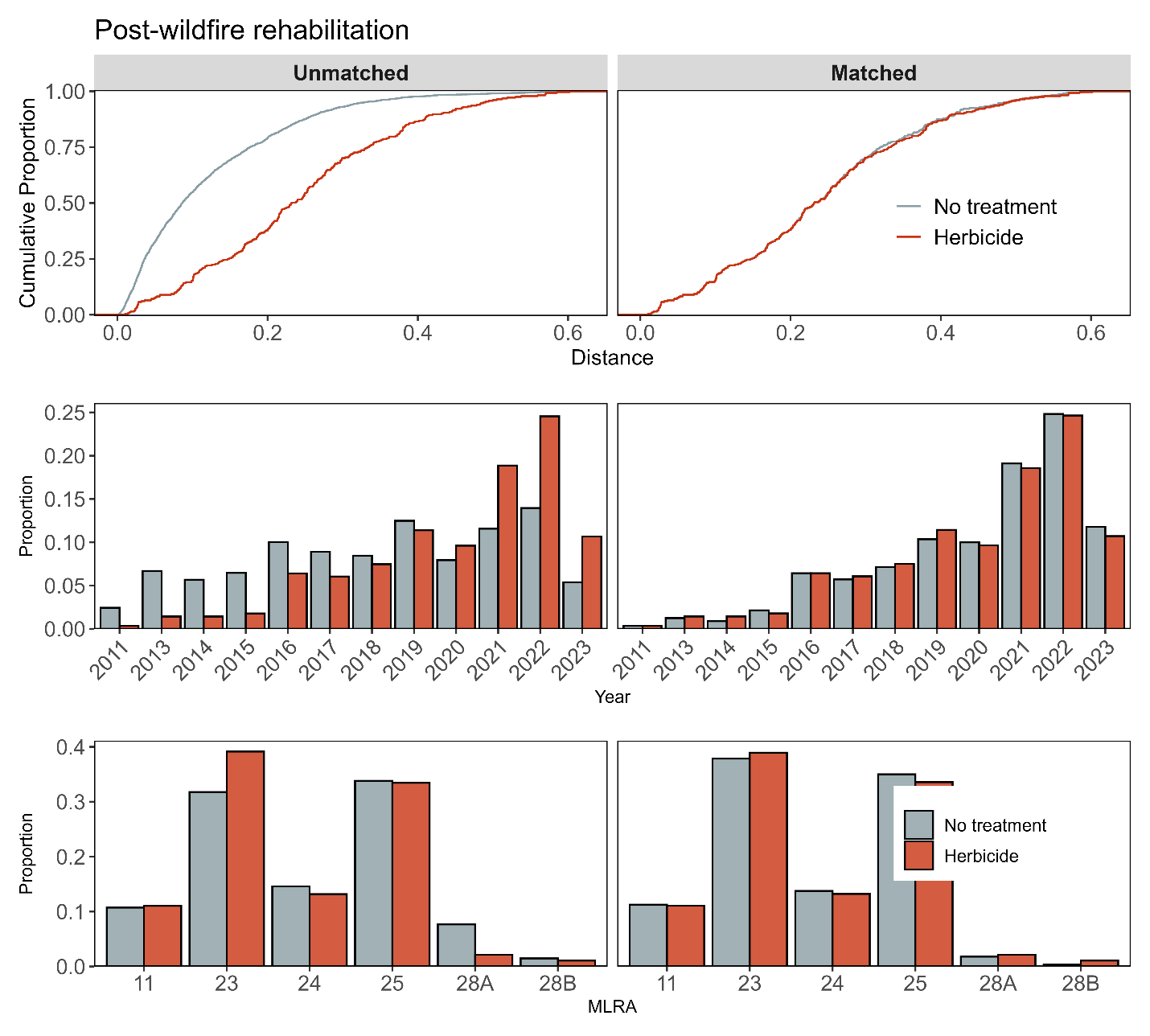

**Figure S11a**. Assessment of covariate balance using propensity score matching (PSM) for Post-fire rehabilitation monitoring plots. The top plot shows the empirical cumulative distribution function (eCDF) for the distance measure of the propensity score. The middle and bottom plots display balance in categorical variables, accounting for the year of monitoring and the major land resource area (MLRA) of the plots, comparing unmatched (left column) and matched sets (right columns) for herbicide and non-treated control monitoring plots.

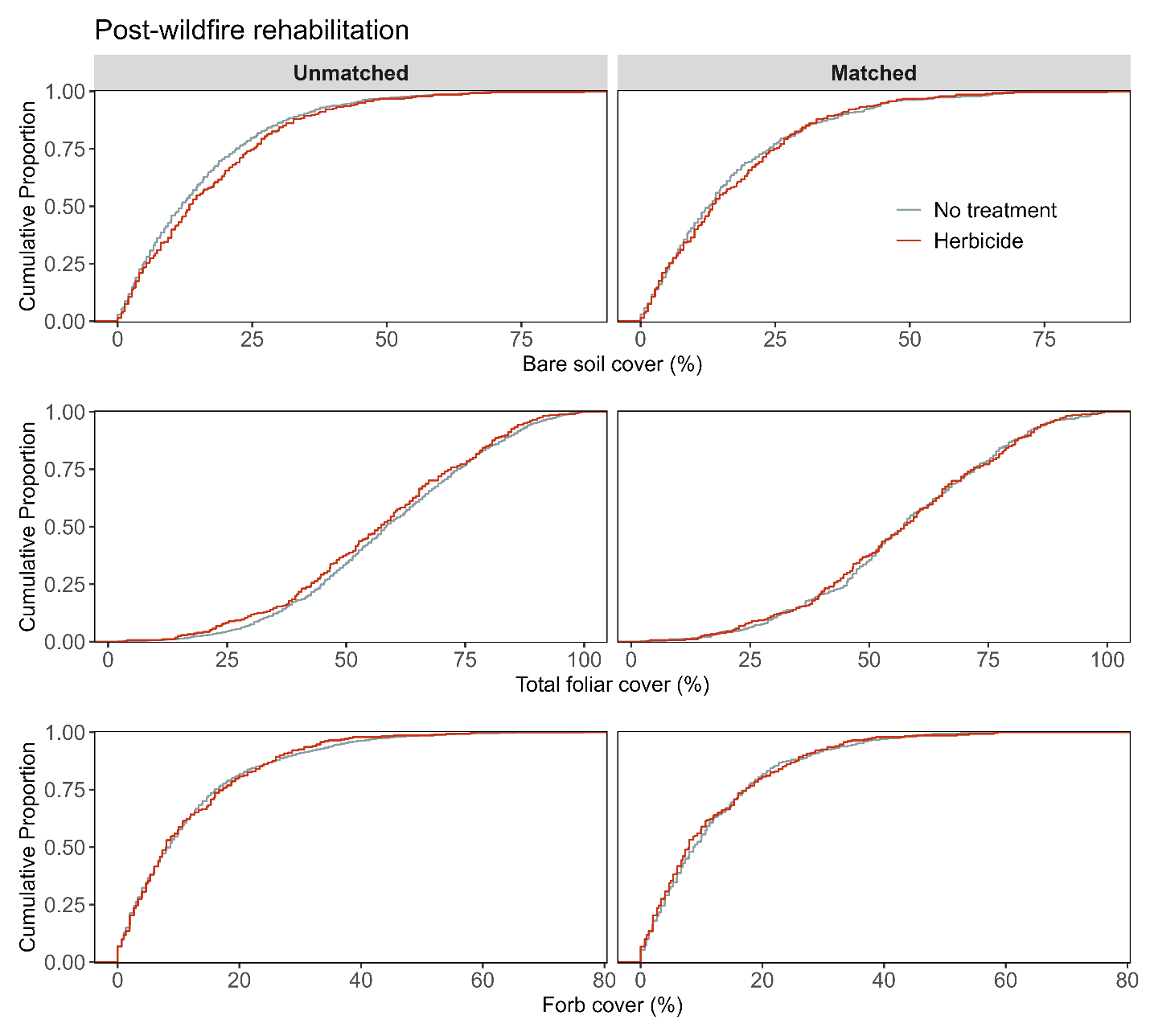

**Figure S11b**. Assessment of covariate balance using PSM for Post-fire rehabilitation monitoring plots. The figure compares eCDFs for bare soil (top), total foliar cover (middle), and forb cover (bottom) percentages between herbicide and non-treated controls, for both unmatched (left column) and matched sets of monitoring plots (right column).

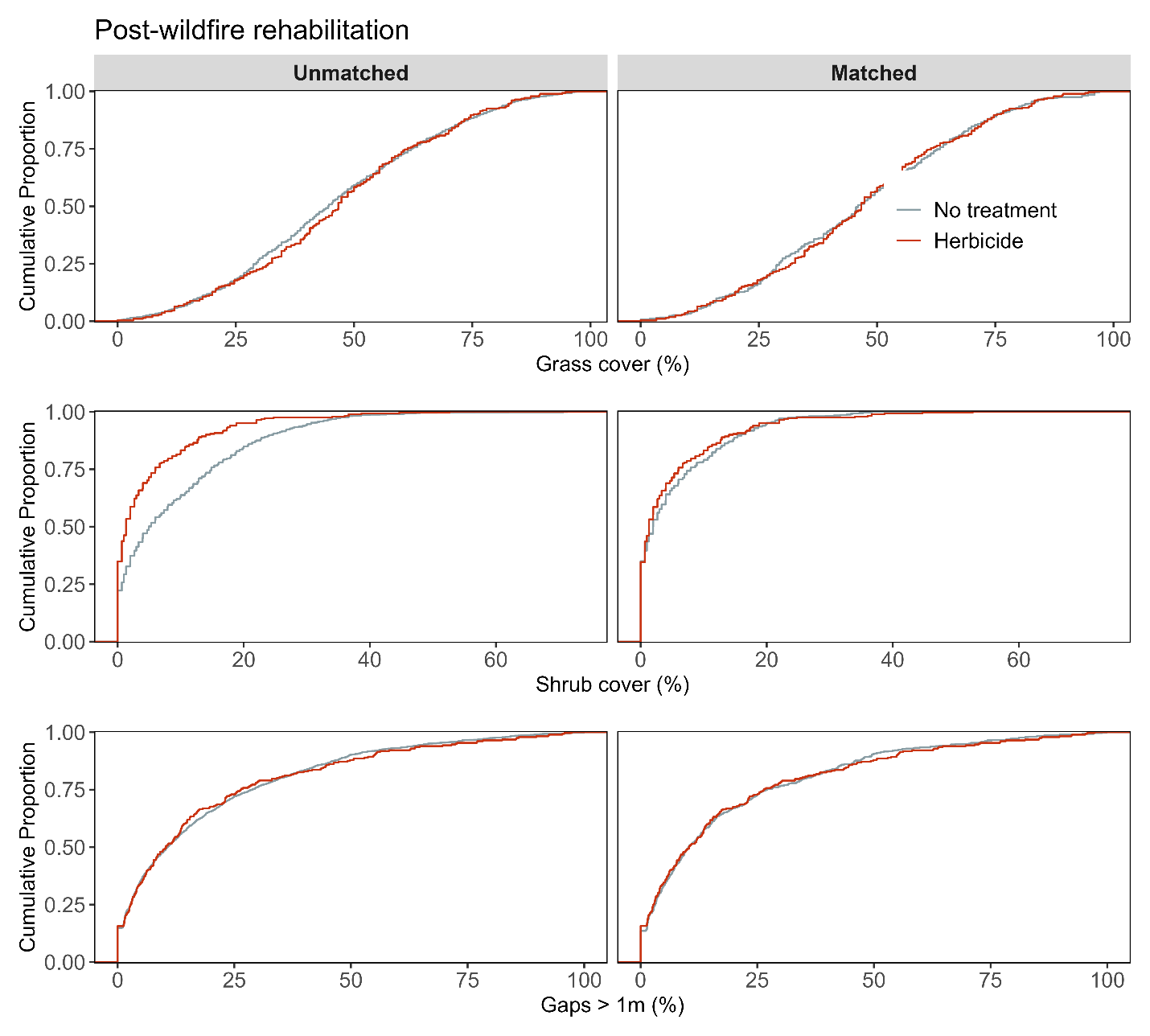

**Figure S11c**. Assessment of covariate balance using PSM for Post-fire rehabilitation monitoring plots. The figure compares eCDFs for grass cover (top), shrub cover (middle), and gaps > 1m (bottom) percentages between herbicide and non-treated controls, for both unmatched (left column) and matched sets of monitoring plots (right column).

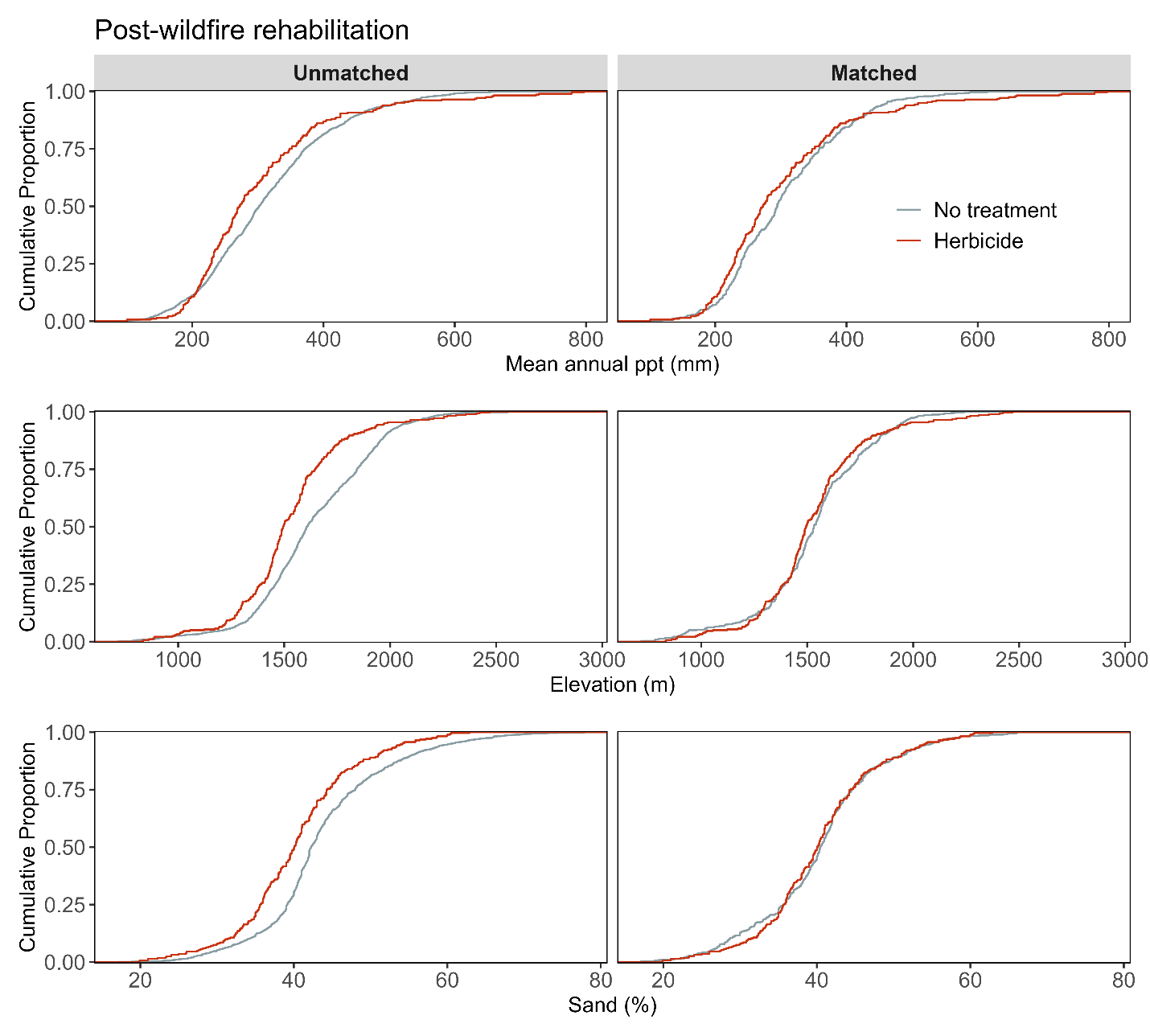

**Figure S4.11d**. Assessment of covariate balance using PSM for Post-fire rehabilitation monitoring plots. The figure compares eCDFs for mean annual precipitation (top), elevation (middle), and sand texture (bottom) percentages between herbicide and non-treated controls, for both unmatched (left column) and matched sets of monitoring plots (right column).

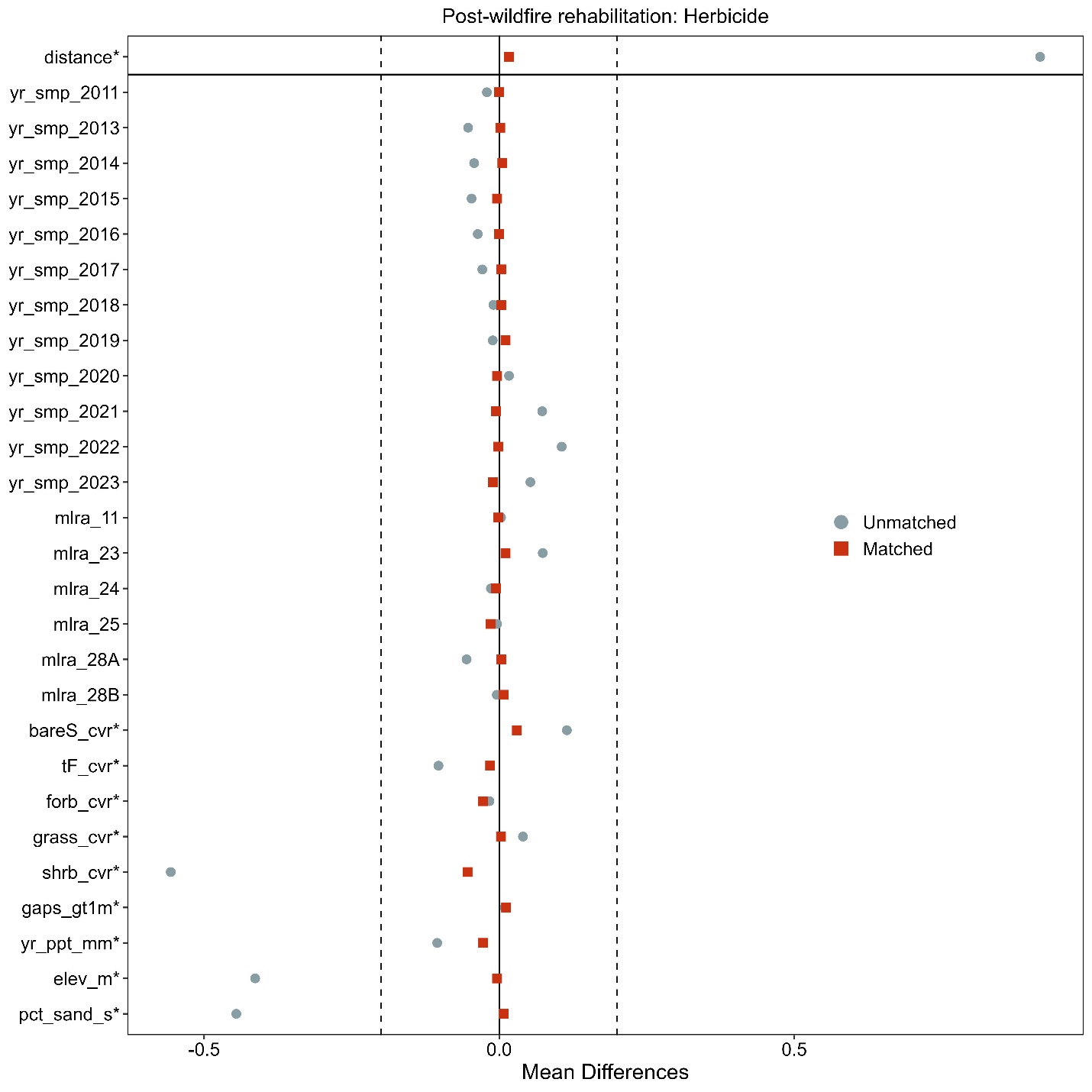

**Figure S11e**. Love-plot illustrating standardized mean differences for covariates in both the full dataset (unmatched) and the matched dataset for herbicide and non-treated controls Post-fire rehabilitation plots. Vertical lines indicate the caliper threshold (0.2 times the standard deviation of the propensity score) applied during matching.

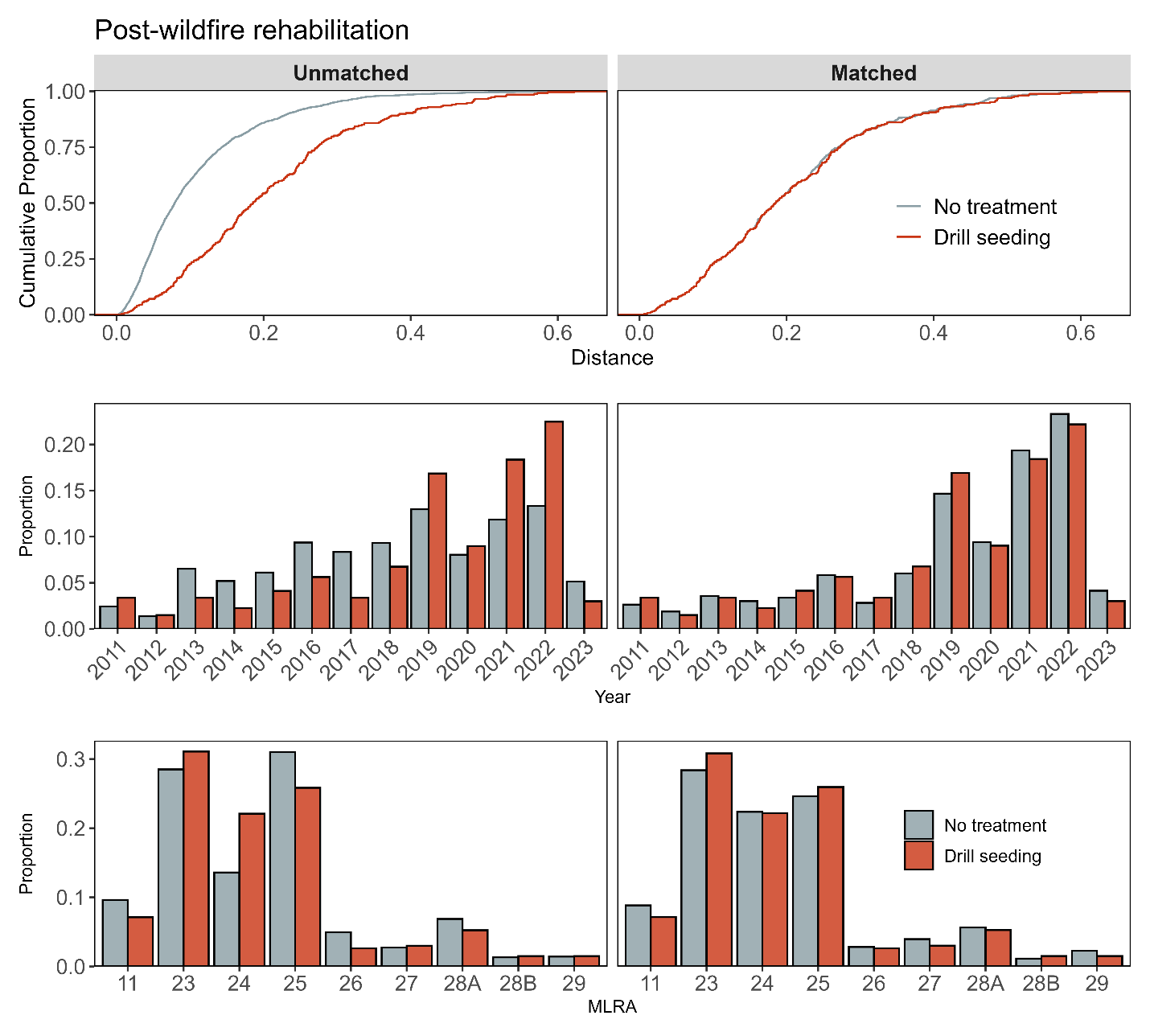

**Figure S12a**. Assessment of covariate balance using propensity score matching (PSM) for Post-fire rehabilitation monitoring plots. The top plot shows the empirical cumulative distribution function (eCDF) for the distance measure of the propensity score. The middle and bottom plots display balance in categorical variables, accounting for the year of monitoring and the major land resource area (MLRA) of the plots, comparing unmatched (left column) and matched sets (right columns) for drill seeding and non-treated control monitoring plots.

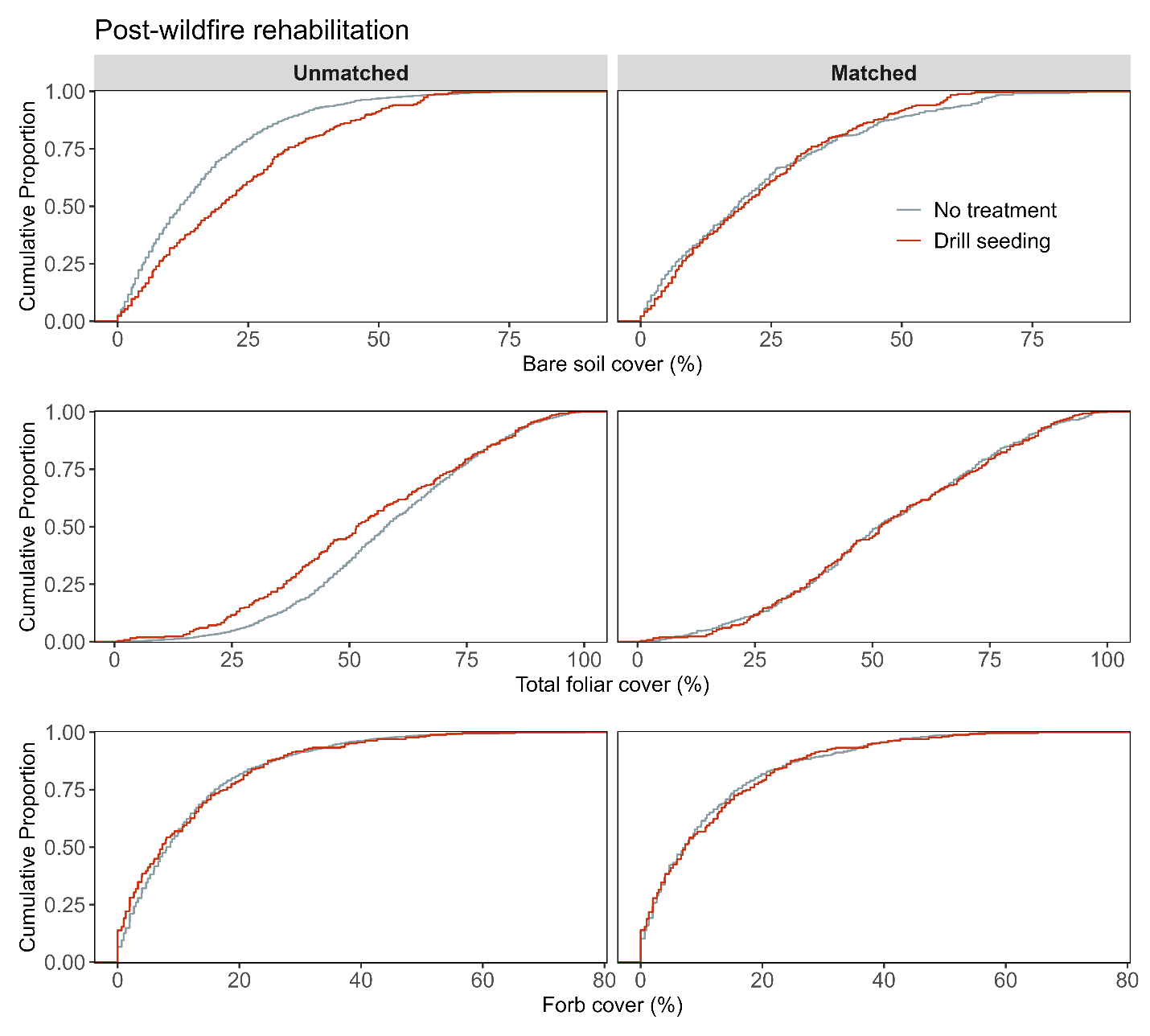

**Figure S12b**. Assessment of covariate balance using PSM for Post-fire rehabilitation monitoring plots. The figure compares eCDFs for bare soil (top), total foliar cover (middle), and forb cover (bottom) percentages between drill seeding and non-treated controls, for both unmatched (left column) and matched sets of monitoring plots (right column).

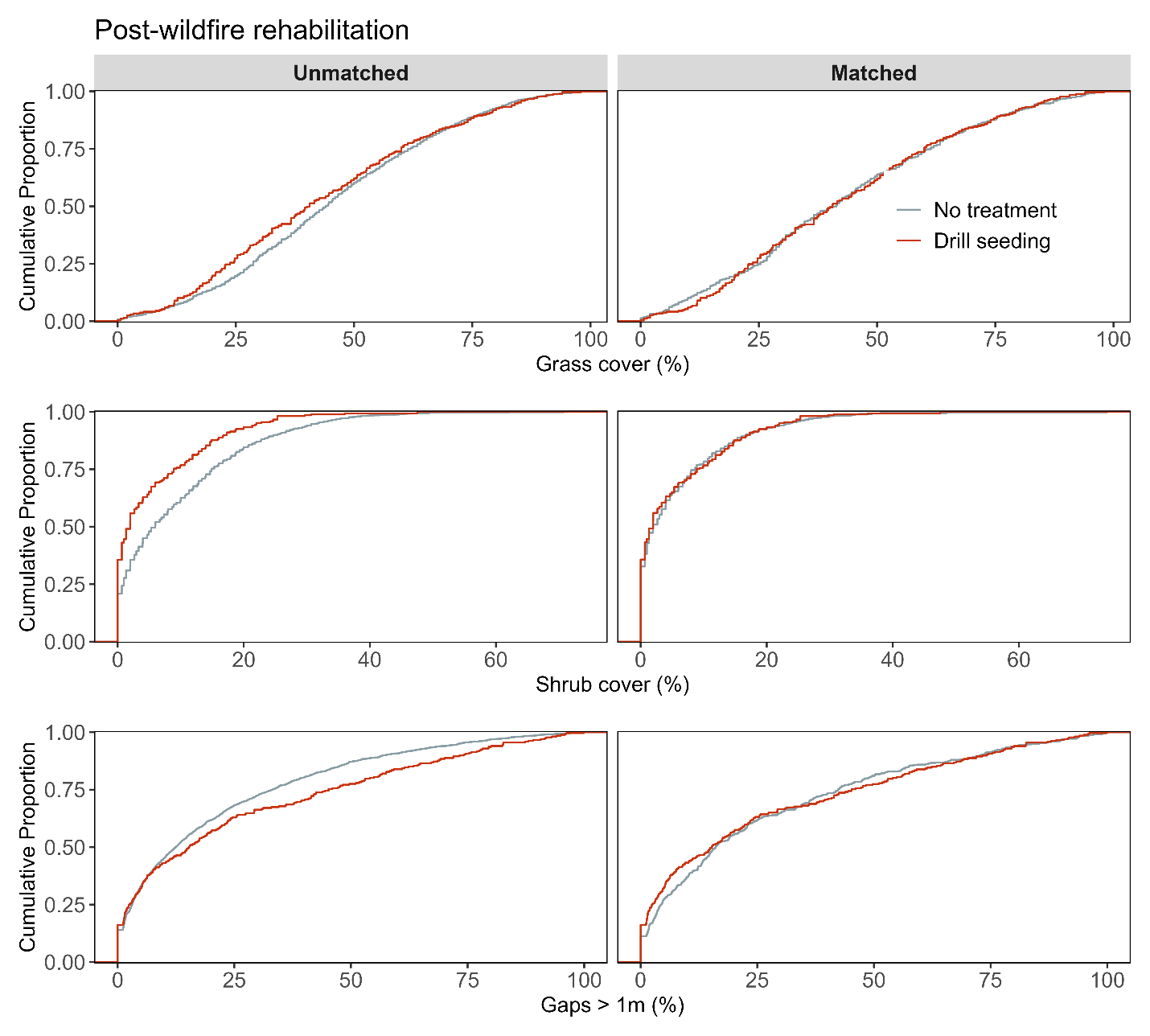

**Figure S12c**. Assessment of covariate balance using PSM for Post-fire rehabilitation monitoring plots. The figure compares eCDFs for grass cover (top), shrub cover (middle), and gaps > 1m (bottom) percentages between drill seeding and non-treated controls, for both unmatched (left column) and matched sets of monitoring plots (right column).

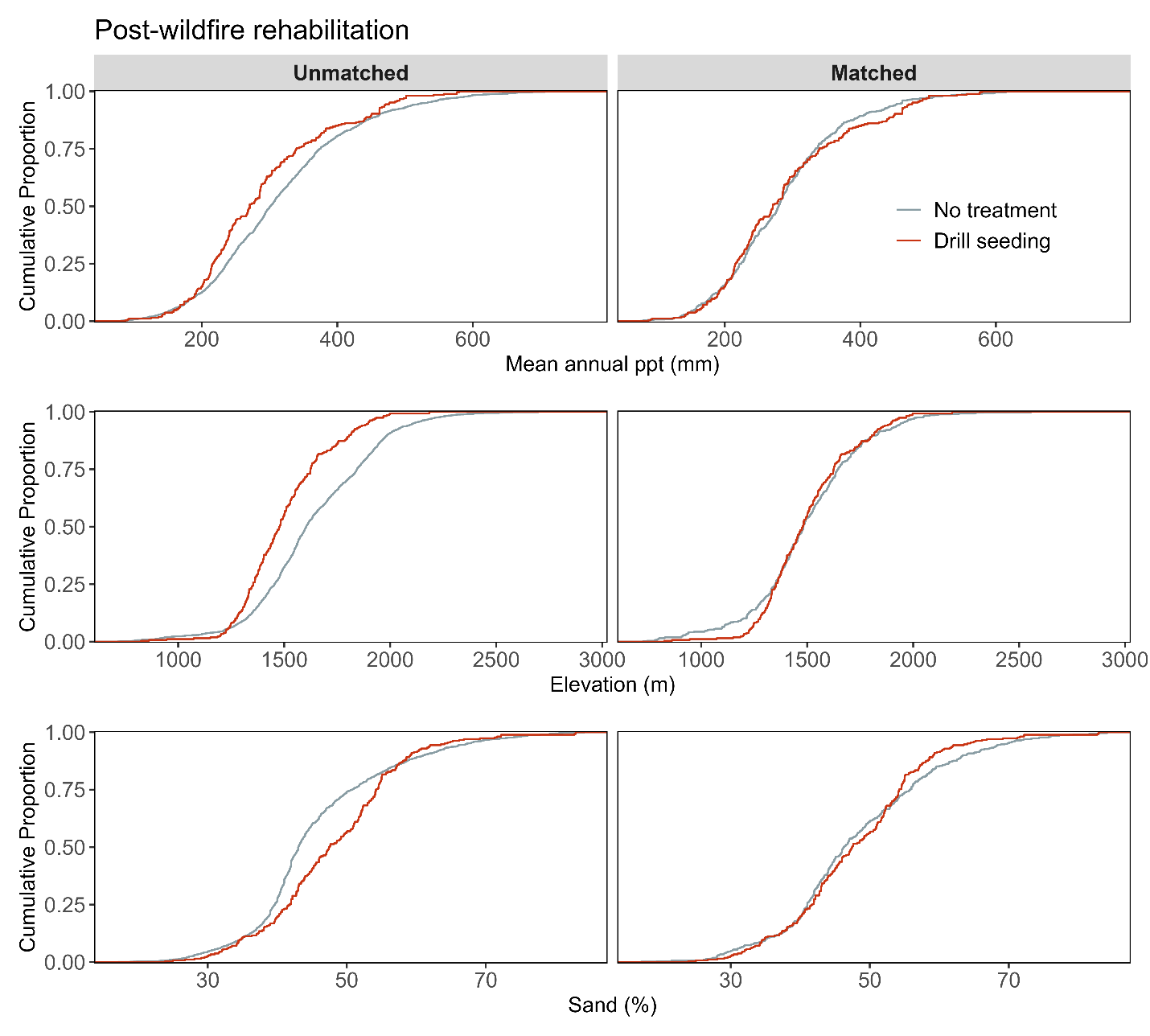

**Figure S12d.** Assessment of covariate balance using PSM for Post-fire rehabilitation monitoring plots. The figure compares eCDFs for mean annual precipitation (top), elevation (middle), and sand texture (bottom) percentages between drill seeding and non-treated controls, for both unmatched (left column) and matched sets of monitoring plots (right column).

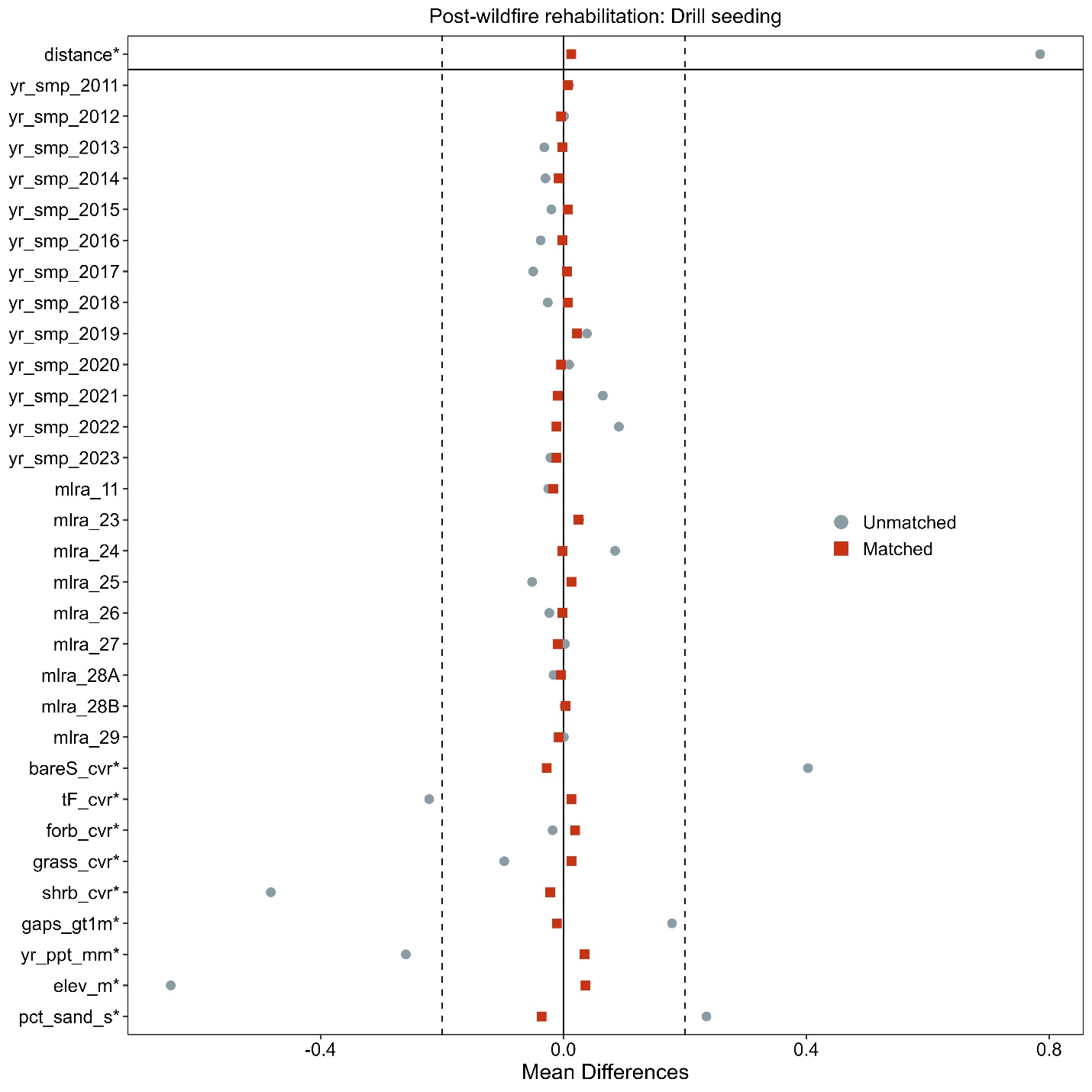

**Figure S12e**. Love-plot illustrating standardized mean differences for covariates in both the full dataset (unmatched) and the matched dataset for drill seeding and non-treated controls Post-fire rehabilitation plots. Vertical lines indicate the caliper threshold (0.2 times the standard deviation of the propensity score) applied during matching.

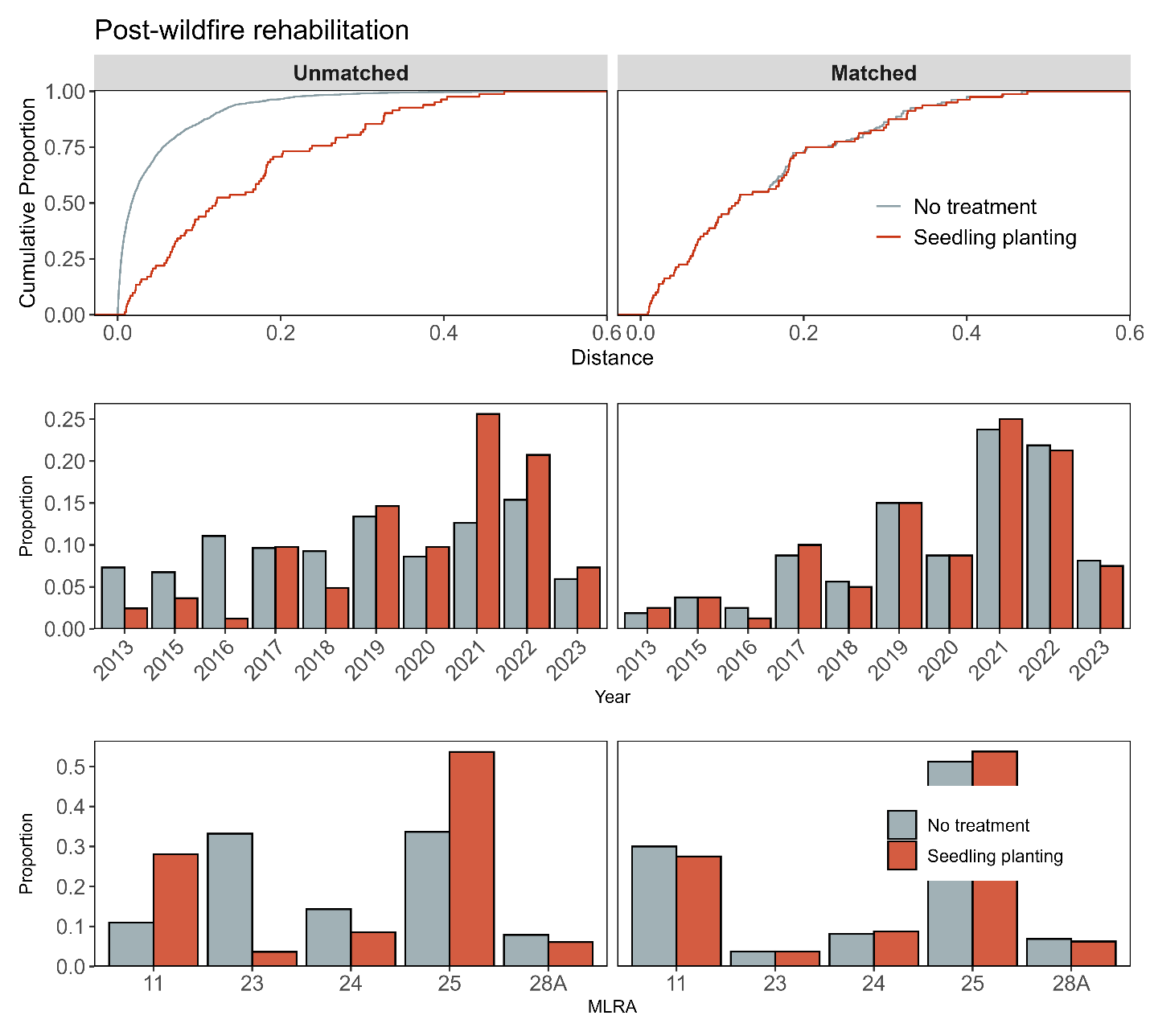

**Figure S13a**. Assessment of covariate balance using propensity score matching (PSM) for Post-fire rehabilitation monitoring plots. The top plot shows the empirical cumulative distribution function (eCDF) for the distance measure of the propensity score. The middle and bottom plots display balance in categorical variables, accounting for the year of monitoring and the major land resource area (MLRA) of the plots, comparing unmatched (left column) and matched sets (right columns) for seedling planting and non-treated control monitoring plots.

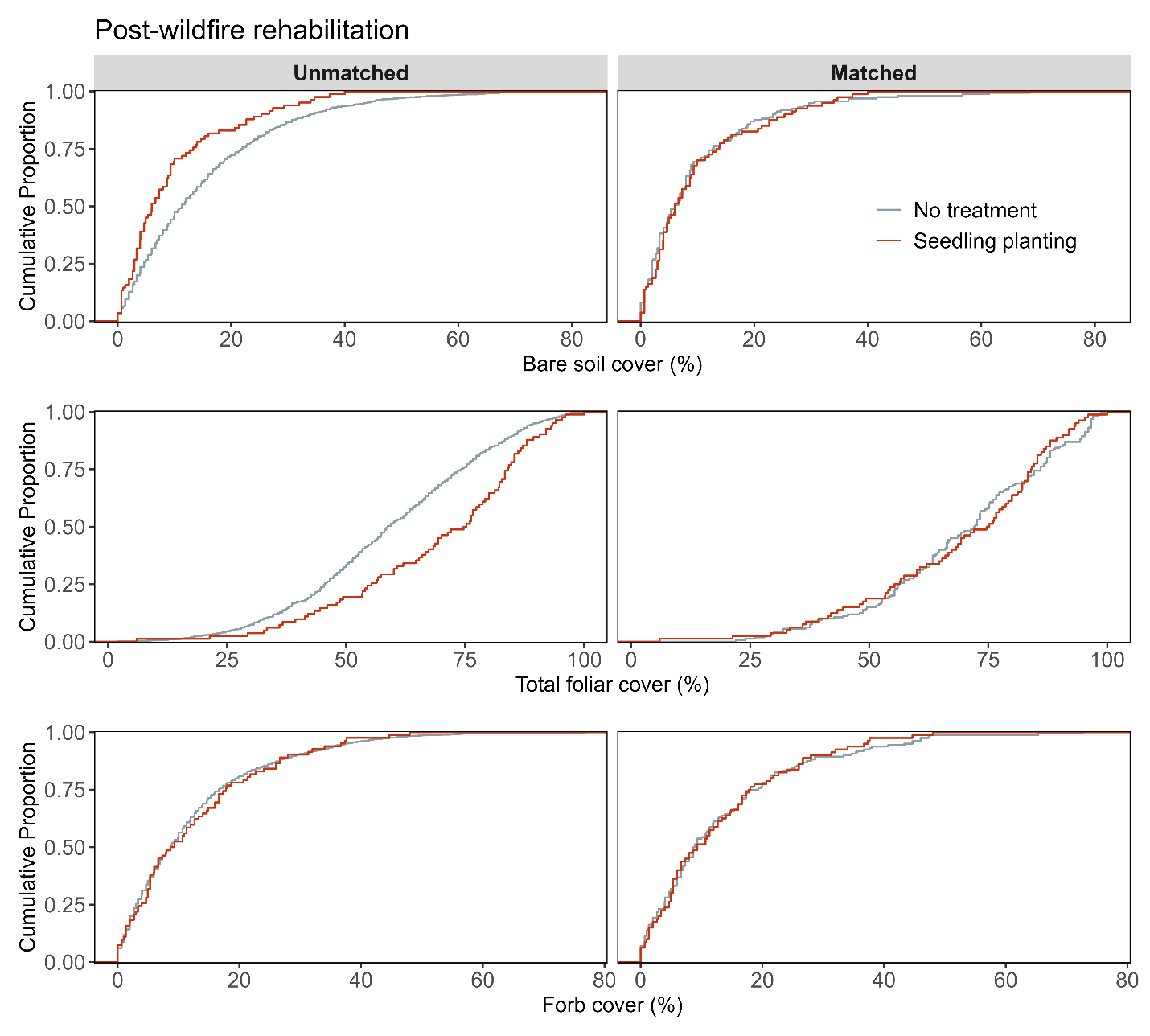

**Figure S13b**. Assessment of covariate balance using PSM for Post-fire rehabilitation monitoring plots. The figure compares eCDFs for bare soil (top), total foliar cover (middle), and forb cover (bottom) percentages between seedling planting and non-treated controls, for both unmatched (left column) and matched sets of monitoring plots (right column).

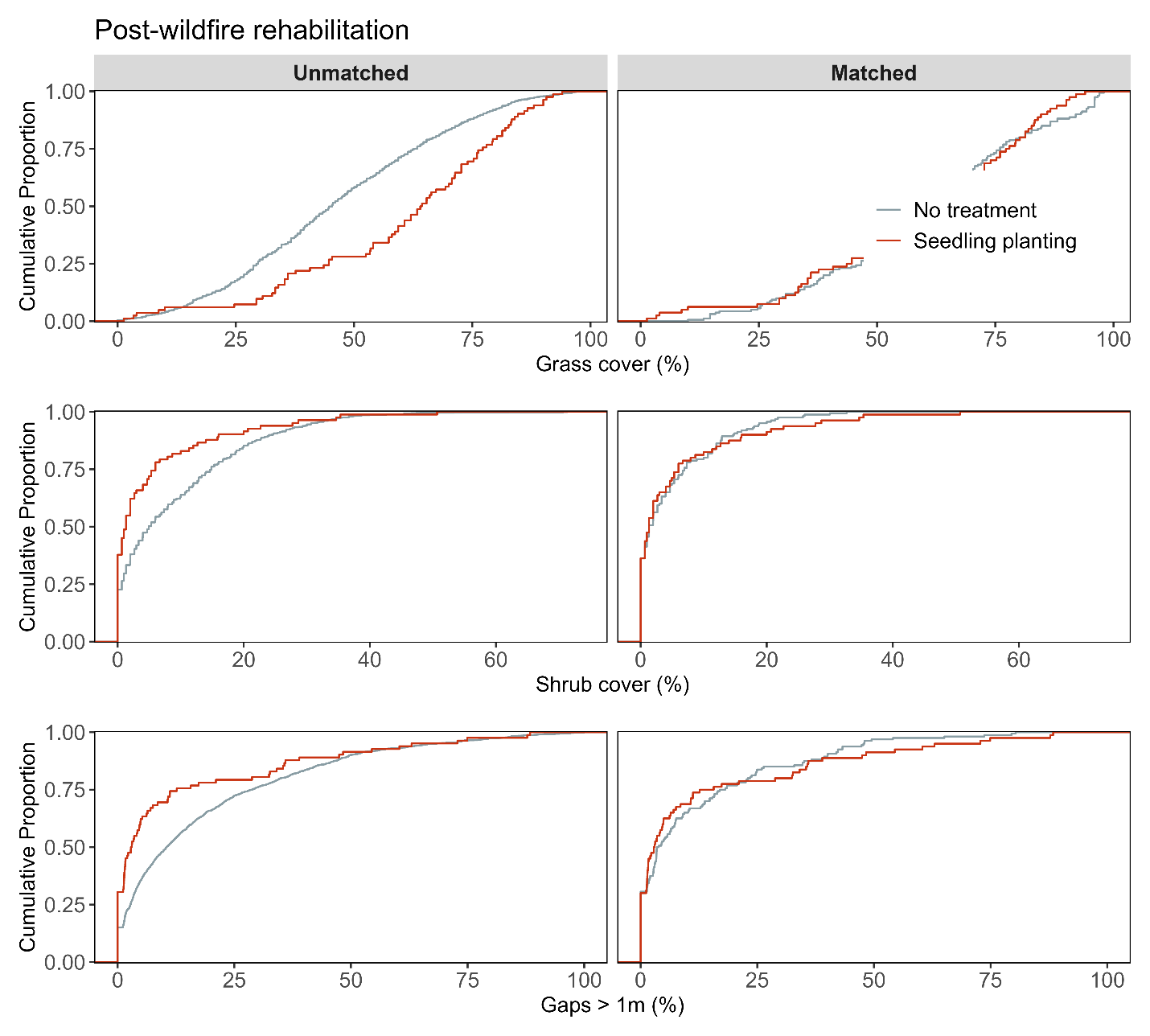

**Figure S13c**. Assessment of covariate balance using PSM for Post-fire rehabilitation monitoring plots. The figure compares eCDFs for grass cover (top), shrub cover (middle), and gaps > 1m (bottom) percentages between seedling planting and non-treated controls, for both unmatched (left column) and matched sets of monitoring plots (right column).

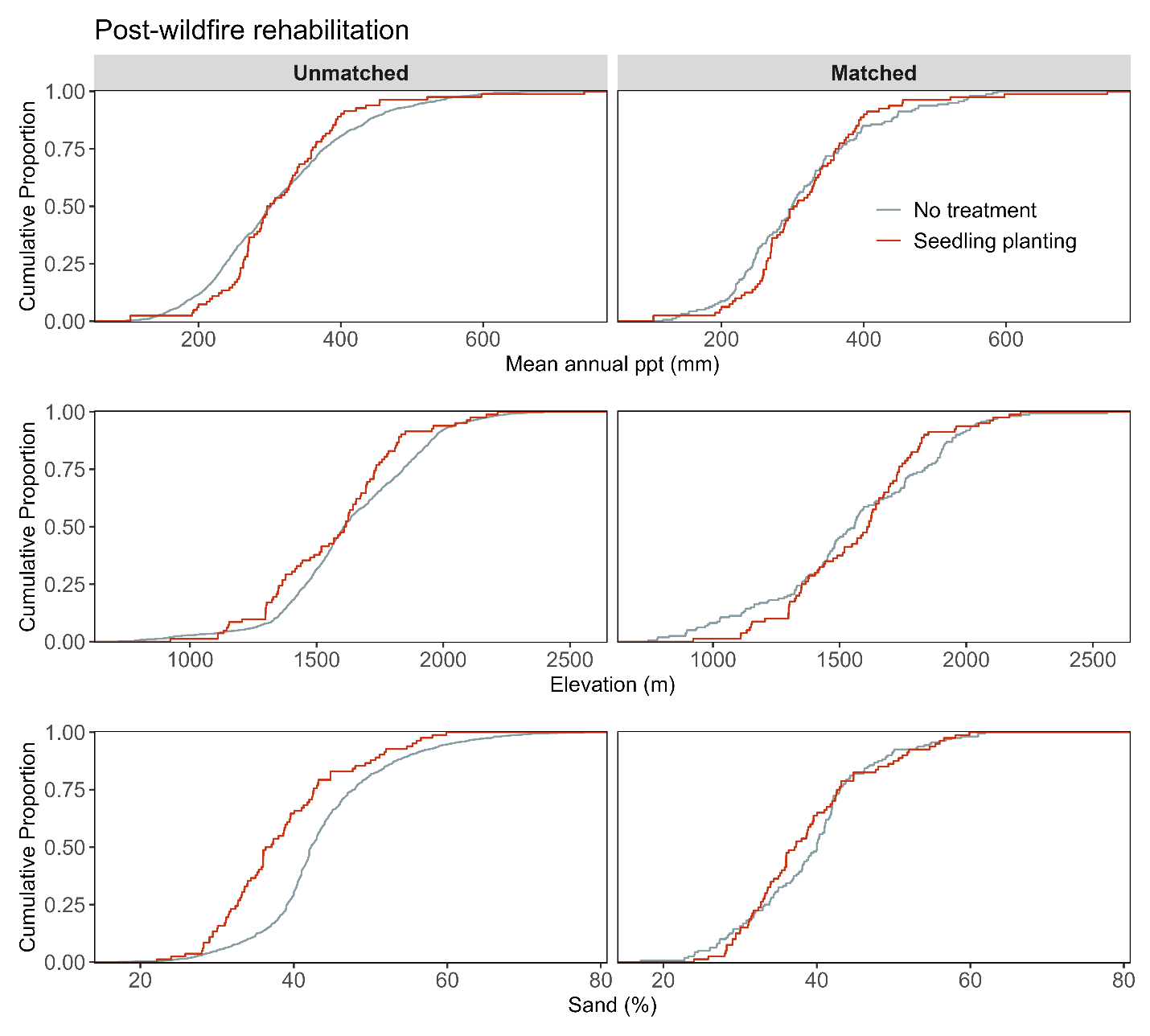

**Figure S13d.** Assessment of covariate balance using PSM for Post-fire rehabilitation monitoring plots. The figure compares eCDFs for mean annual precipitation (top), elevation (middle), and sand texture (bottom) percentages between seedling planting and non-treated controls, for both unmatched (left column) and matched sets of monitoring plots (right column).

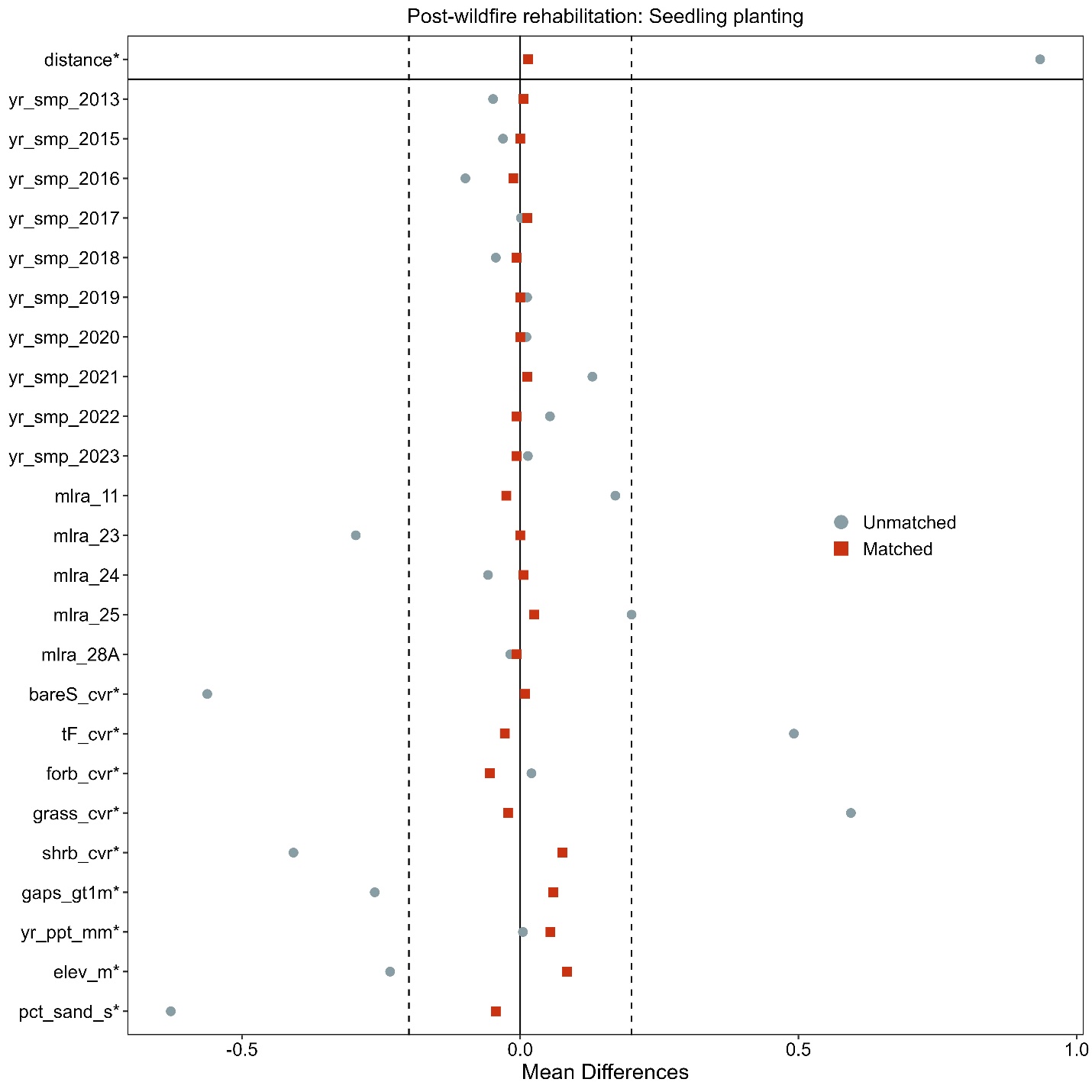

**Figure S13e**. Love-plot illustrating standardized mean differences for covariates in both the full dataset (unmatched) and the matched dataset for seedling planting and non-treated controls Post-fire rehabilitation plots. Vertical lines indicate the caliper threshold (0.2 times the standard deviation of the propensity score) applied during matching.

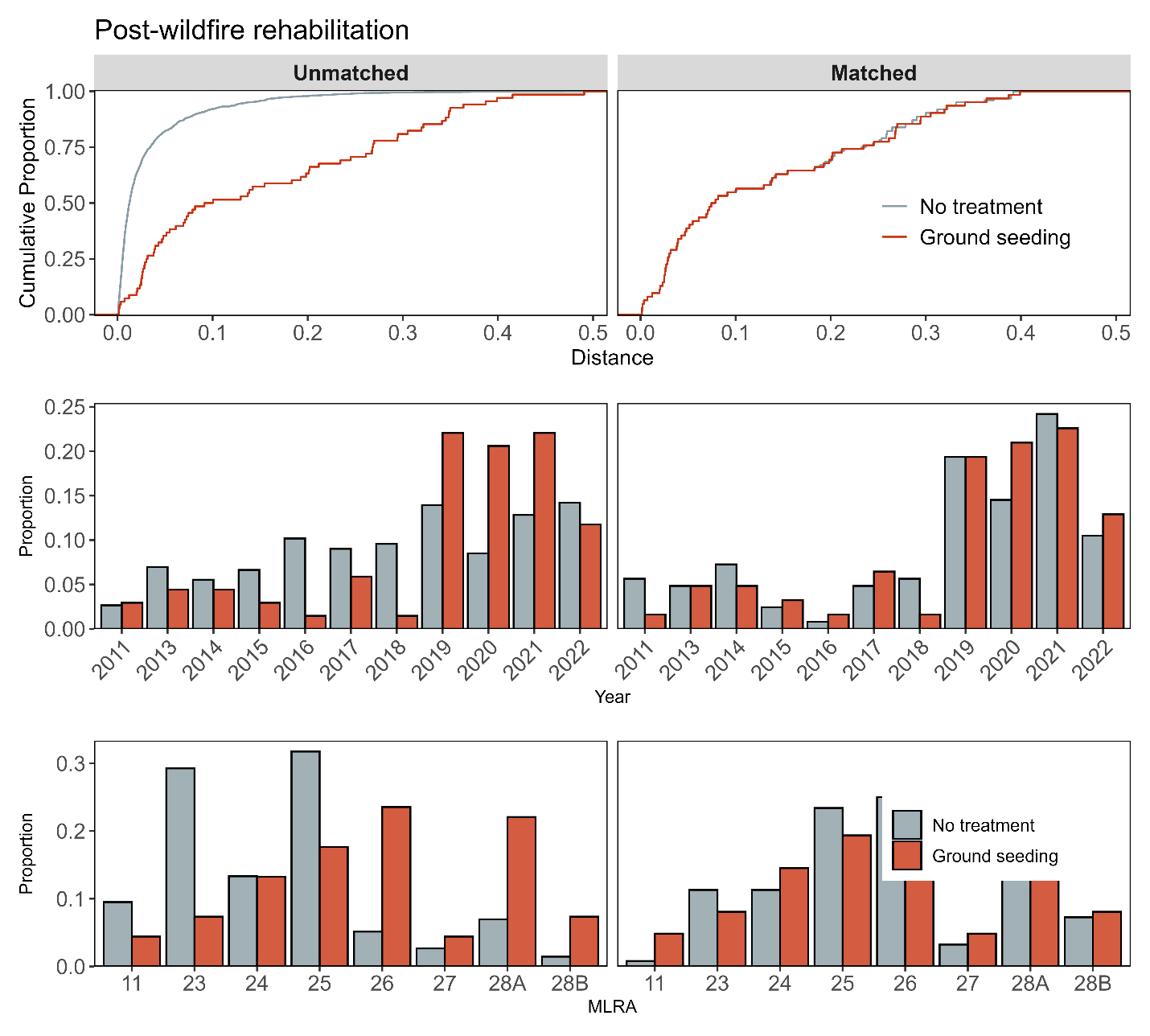

**Figure S14a**. Assessment of covariate balance using propensity score matching (PSM) for Post-fire rehabilitation monitoring plots. The top plot shows the empirical cumulative distribution function (eCDF) for the distance measure of the propensity score. The middle and bottom plots display balance in categorical variables, accounting for the year of monitoring and the major land resource area (MLRA) of the plots, comparing unmatched (left column) and matched sets (right columns) for ground seeding and non-treated control monitoring plots.

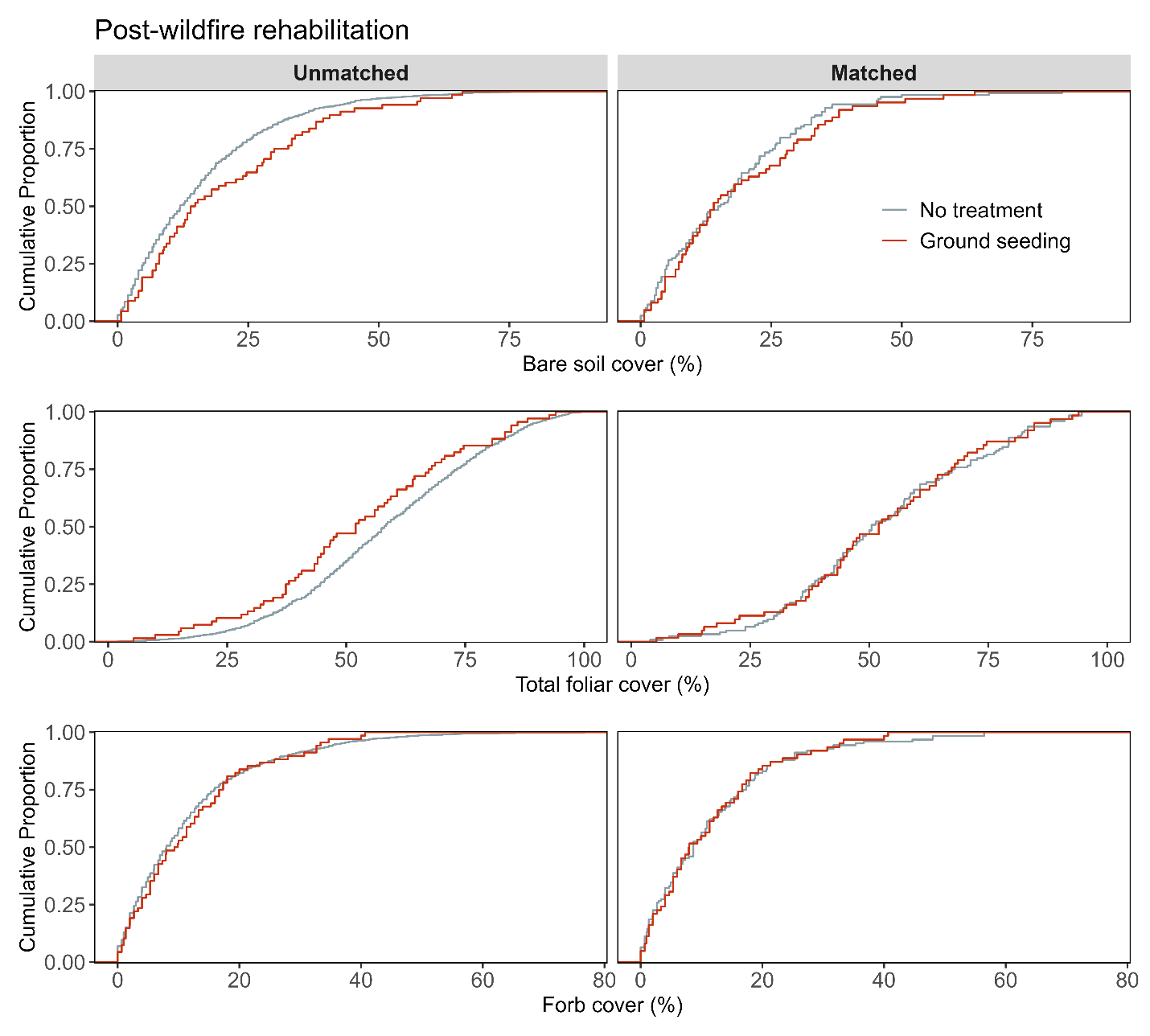

**Figure S14b**. Assessment of covariate balance using PSM for Post-fire rehabilitation monitoring plots. The figure compares eCDFs for bare soil (top), total foliar cover (middle), and forb cover (bottom) percentages between ground seeding and non-treated controls, for both unmatched (left column) and matched sets of monitoring plots (right column).

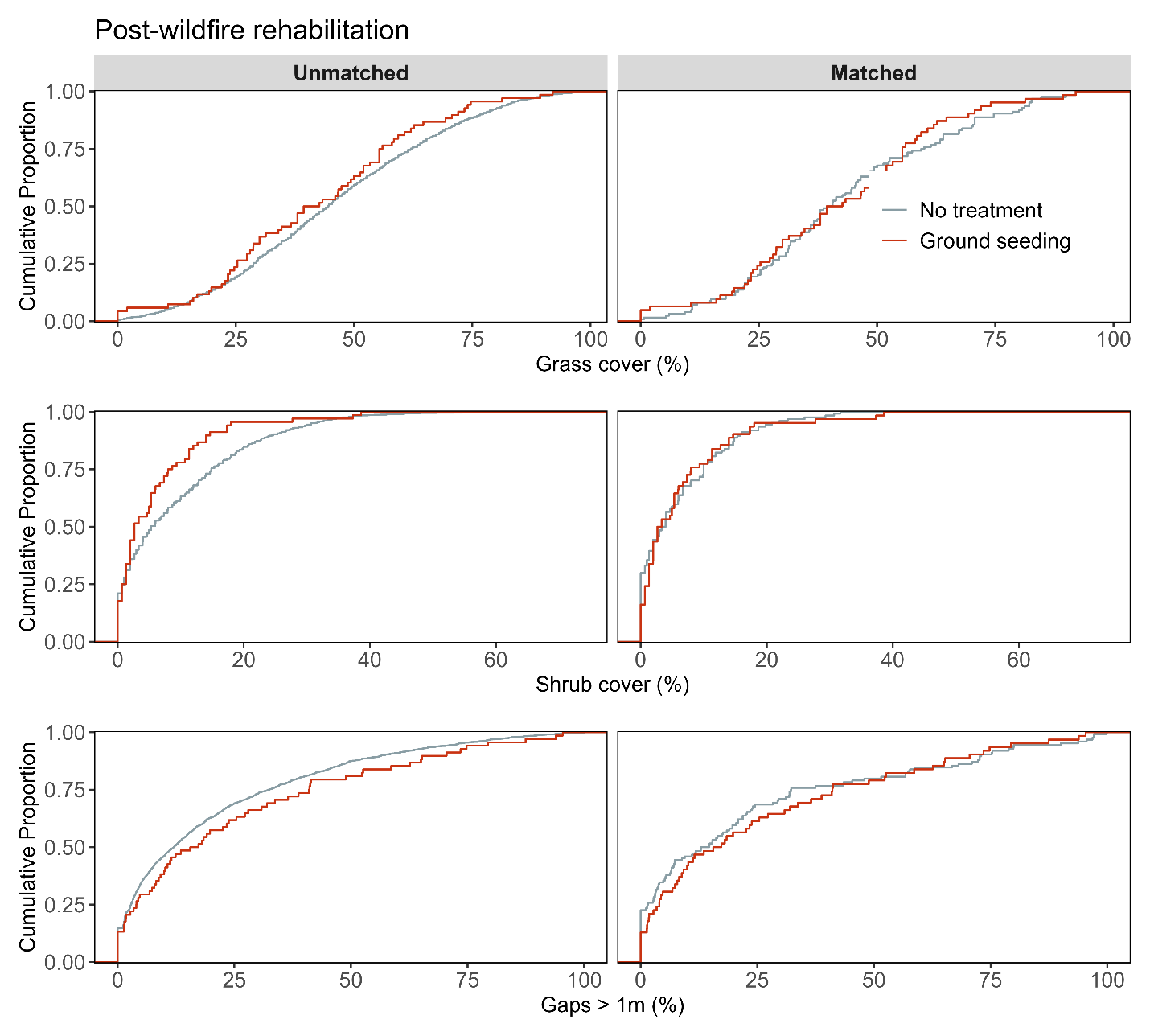

**Figure S14c**. Assessment of covariate balance using PSM for Post-fire rehabilitation monitoring plots. The figure compares eCDFs for grass cover (top), shrub cover (middle), and gaps > 1m (bottom) percentages between ground seeding and non-treated controls, for both unmatched (left column) and matched sets of monitoring plots (right column).

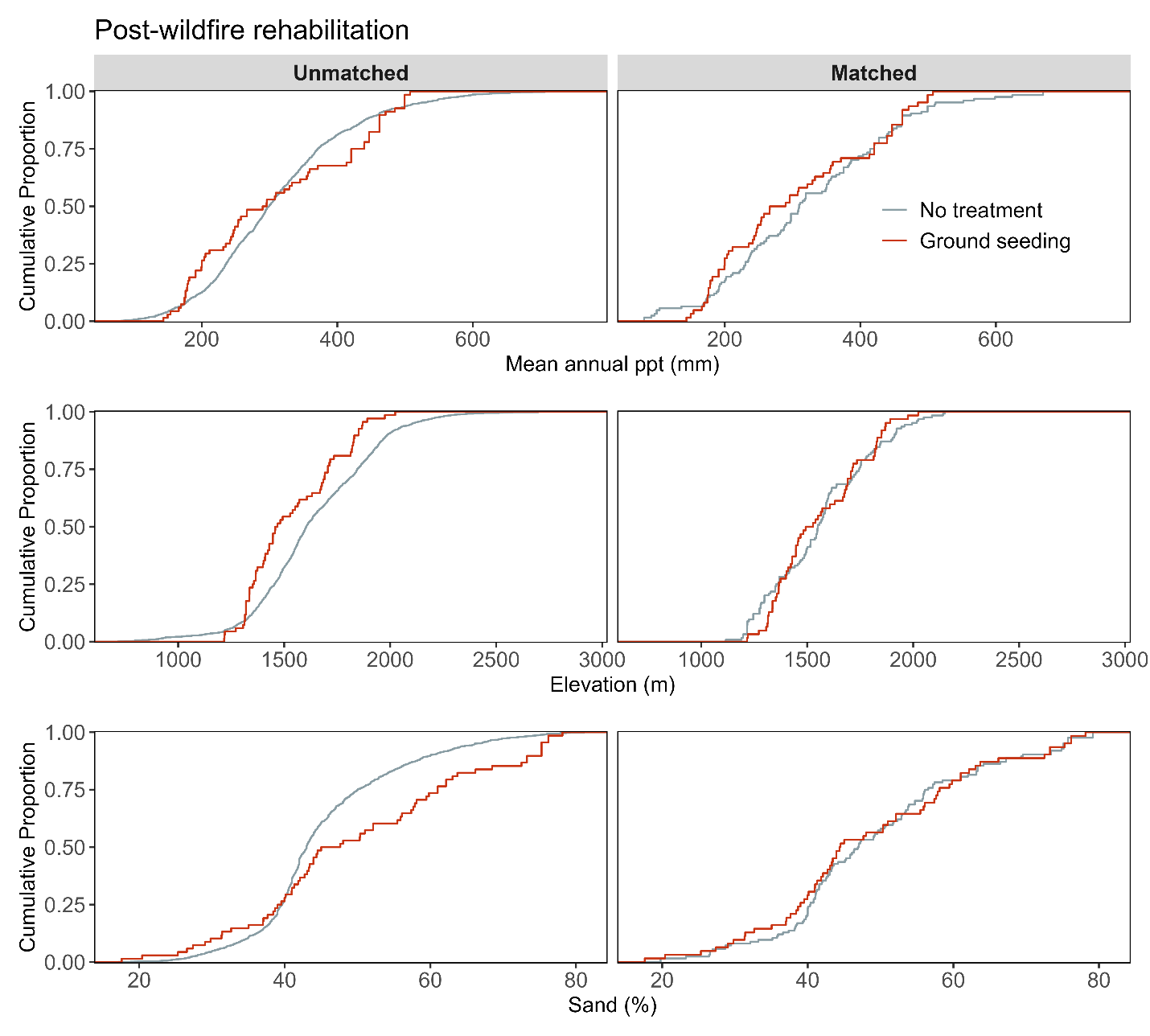

**Figure S14d.** Assessment of covariate balance using PSM for Post-fire rehabilitation monitoring plots. The figure compares eCDFs for mean annual precipitation (top), elevation (middle), and sand texture (bottom) percentages between ground seeding and non-treated controls, for both unmatched (left column) and matched sets of monitoring plots (right column).

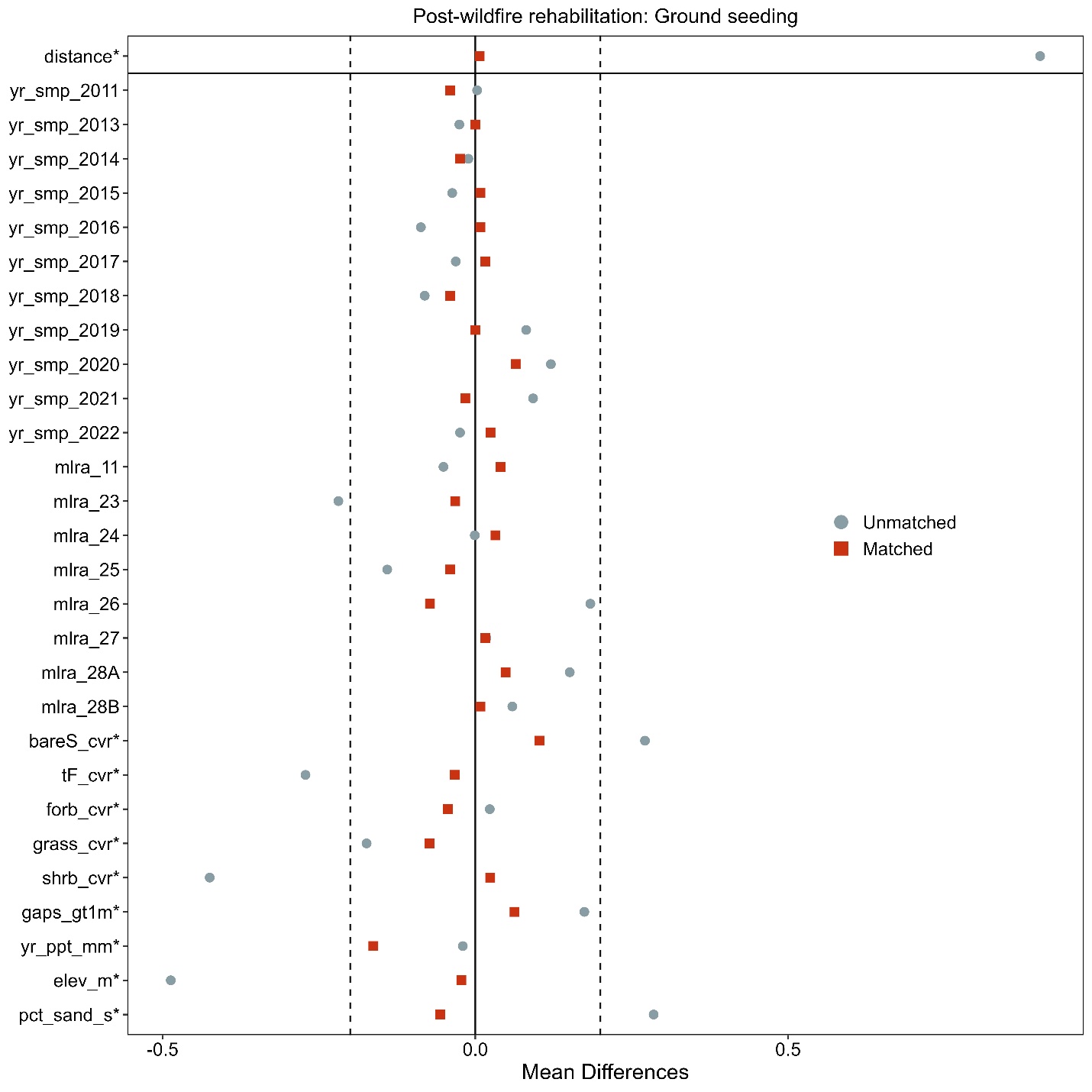

**Figure S14e**. Love-plot illustrating standardized mean differences for covariates in both the full dataset (unmatched) and the matched dataset for ground seeding and non-treated controls Post-fire rehabilitation plots. Vertical lines indicate the caliper threshold (0.2 times the standard deviation of the propensity score) applied during matching.

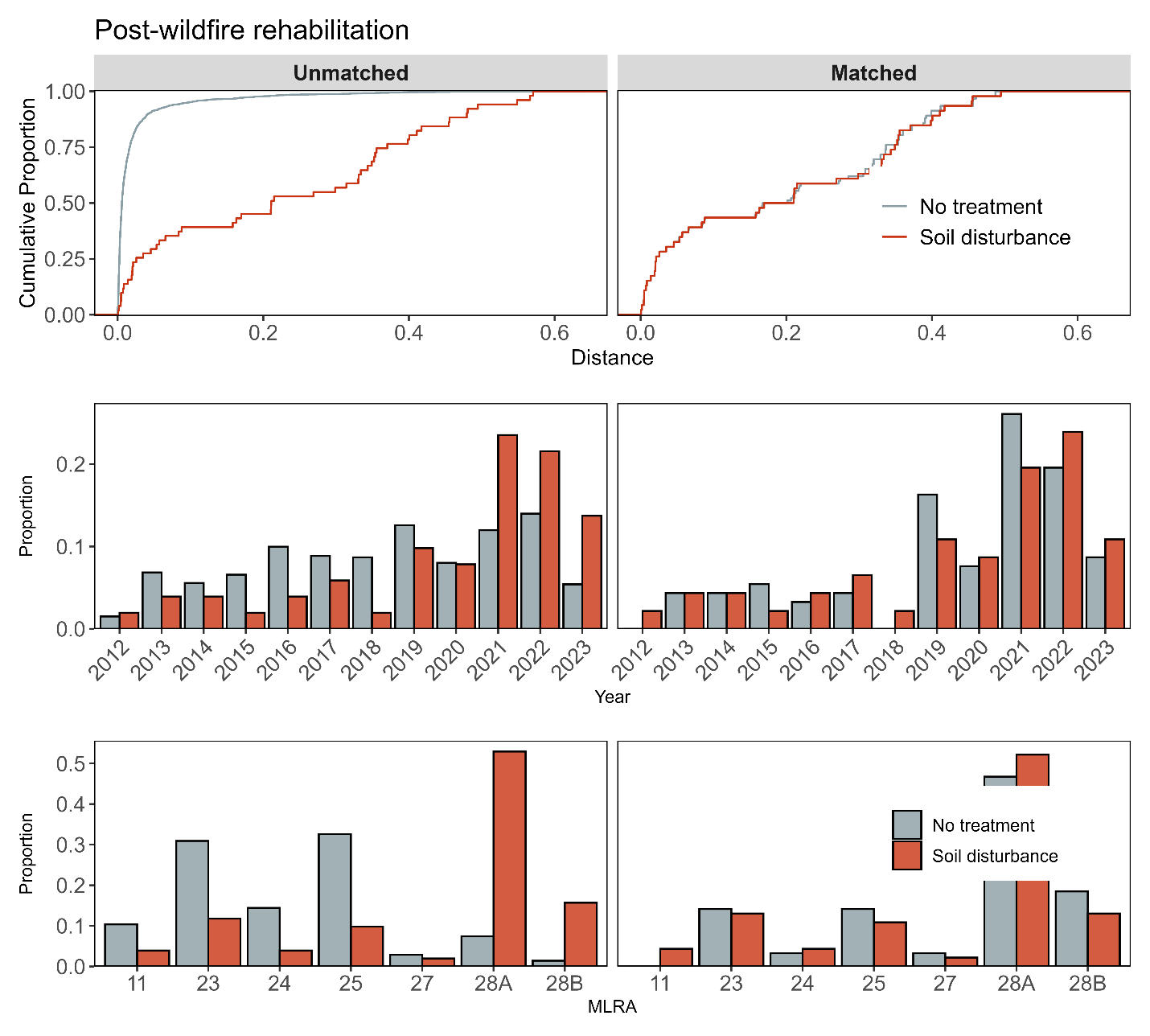

**Figure S15a**. Assessment of covariate balance using propensity score matching (PSM) for Post-fire rehabilitation monitoring plots. The top plot shows the empirical cumulative distribution function (eCDF) for the distance measure of the propensity score. The middle and bottom plots display balance in categorical variables, accounting for the year of monitoring and the major land resource area (MLRA) of the plots, comparing unmatched (left column) and matched sets (right columns) for soil disturbance and non-treated control monitoring plots.

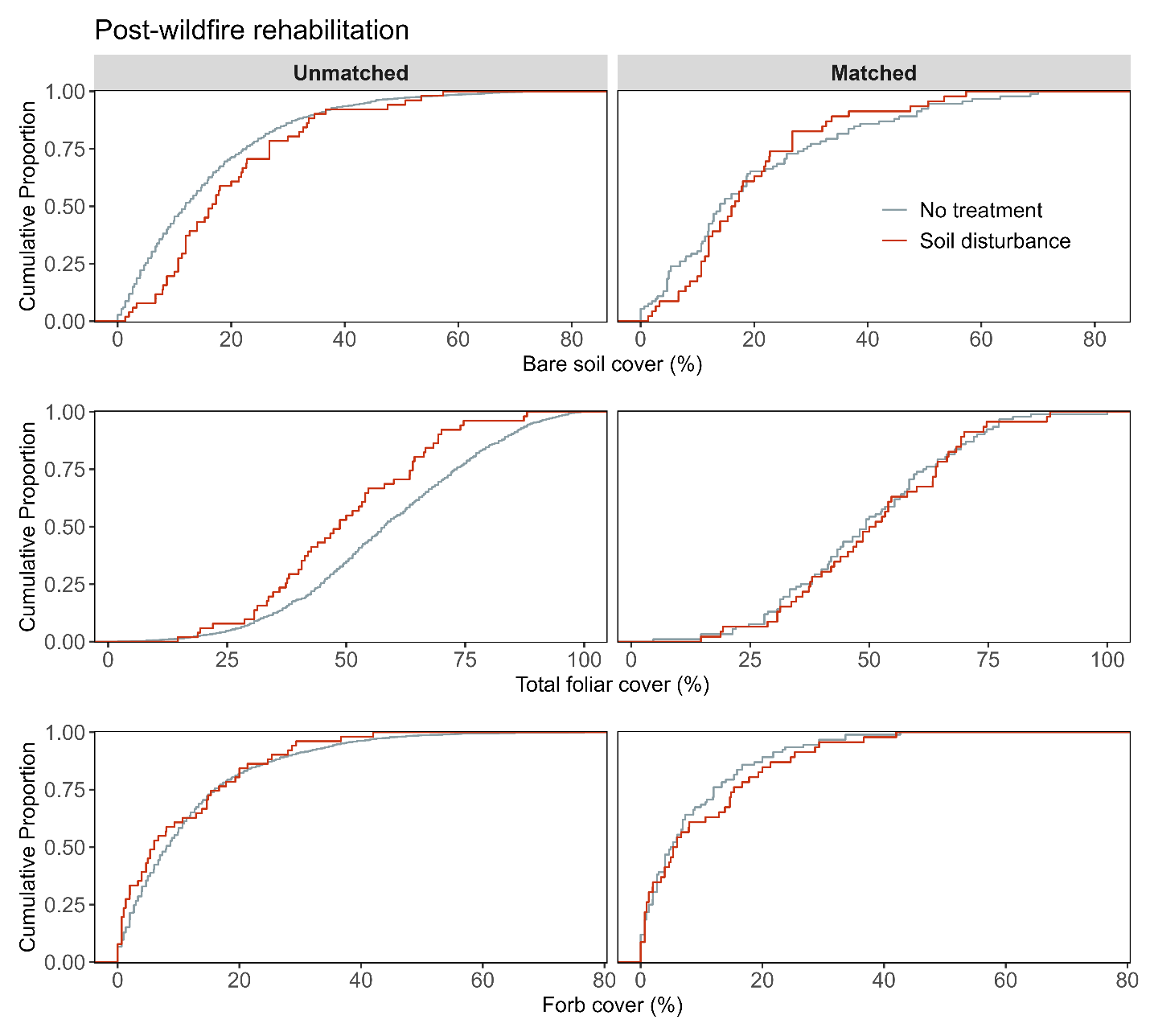

**Figure S15b**. Assessment of covariate balance using PSM for Post-fire rehabilitation monitoring plots. The figure compares eCDFs for bare soil (top), total foliar cover (middle), and forb cover (bottom) percentages between soil disturbance and non-treated controls, for both unmatched (left column) and matched sets of monitoring plots (right column).

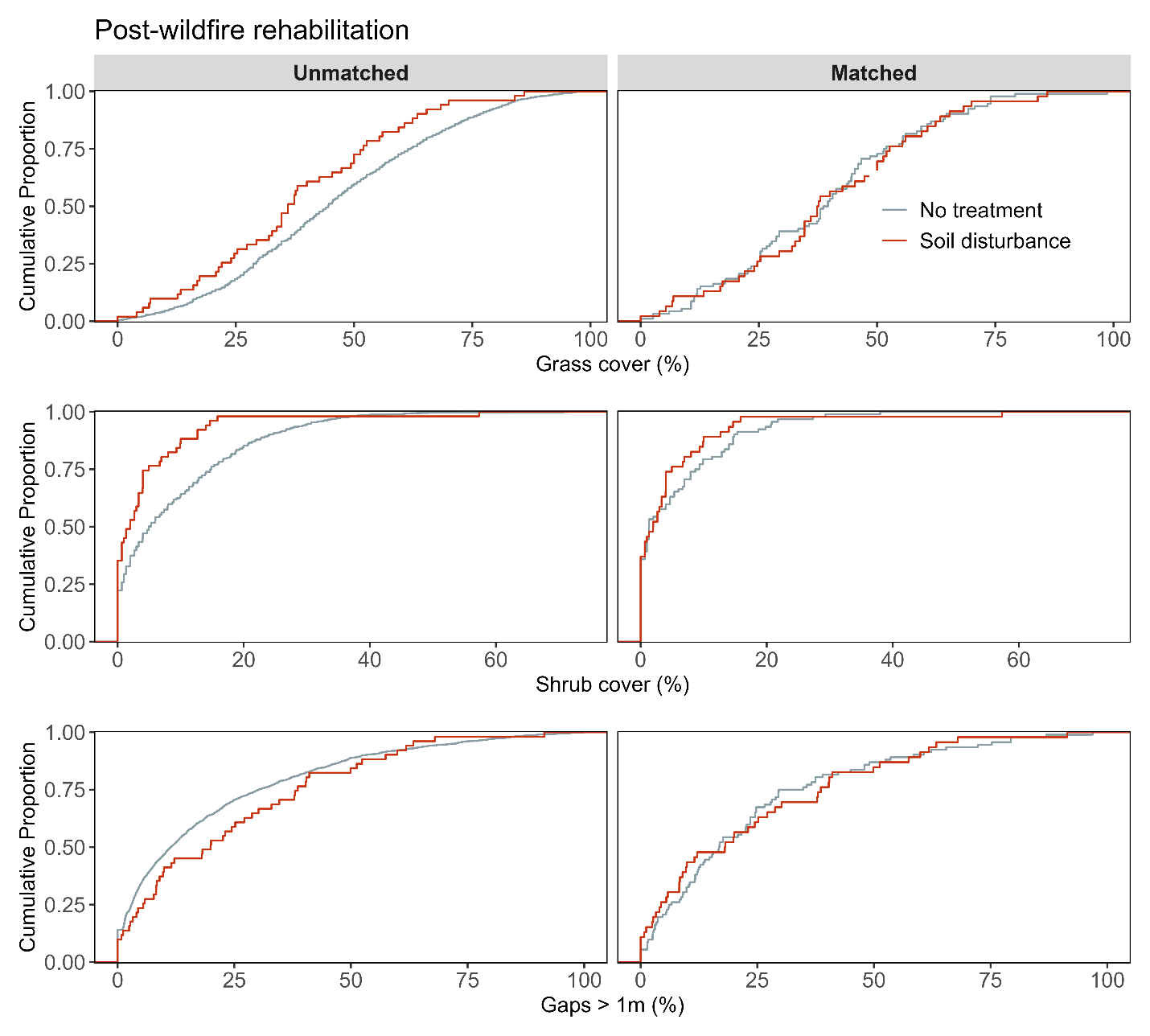

**Figure S15c**. Assessment of covariate balance using PSM for Post-fire rehabilitation monitoring plots. The figure compares eCDFs for grass cover (top), shrub cover (middle), and gaps > 1m (bottom) percentages between soil disturbance and non-treated controls, for both unmatched (left column) and matched sets of monitoring plots (right column).

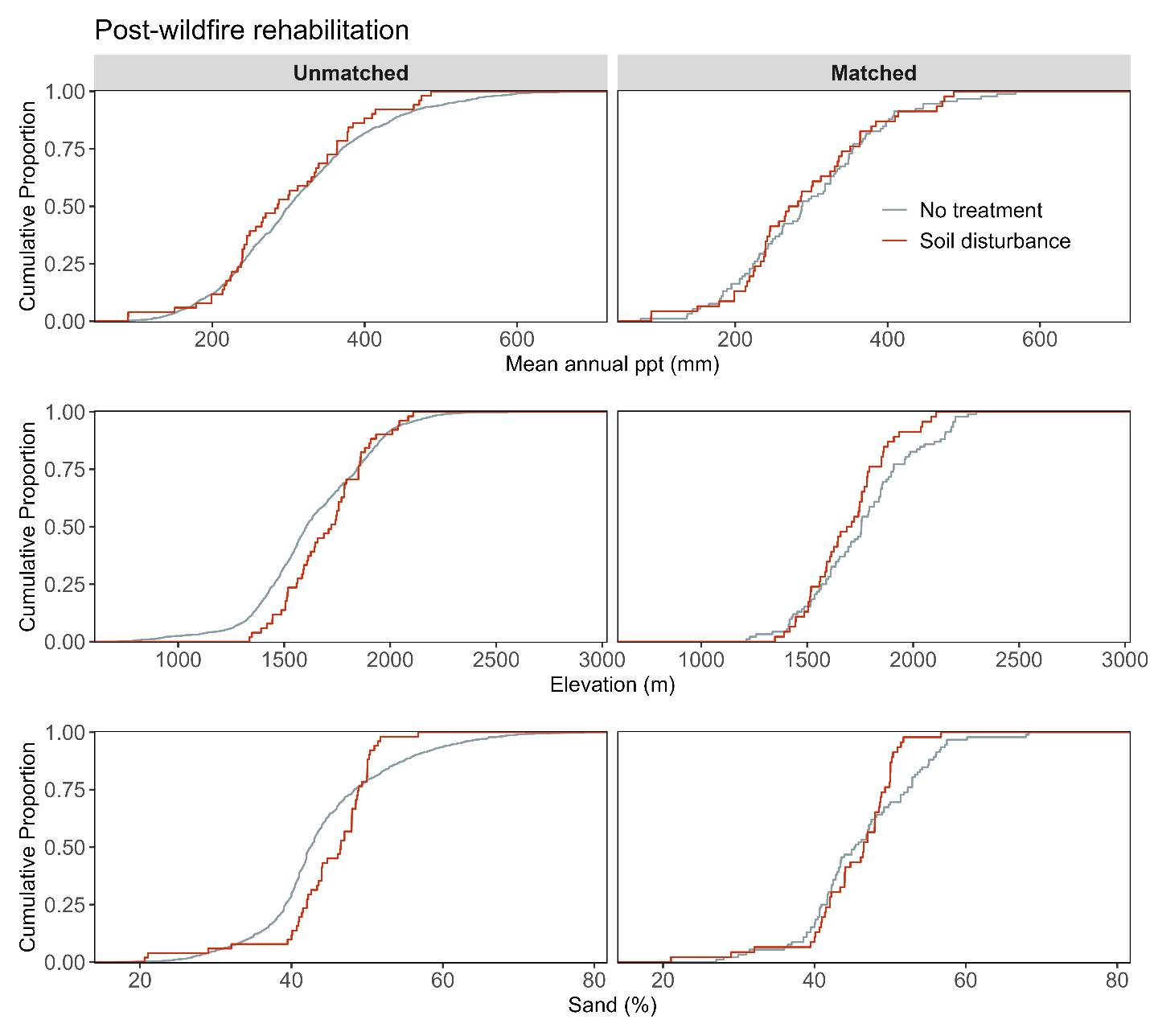

**Figure S15d.** Assessment of covariate balance using PSM for Post-fire rehabilitation monitoring plots. The figure compares eCDFs for mean annual precipitation (top), elevation (middle), and sand texture (bottom) percentages between soil disturbance and non-treated controls, for both unmatched (left column) and matched sets of monitoring plots (right column).

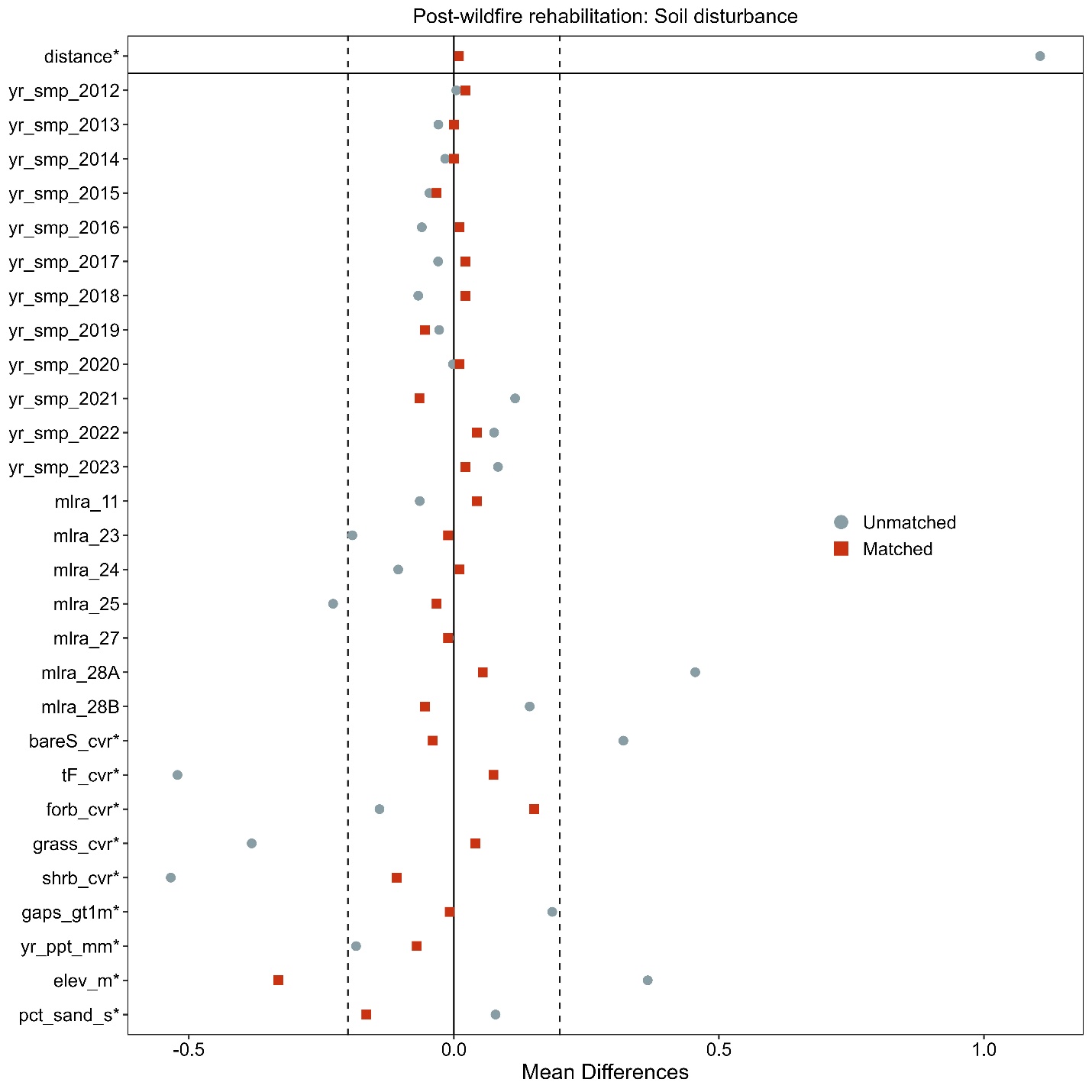

**Figure S15e**. Love-plot illustrating standardized mean differences for covariates in both the full dataset (unmatched) and the matched dataset for soil disturbance and non-treated controls Post-fire rehabilitation plots. Vertical lines indicate the caliper threshold (0.2 times the standard deviation of the propensity score) applied during matching.

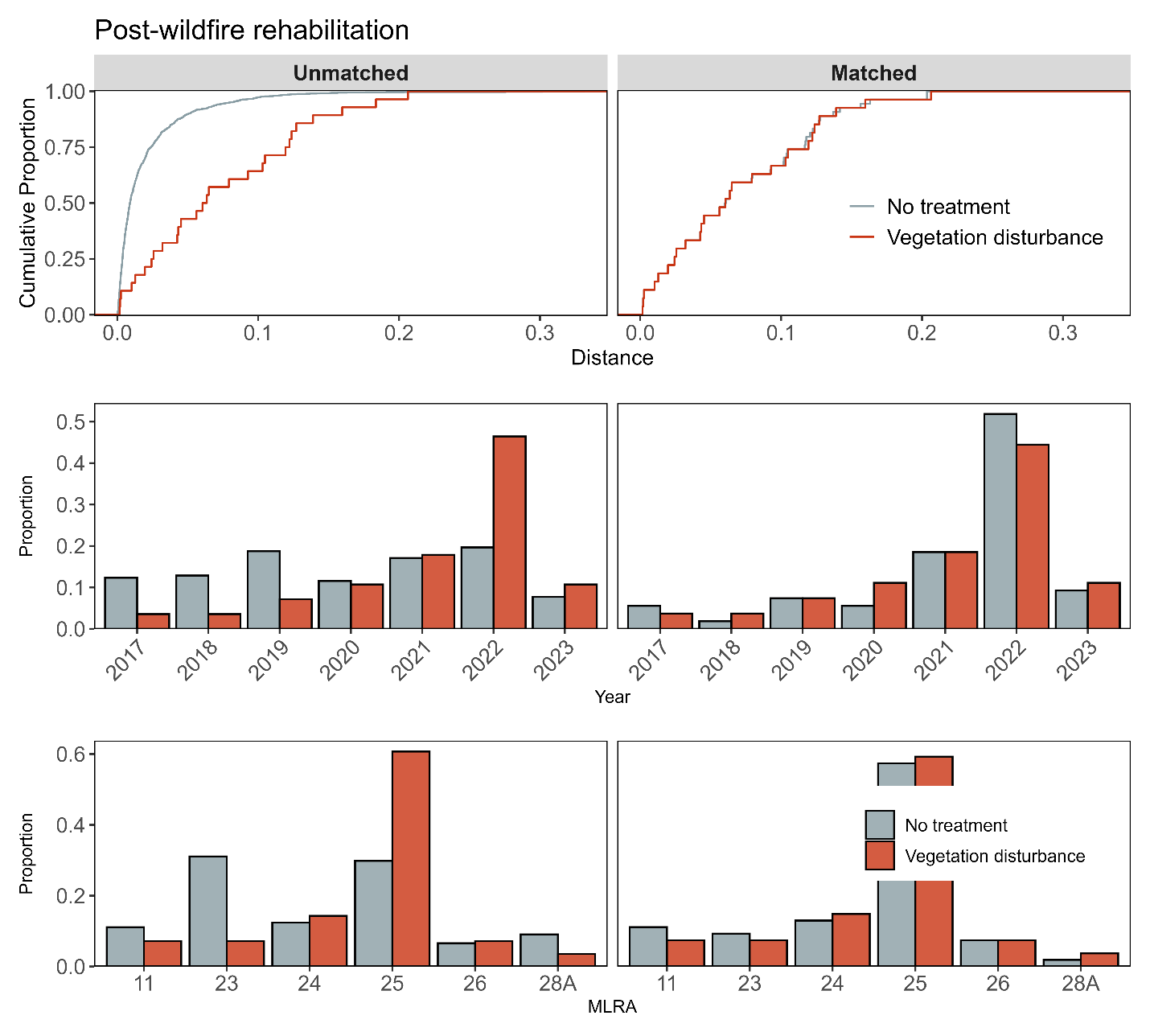

**Figure S16a**. Assessment of covariate balance using propensity score matching (PSM) for Post-fire rehabilitation monitoring plots. The top plot shows the empirical cumulative distribution function (eCDF) for the distance measure of the propensity score. The middle and bottom plots display balance in categorical variables, accounting for the year of monitoring and the major land resource area (MLRA) of the plots, comparing unmatched (left column) and matched sets (right columns) for vegetation disturbance and non-treated control monitoring plots.

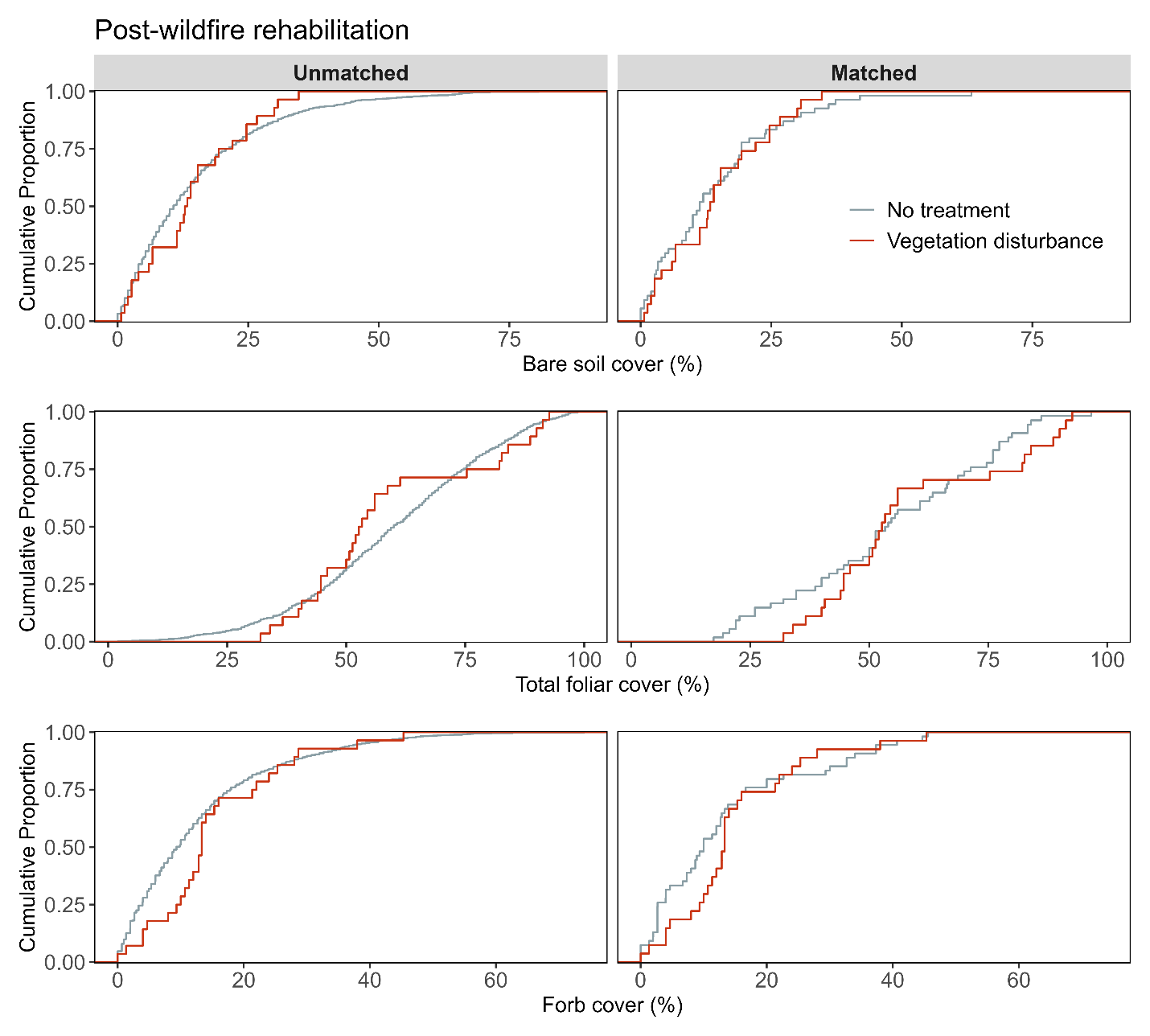

**Figure S16b**. Assessment of covariate balance using PSM for Post-fire rehabilitation monitoring plots. The figure compares eCDFs for bare soil (top), total foliar cover (middle), and forb cover (bottom) percentages between vegetation disturbance and non-treated controls, for both unmatched (left column) and matched sets of monitoring plots (right column).

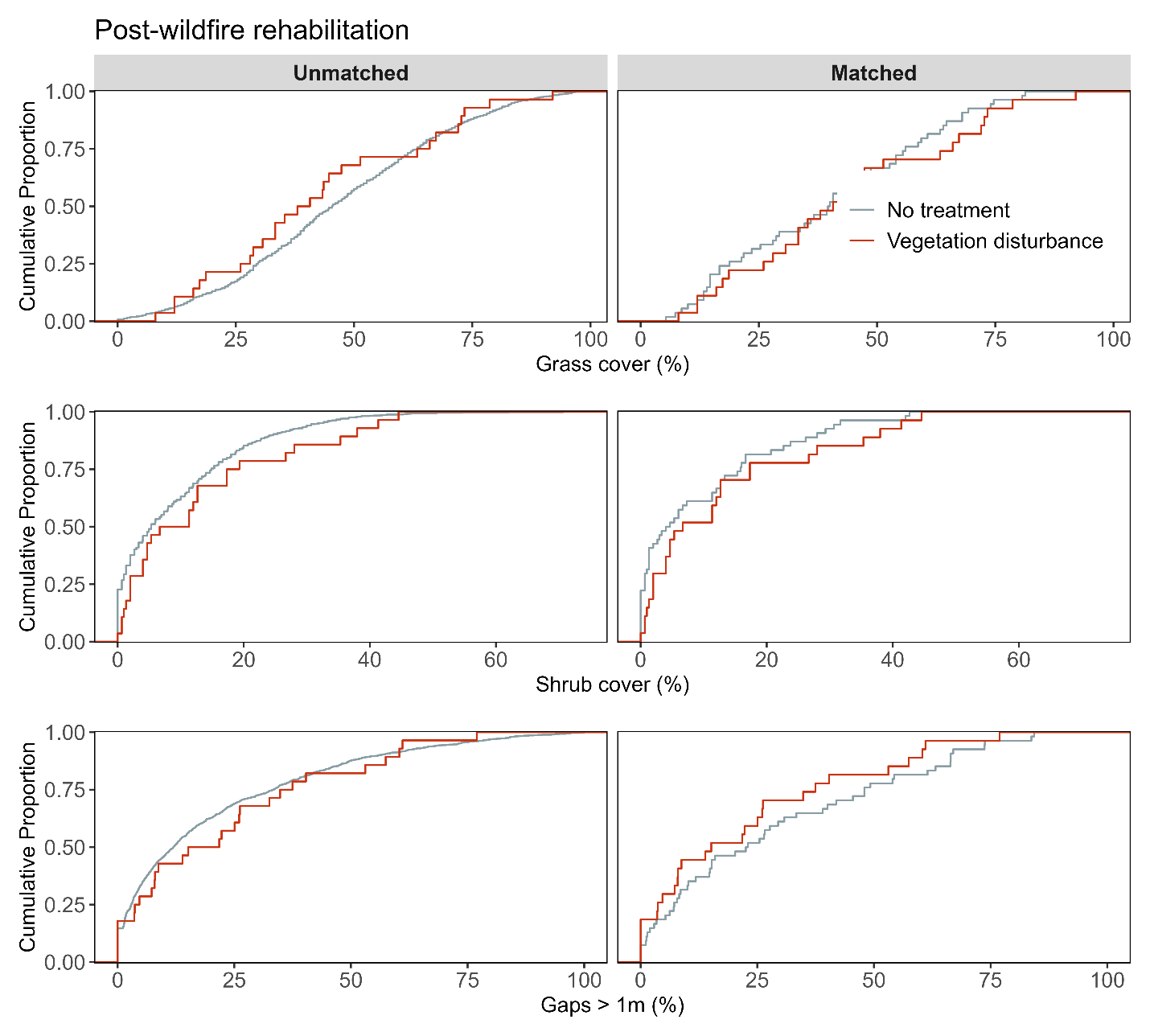

**Figure S16c**. Assessment of covariate balance using PSM for Post-fire rehabilitation monitoring plots. The figure compares eCDFs for grass cover (top), shrub cover (middle), and gaps > 1m (bottom) percentages between vegetation disturbance and non-treated controls, for both unmatched (left column) and matched sets of monitoring plots (right column).

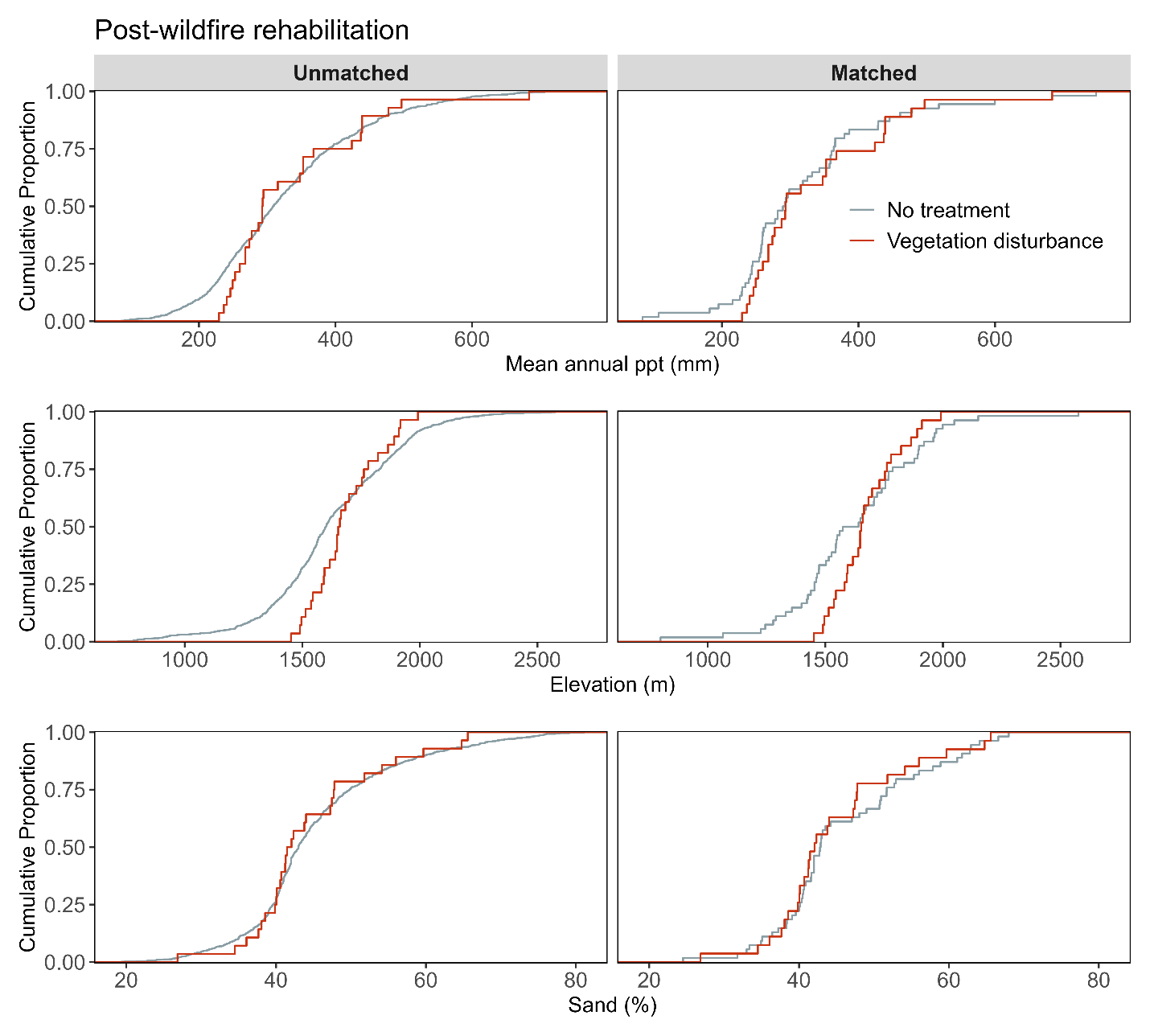

**Figure S16d.** Assessment of covariate balance using PSM for Post-fire rehabilitation monitoring plots. The figure compares eCDFs for mean annual precipitation (top), elevation (middle), and sand texture (bottom) percentages between vegetation disturbance and non-treated controls, for both unmatched (left column) and matched sets of monitoring plots (right column).

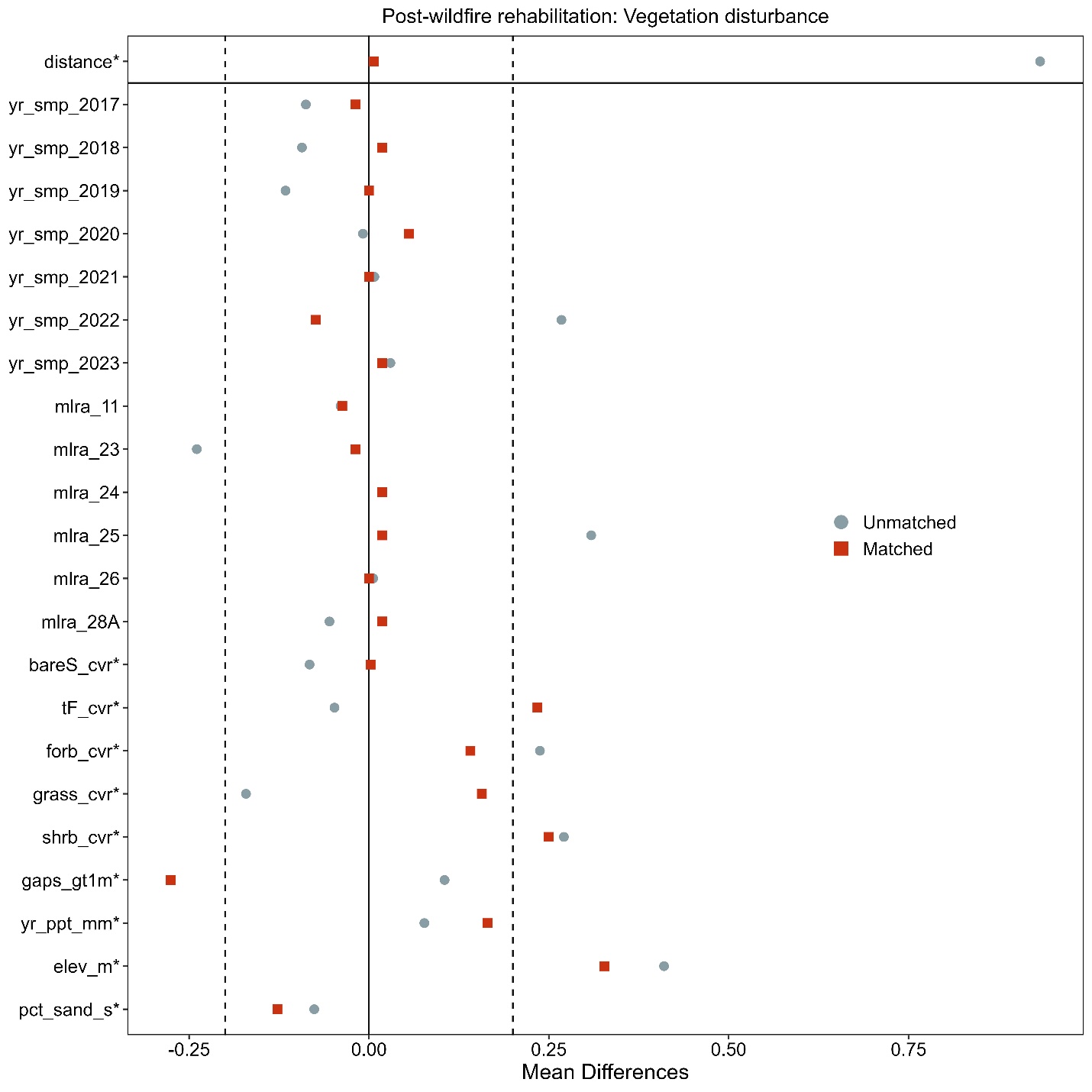

**Figure S16e**. Love-plot illustrating standardized mean differences for covariates in both the full dataset (unmatched) and the matched dataset for vegetation disturbance and non-treated controls Post-fire rehabilitation plots. Vertical lines indicate the caliper threshold (0.2 times the standard deviation of the propensity score) applied during matching.

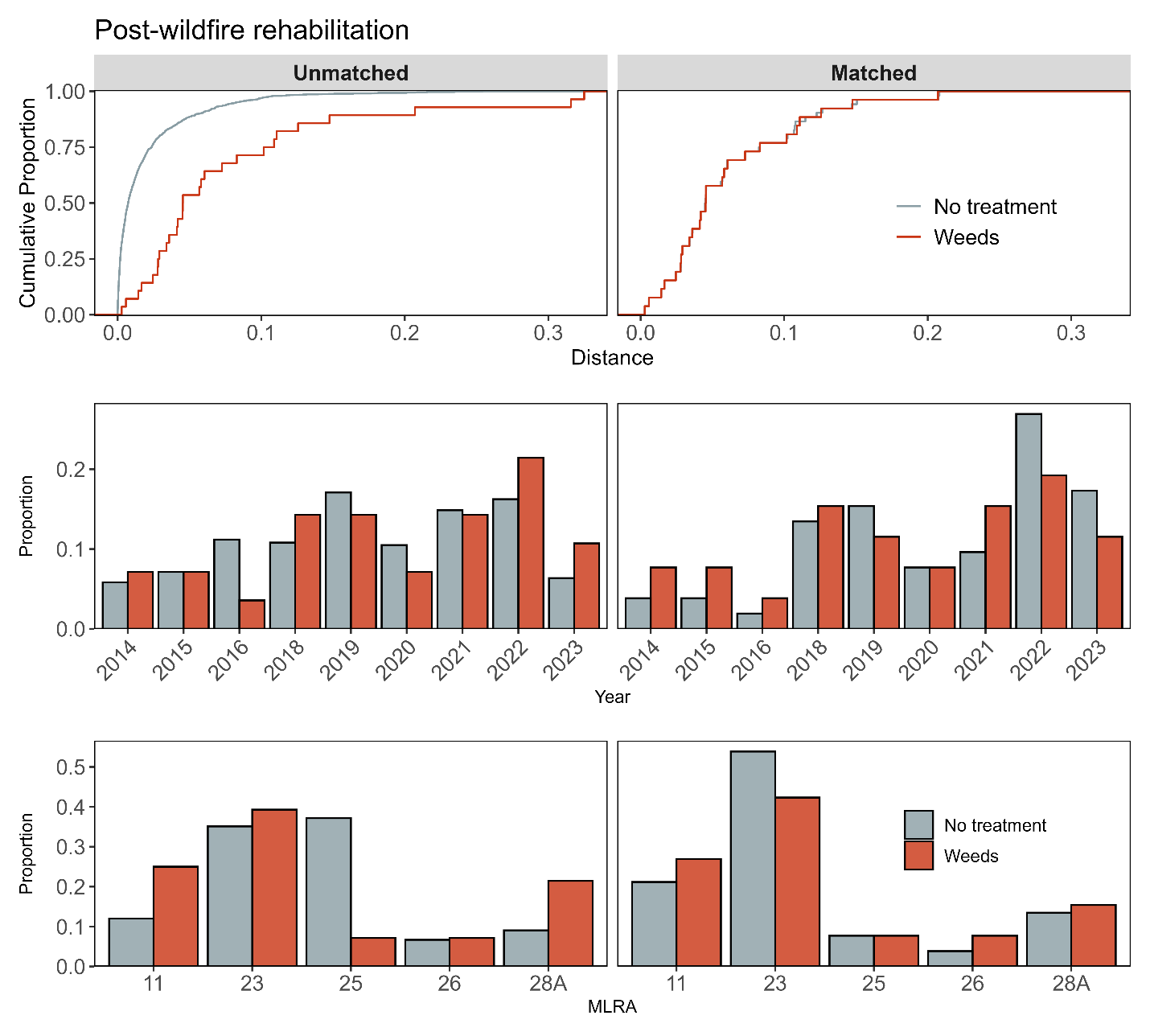

**Figure S17a**. Assessment of covariate balance using propensity score matching (PSM) for Post-fire rehabilitation monitoring plots. The top plot shows the empirical cumulative distribution function (eCDF) for the distance measure of the propensity score. The middle and bottom plots display balance in categorical variables, accounting for the year of monitoring and the major land resource area (MLRA) of the plots, comparing unmatched (left column) and matched sets (right columns) for weeds treatment and non-treated control monitoring plots.

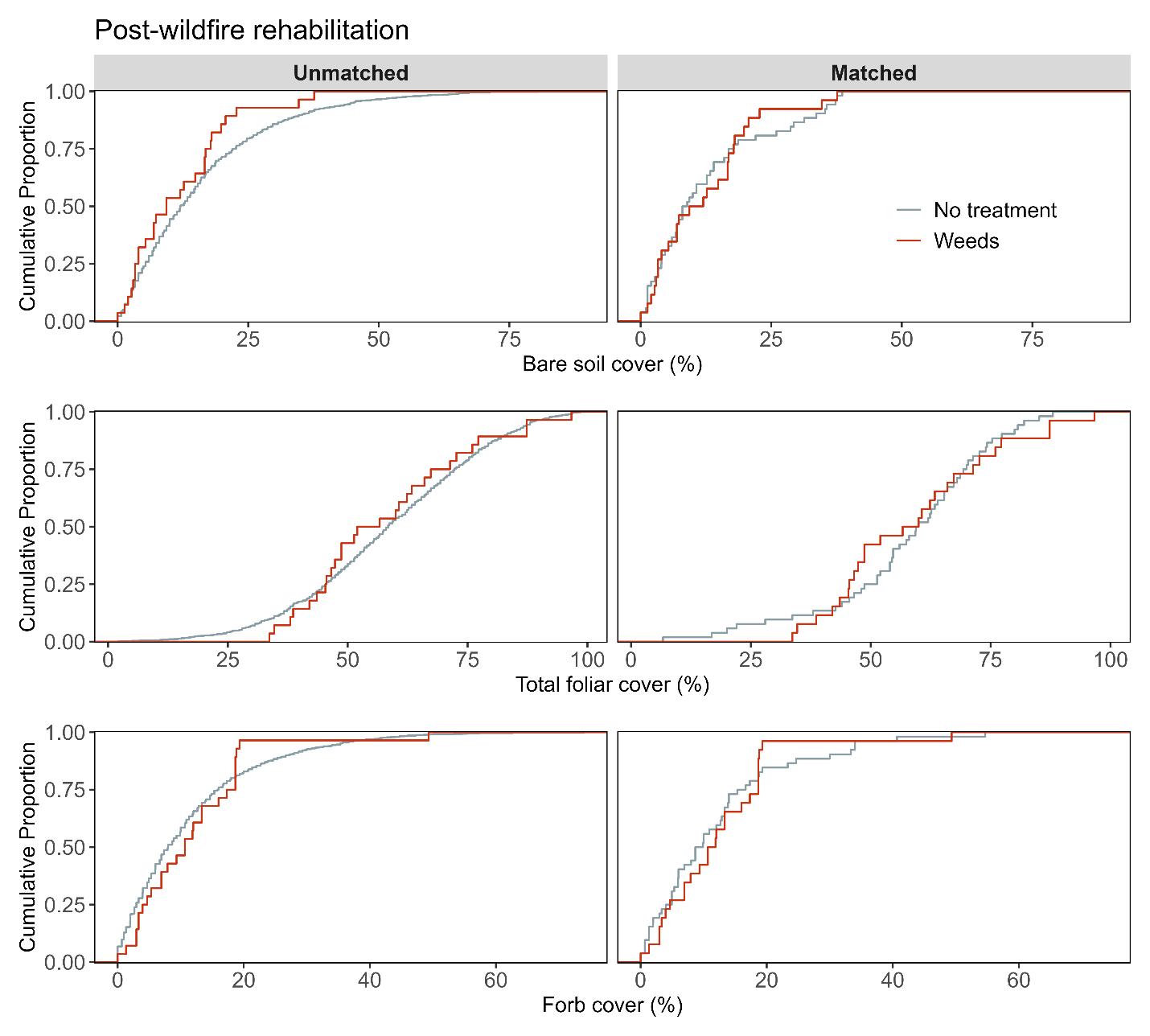

**Figure S17b**. Assessment of covariate balance using PSM for Post-fire rehabilitation monitoring plots. The figure compares eCDFs for bare soil (top), total foliar cover (middle), and forb cover (bottom) percentages between weeds treatment and non-treated controls, for both unmatched (left column) and matched sets of monitoring plots (right column).

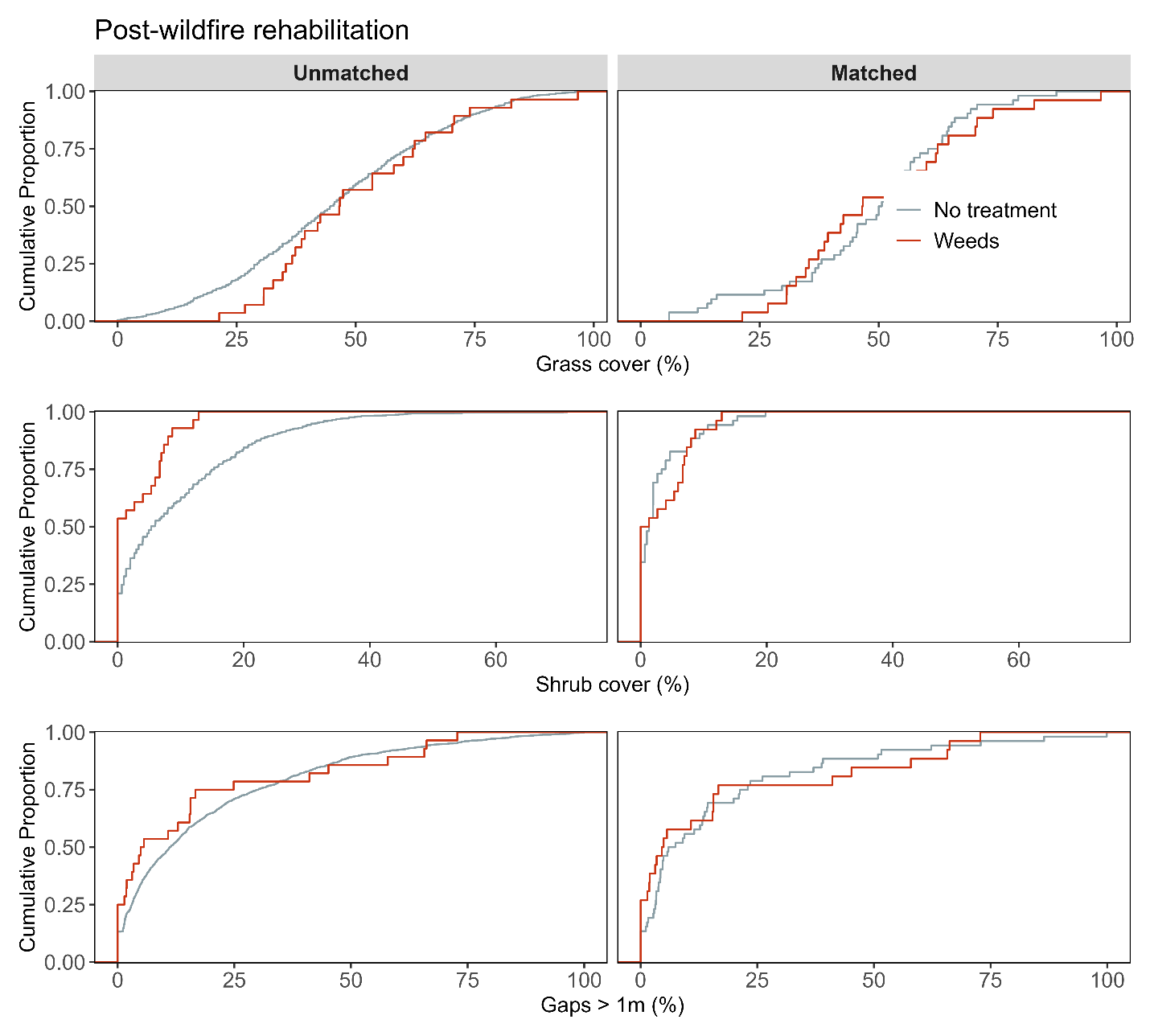

**Figure S17c**. Assessment of covariate balance using PSM for Post-fire rehabilitation monitoring plots. The figure compares eCDFs for grass cover (top), shrub cover (middle), and gaps > 1m (bottom) percentages between weeds treatment and non-treated controls, for both unmatched (left column) and matched sets of monitoring plots (right column).

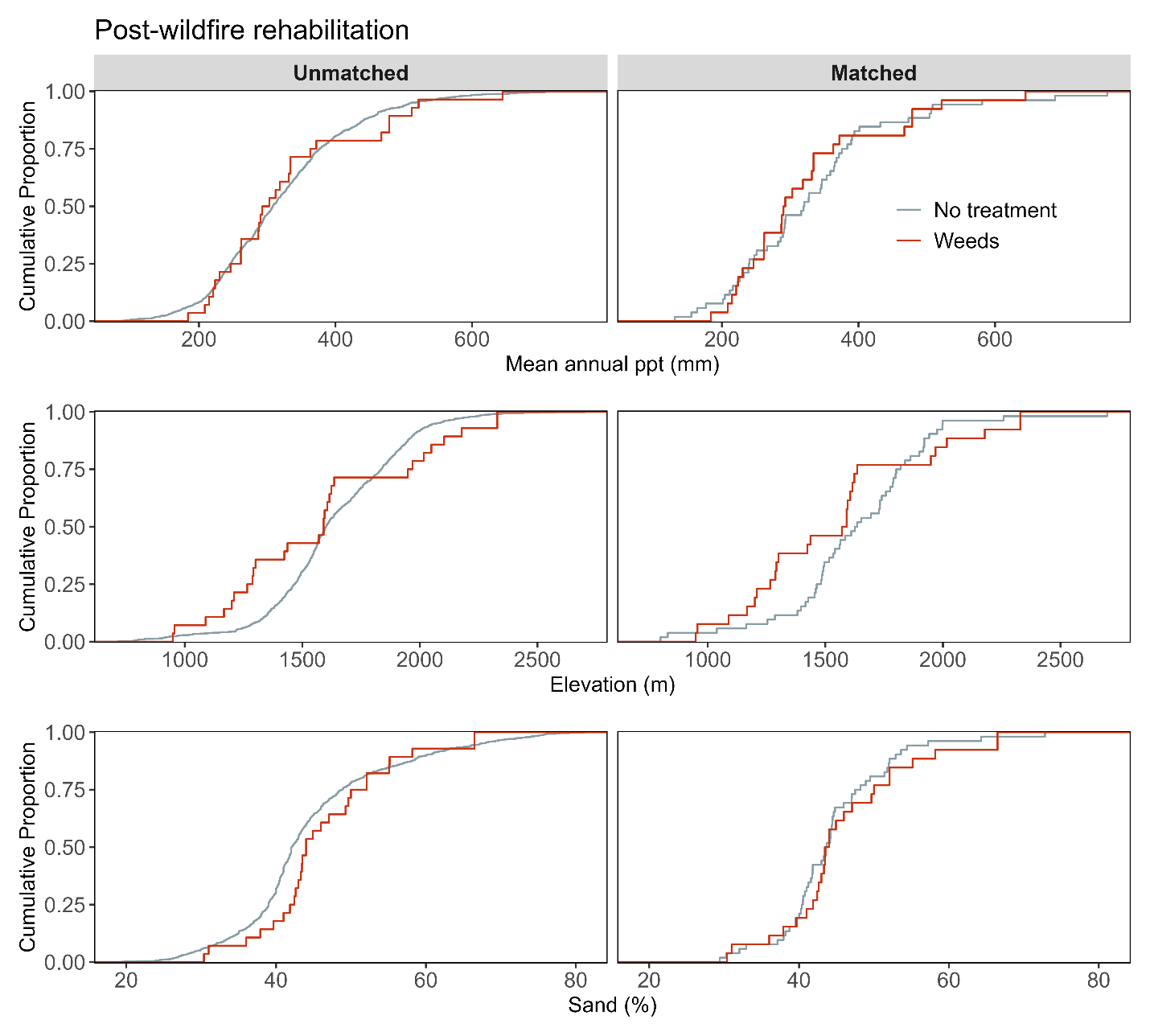

**Figure S17d.** Assessment of covariate balance using PSM for Post-fire rehabilitation monitoring plots. The figure compares eCDFs for mean annual precipitation (top), elevation (middle), and sand texture (bottom) percentages between weeds treatment and non-treated controls, for both unmatched (left column) and matched sets of monitoring plots (right column).

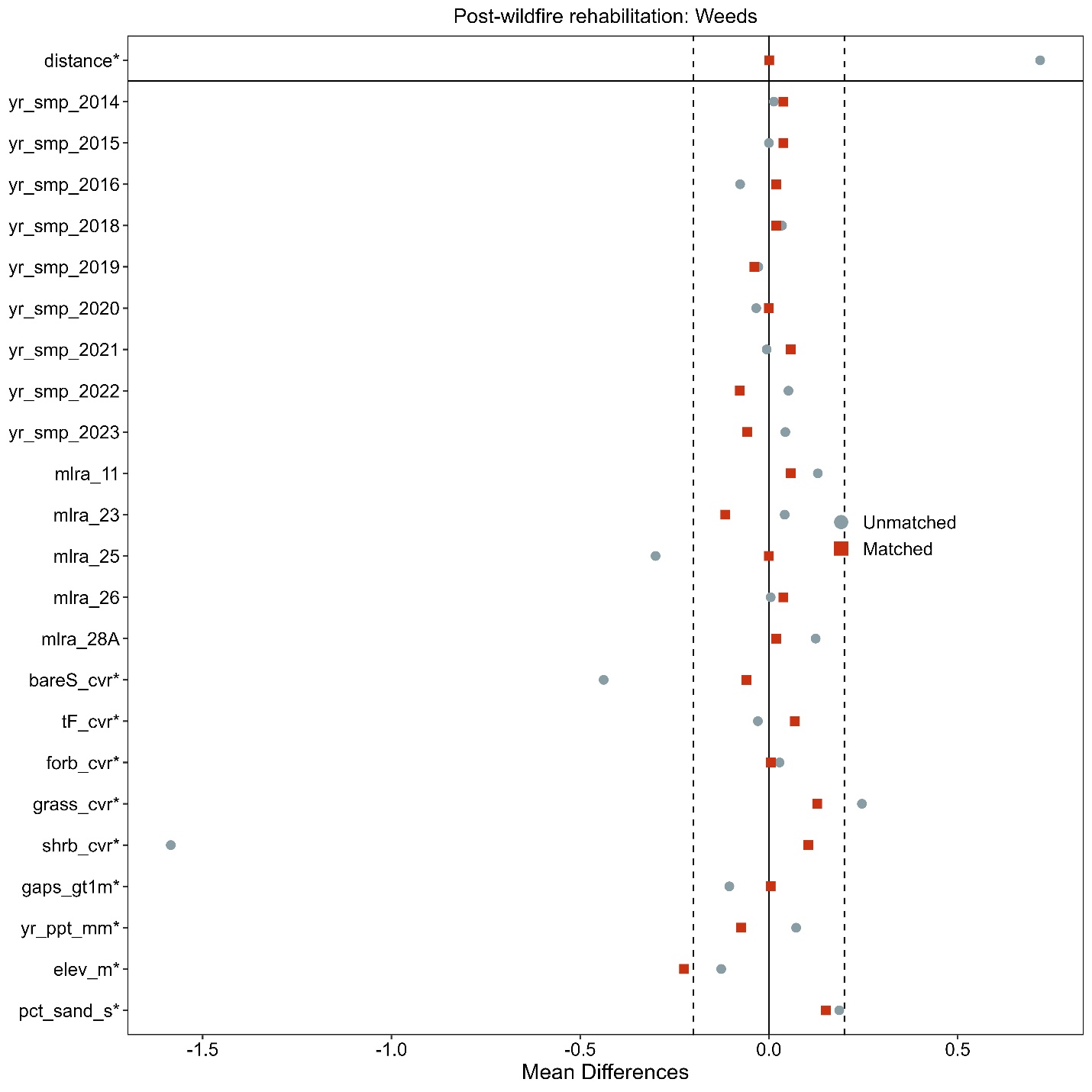

**Figure S17e**. Love-plot illustrating standardized mean differences for covariates in both the full dataset (unmatched) and the matched dataset for weeds treatment and non-treated controls Post-fire rehabilitation plots. Vertical lines indicate the caliper threshold (0.2 times the standard deviation of the propensity score) applied during matching.
